# Absolute Quantification of Fluorescent Protein Fusions by Proteomics

**DOI:** 10.1101/2025.09.19.677389

**Authors:** Anna Shevchenko, Archishman Ghosh, Andrea Schuhmann, Aliona Bogdanova, Henrik Thomas, Viditha Rao, Eric R. Geertsma, T-Y. Dora Tang, Andrej Shevchenko

## Abstract

Fusions with fluorescence proteins (FPs) play a pivotal role in experimental biology because of their sensitive and spatially precise visualization by spectroscopy. However, observed fluorescence is not always proportional to their molar abundance. Only a fraction of the fusion protein containing the mature fluorescence chromophore is detectable by spectroscopy and there is no generic method for estimating its molar abundance. We developed a fluorescence-independent mass-spectrometry based approach for accurate absolute sub-femtomole quantification of FP-fusions that also estimates the protein faction having fully matured chromophore. The method exploits an isotopically labelled 68 kDa recombinant protein standard containing peptide proxies for 6 prototypical FPs (mCherry, mScarlet-I, mKate2, EGFP, mNeonGreen and Dendra2) and two self-labelling (Halo- and SNAP-) tags. It enables quantification of proteins fused to any of more than 70 FPs or self-labelling tags. It is versatile, robust, precise and can be used broadly for the absolute quantification of fluorescent fusions in *in-vivo* and *in-vitro* cellular systems. As case study we combined mass spectrometry with fluorescence spectroscopy to study expression kinetics of FP fusions in cell-free systems. Molar concentrations of the expressed fusion protein, its fraction containing the mature fluorescent chromophore and of the fluorescent protein were integrated into a mathematical model to obtain kinetic rates of translation, chromophore maturation and folding.

## INTRODUCTION

**<u>F</u>**luorescent **<u>p</u>**roteins (FPs) revolutionized molecular and cell biology by providing a visual tool to track proteins in *in vitro* and *in vivo* models. Their discovery and development were recognized by the Nobel Prize awarded to O.Shimomura, M.Chalfie and R.Y. Tsien in 2008 (*1*). The FP family comprises several hundred natural and designed proteins (*2*) (*see* www.fpbase.com) each having distinctive photochemical (*e.g*. excitation and emission maxima or brightness) and physicochemical (*e.g*. lifetime, photostability or maturation time) characteristics (*3-6*) and the potential for engineering novel, application-specific FPs remains significant (*7*). To enhance optical detectability and imaging properties, a family of self-labelling protein tags (*e.g*. Halo, SNAP or CLIP) has been developed alongside prototypical FPs. While these tags themselves are not fluorescent, they covalently bind small fluorescent dyes (*8-10*), which exhibit superior brightness and photostability.

Fusions of FP to non-fluorescent proteins (further termed as “FP-fusions”) or to specific recognition domains are utilized in a variety of applications. FP-fusions serve as markers of protein localization, indicators of gene expression and biosensors for non-invasive, spatial and temporal imaging of biochemical processes and metabolites in living systems (*11-13*). They can be visualized in cells, organelles or even smaller sub-cellular structures by means of ultrahigh-resolution fluorescence microscopy with extraordinary specificity and spatial precision (reviewed in (*14, 15*)). However, the observed fluorescent signal provided by the genetically encoded FP-reporter is not always proportional to the molar abundance of FP-fusion (*16*). To become fluorescent, the chromophore in an FP sequence (with the exception of non-prototypical group of FPs (*4, 17-19*)) matures *via* cyclization, oxidation and dehydration of three successively positioned amino acid residues. This chromophore-forming triad comprising the amino acids sequence of …(M/Q/G/H)YG… in green and red FPs or another aromatic amino acid residue at the second position in blue FPs should yield a conjugated cyclic structure (*20-22*). Completing the chromophore maturation may take anywhere from minutes to hours (*23*). The observed fluorescence could be affected by the intrinsic properties of FP-fusion and by experiment-dependent factors including photobleaching, blinking, ligands binding or multimerization, to mention only a few. Hence, a significant (conveniently termed as “dark”) fraction of full-length FP-fusions may not be able to absorb or emit light because it lacks the correctly assembled chromophore. For self-labelling tags quantitative readout could be compromised by limited labelling efficiency as well as hindered delivery, unspecific binding or incomplete folding of ligands (*24*).

There is considerable interest in achieving robust and reproducible molar quantification of FP-fusions (*7, 14, 25-27*). Knowledge of the molar amount of FP-fusion enables to refer its abundance to the abundance of endogenous proteins and help to elucidate proteins interaction stoichiometry, kinetics of protein expression, folding and turnover *in vivo* and *in vitro*. Complementing fluorescent spectroscopy with absolute quantification of FP-fusions by mass spectrometry will ensure the analytical rigor necessary for a broad spectrum of biological applications.

Absolute (molar) quantification of proteins by mass spectrometry typically relies on the known amount of isotopically labelled peptide standards spiked into proteins digest prior LC-MS/MS analysis (reviewed in (*28-32*)). The peptide standards can be chemically synthetized or produced by in-situ enzymatic cleavage of metabolically labelled protein chimeras comprising concatenated peptide proxies for multiple target proteins (*33-38*). While effective, this approach is cumbersome and inflexible because for each target protein several unique peptides should be selected, synthetized and evaluated individually. Furthermore, when applied to FP-fusions, such analyses can only determine their total amount, while the proportion of the “dark” fraction, lacking a mature chromophore, remains unknown.

Here we demonstrate that absolute quantification of FP-fusions could be performed by mass spectrometry using a single generic chimeric protein standard containing peptide proxies of prototypical FPs and self-labelling tags. The same analysis could determine the molar fraction of FPs having the mature chromophore structure. We show that in cell free systems the combination of quantitative mass spectrometry and fluorescence spectroscopy reveals the kinetics of expression of FP fusions and maturation of their fluorescent chromophore, which could be described by generic sequential reaction model.

## RESULTS AND DISCUSSION

### Design of chimeric protein standard and quantification workflow

Within a protein fusion, FP or self-labelling protein tag is genetically encoded to the protein of interest in a known (typically, equimolar) stoichiometric ratio. We reasoned that peptides originating from the FP part of the fusion could proxy the molar quantification of the entire protein construct. Conceivably, it should be possible to quantify any FP-fusion using a single set of proxy peptides representing frequently used FPs.

To select the peptide proxies we first analysed tryptic digests of six prototypical FPs: the constitutively fluorescent red mCherry and mScarlet-I; the far-red mKate2; the green mEGFP, the yellow-green mNeonGreen, and the photoconvertible green-to-red Dendra2 (see Supplementary Table S1 and FPbase resource (*2*) for details), and the self-labelling Halo- and SNAP-Tag proteins (*8, 9*) - for brevity, here we will collectively term them as FPs. In LC-MS/MS chromatograms we picked abundant consistently detectable fully tryptic peptides whose sequences (preferably, but not necessarily) contained no Cys or Met, and no N-terminal Glu or Asp residues (*39*) that served as quantotypic peptide candidates (reviewed in (*40*)). To leverage incomplete tryptic cleavage, we further extended the sequences of some candidate peptides by 2 to 3 amino acid residues to mimic corresponding sequence stretches in the source FPs (*41*). Taken together, we selected 31 quantotypic peptide candidates consisting of 7 to 26 amino acid residues such that each of the eight FPs was represented by 3 to 5 unique sequences. Next, we produced a metabolically labelled chimeric protein termed qFP-8 (for <u>**q**</u>uantification of <u>**8**</u> <u>**FPs**</u>) that comprised these 31 peptides together with 6 peptides from the reference protein (BSA) and 4 peptides from an alternative reference protein (PhosB) flanked by sequences of Strep- and His-tags; see ref.(*35*) for details on its design and expression in the *Δarg1Δlys1* strain of *E.coli*, metabolic labelling with ^13^C_6_ ^15^N_4_ -Arg and ^13^C_6_-Lys, and quality control by LC-MS/MS (**Figure 1A**; Supplementary Fig S1). It was designed as generic internal standard that enables absolute quantification of any protein fused to any of over 70 individual FPs sharing quantotypic peptides with the qFP-8 sequence (Supplementary Table S1). No protein-specific standards are required and proteins fused to different FPs could be quantified in parallel.

**Figure 1.**
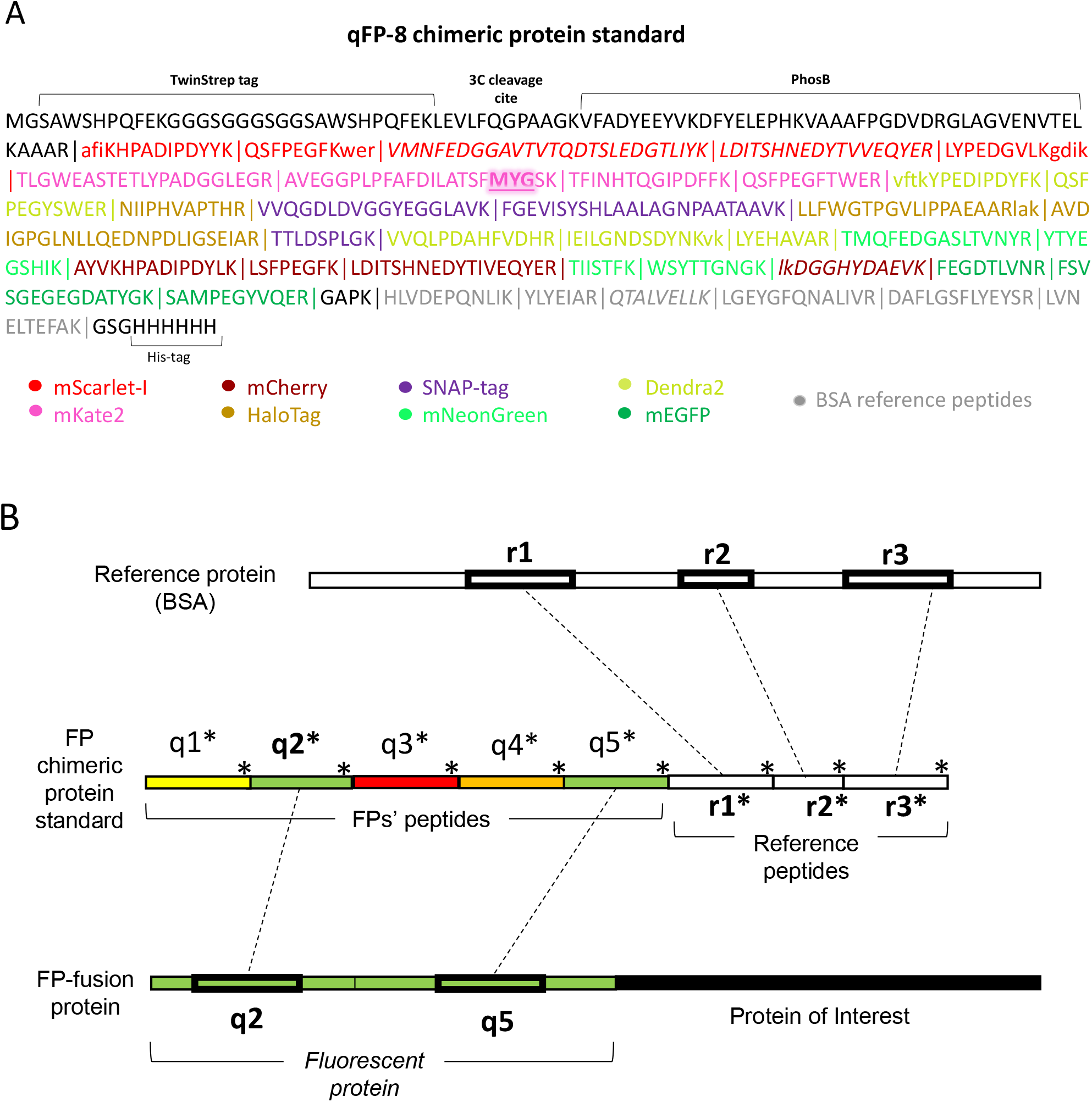
Absolute quantification of FP-fusions using qFP-8 standard. **A:** Sequence and modular composition of qFP-8. Peptide proxies from FPs and BSA reference peptides are colour coded; tryptic peptides are designated by vertical lines. Sequences extending some quantotypic peptides are in lower case. Peptides in italic did not pass validation check and were not used for quantification. The chromophore tripeptide […MYG…] within one of the mKate2 peptides is shadowed. **B:** Quantification workflow: q1..q5 represent (in this example, five) FP peptide proxies, colours indicate proxies of different FPs; r1…r3 are reference peptides identical to peptide from BSA. Asterisks indicate qFP-8 peptides metabolically labelled with ^13^C_6_ ^15^N_4_ -Arg or ^13^C_6_ -Lys. To exemplify the workflow, here green FP is fused to the N-terminus of a protein of interest

To quantify FP-fusions, we adapted the workflow based on the MS Western and FastCAT protocols (*35, 38*) (**Figure 1B** and Supplementary Figure S2). Briefly, FP-fusions, metabolically labelled qFP-8 and a known amount of reference protein (BSA) were co-digested with trypsin and the digest was analysed by LC-MS/MS. By comparing areas of XIC peaks of unlabelled reference peptides (from BSA) and corresponding labelled peptides (from qFP-8), we determined the exact amount of qFP-8 and, consequently, of all quantotypic peptides encoded within the qFP-8 sequence. Then pairs of labelled (from qFP8) and unlabelled (from FP fusion) peptides served as independent single-point calibrants for quantifying the molar abundance of the FP moiety and, in turn, the entire FP-fusion.

### Validation of absolute quantification of FP-fusions

Quantification of FP-fusion is based on the extensively validated MS Western protocol (*35*). However, both MS Western and its more advanced variant FastCAT (*38*) rely on quantotypic peptides picked by LC-MS/MS analyses of tryptic digests of full-length target proteins. We wondered if FP peptide proxies can accurately reflect the abundance of the full-length FP-fusion that could contain peptides with superior ionization properties. Since no FP-fusion standards with exactly known concentration and guaranteed purity are commercially available, we validated the quantification protocol in two ways. First, we tested if the yield of FP peptide proxies in the digests of FP-fusions and of qFP-8 standard was the same. Second, we checked if the targeted quantification using qFP-8 peptides corroborates independent LC-MS/MS determinations based on other (and, potentially, more abundant) peptides originating from the entire sequence of FP-fusions.

To address the first question, we co-digested each of nine FP-fusions containing different FPs (coded #A to #I in Supplementary Table S1 and Supplementary Dataset S1) with the isotopically labelled qFP-8 standard. Then, for each fusion protein, we compared normalized ratios of intensities of XIC peaks of quantotypic peptides from its FP tag and from the qFP-8 standard (*35, 38*). Excellent agreement between the ratios of 27 out of the total of 30 native (from FP-fusions) and isotopically labelled (from qFP-8) peptide pairs suggested complete digestion and unbiased peptide recovery, regardless of the FP tag location in the full-length FP-fusion sequence (**Figure 2A**, Supplementary Table S2, Supplementary Dataset S1).

**Figure 2.**
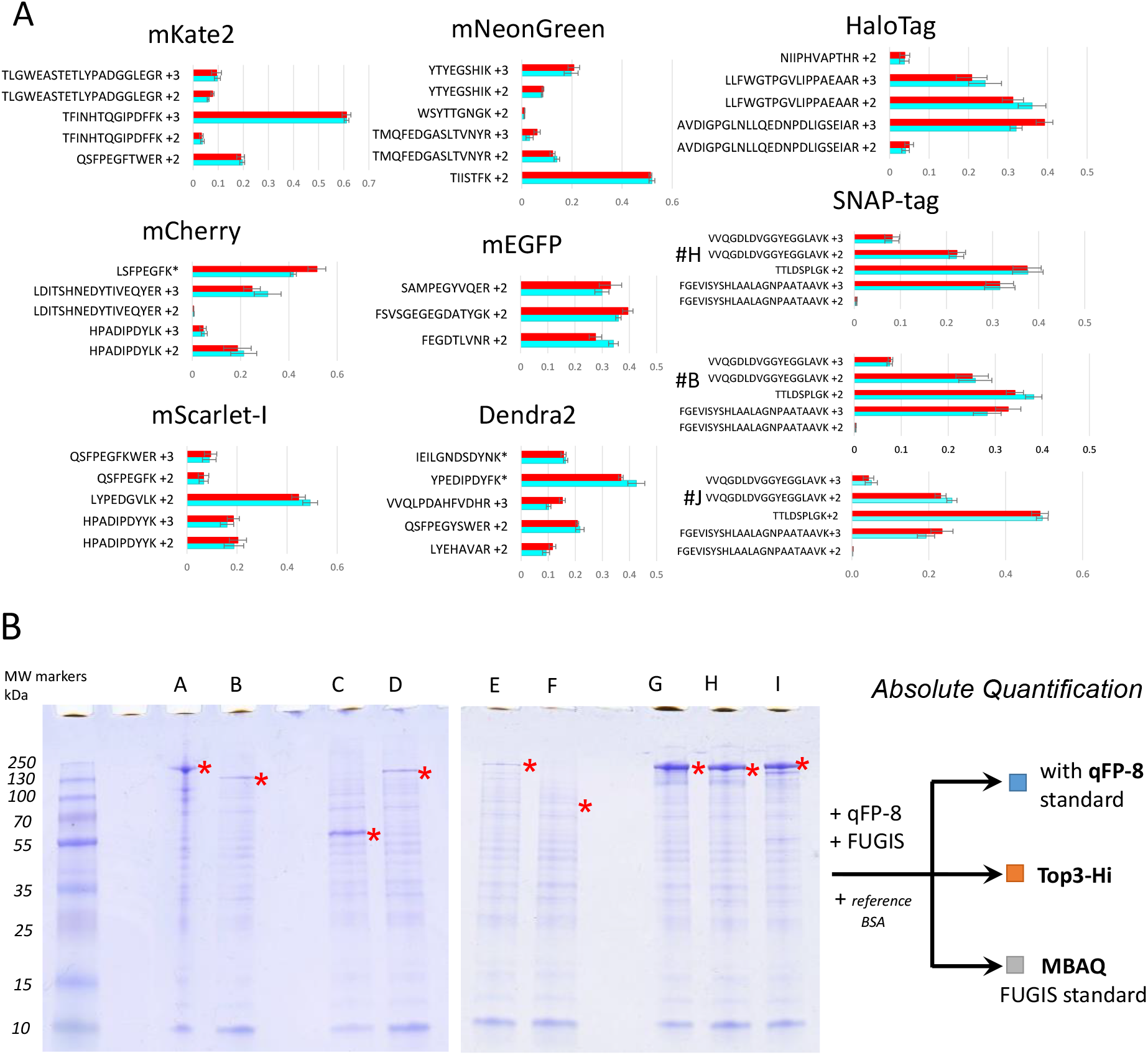
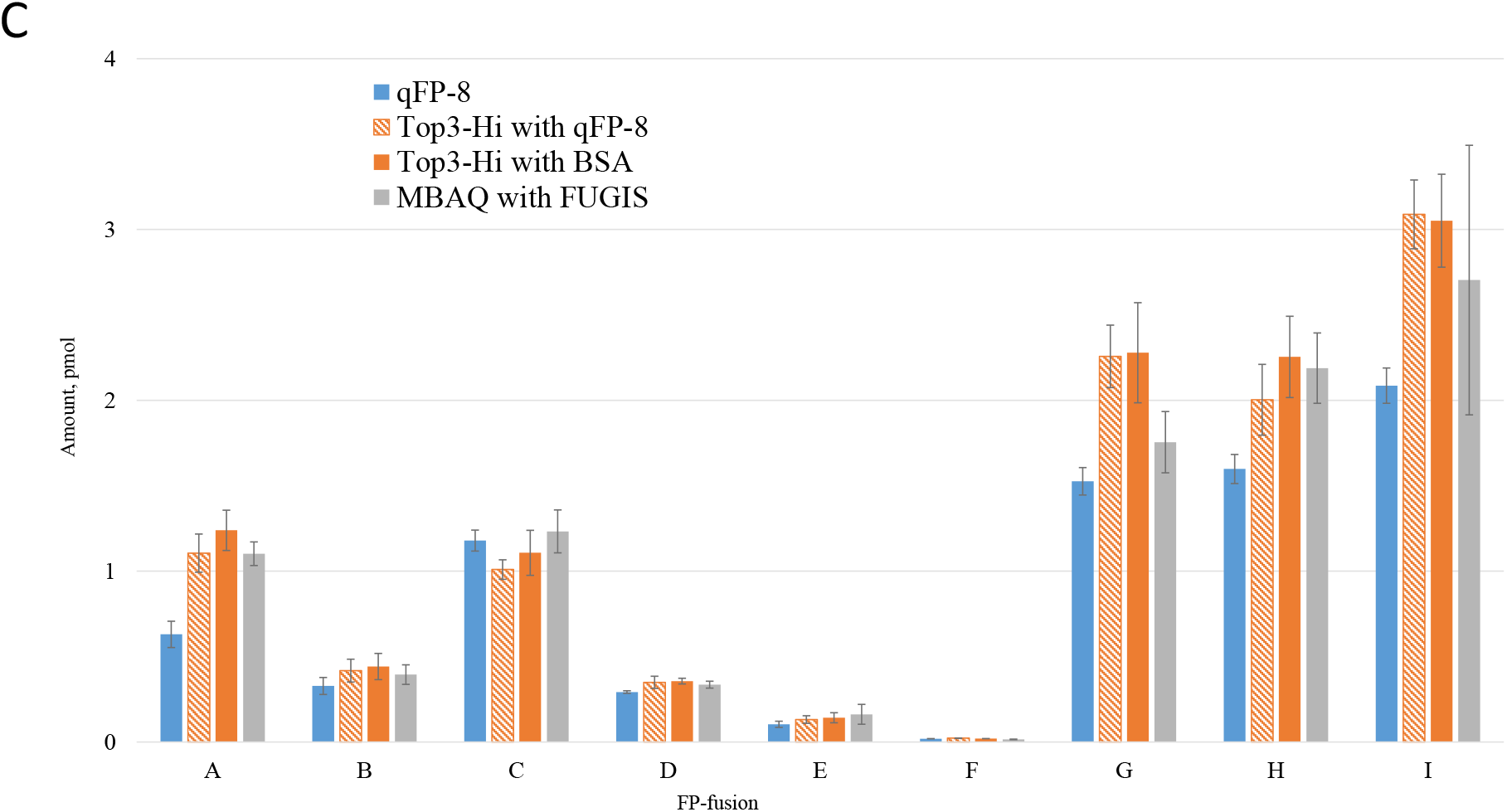
Validation of absolute quantification of FP-fusions using qFP-8 standard. **A:** Ratios of normalized abundances (*x*-axis) of quantotypic peptides (*y*-axes) originating from qFP-8 (in red) and from corresponding FP-fusions (in turquoise). Different charge states of the same peptide are shown as separate bars. Asterisks (*) designate tryptic peptides with the added abundances of mis-cleaved forms. #B, #H and #J stand for FP-fusions with Snap-tag located at the C-terminus, in the middle and the N-terminus of the full-length sequence, respectively (Supplementary Table S2). **B:** FP-fusions #A to #I were analysed by three methods: targeted quantification with qFP-8 and untargeted quantification Top3-Hi and MBAQ / FUGIS. Bands corresponding to full-length FP-fusions are designated with an asterisk (*); bands of MW markers are at the left-hand side. **C:** Molar amounts of FP-fusions per gel band determined by qFP-8 standard (blue bars); by Top3-Hi method using qFP-8 and by BSA as references (orange); by MBAQ / FUGIS (grey bars). Error bars are SD (*n=4*).

Next, we demonstrated that the abundance of quantotypic peptides from the FP tag truly reflected the abundance of the full-length FP-fusion. To this end, each of nine FP-fusions was co-digested with qFP-8 and with generic protein standard FUGIS designed for rapid proteome-wide absolute quantification (*42*), and analyzed the digests by LC-MS/MS. The molar amount of each FP-fusion was independently determined in three ways (**Figure 2B**, Supplementary Table S3): *a*) by targeted quantification using qFP-8 as above (**Figure 1B**, Supplementary Figure S2); *b*) by “best 3” peptides from the fusion referenced to peptides from FUGIS standard (MBAQ method, see ref. (*42*)); and *c*) by “top 3” peptides from the fusion referenced to peptides from full-length qFP-8 and, independently, from BSA (*43*)). Prior to the LC-MS/MS analysis we separated FP-fusions by 1D SDS PAGE and digested corresponding protein bands *in-gel* (**Figure 2A**, Supplementary Figures S3 and S4) to ensure that only full-length proteins were analysed. We underscore that, in contrast to the method *a*), the methods *b*) and *c*) did not rely on the identity of endogenous (from fusions) and standard (from FUGIS or qFP-8) peptides and also used different principles of peptide selection. Consequently, “best 3” (method *b*) and “top 3” (method *c*) peptides could differ even when selected from the same protein digest.

Molar amounts of FP-fusions were determined by the three independent methods with better than 20% precision (**Figure 2C**, Supplementary Dataset S2). This was particularly encouraging since untargeted methods (MBAQ / FUGIS and “top 3”) were designed for rapid proteome-wide quantification. For targeted quantification using qFP-8, coefficients of variance (CVs) among peptide proxies of the same protein were, on average, 14%. Note that the molar amount of the double-tagged FP-fusion #J independently determined using mEGFP and Snap-Tag peptide proxies differed by only 15% (Supplementary Table S4). Depending on the rate of FPs expression (Supplementary Table S5), they could be quantified in extracts of 10 to 100 HeLa cells (Supplementary Figure S5). The sensitivity could reach single-cell level using the method of PRM (*44*).

We note that in the untargeted methods *b*) (“best 3”) and *c*) (“top 3 Hi”) peptides were mostly selected from a non-FP part of FP-fusions. Selecting them only from sequences of FP tags biased the quantification (Supplementary Dataset S2) due to their lower ionization capacity, as indicated by a below-average intensity of their XIC peaks compared to other tryptic peptides. Targeted quantification with FP proxies allowed us to a quantify fusions with proteins that yielded no suitable quantotypic peptides, such as FUS (FP-fusion #B) or Claudin (FP-fusion #C) (Supplementary Dataset S1).

In summary, we concluded that the qFP-8 standard is suitable for the targeted absolute quantification of fusions comprising any combination of FP and non-FP sequences.

### Peptides covering linear and cyclic forms of fluorescent chromophores

Fluorescent chromophore maturation influences FP-fusion detection and quantification by spectroscopy. However, there is currently no generic assay to determine what fraction of the FP-fusion is having the mature fluorescent chromophore (*13, 45*). “Dark” and fluorescent chromophores are structurally distinct (*13, 20, 21, 46*) and it should be possible to quantify both forms in LC-MS/MS experiment.

To this end, in LC-MS/MS analyses of FP digests (Supplementary Table S1, proteins #1-6 and 9) we sought peptides that: *i*) comprise the known chromophore sequence and *ii*) whose masses differ by the equivalent loss of two hydrogen atoms and water (Δm= - 20.031 Da) or four hydrogen atoms and water (Δm= - 22.047 Da). These mass losses are characteristic to the mature cyclic forms of green- and red-type chromophores (**Figure 3A**), respectively (*21, 47-50*). For all seven FPs both linear and cyclic forms of chromophore-containing peptides were detected in FT MS spectra and confirmed by MS/MS (Supplementary Figures S6 and S7). Interestingly, MS/MS spectra of peptides with mature red-type chromophore also contained the characteristic *y-*ion produced by internal cleavage of its cyclic structure (designated “y_i_” in **Figure 3**, Supplementary Figures S6 and S7B). No such cleavage was observed in peptides with green-type chromophores or any intermediate form of the red chromophore.

**Figure 3.**
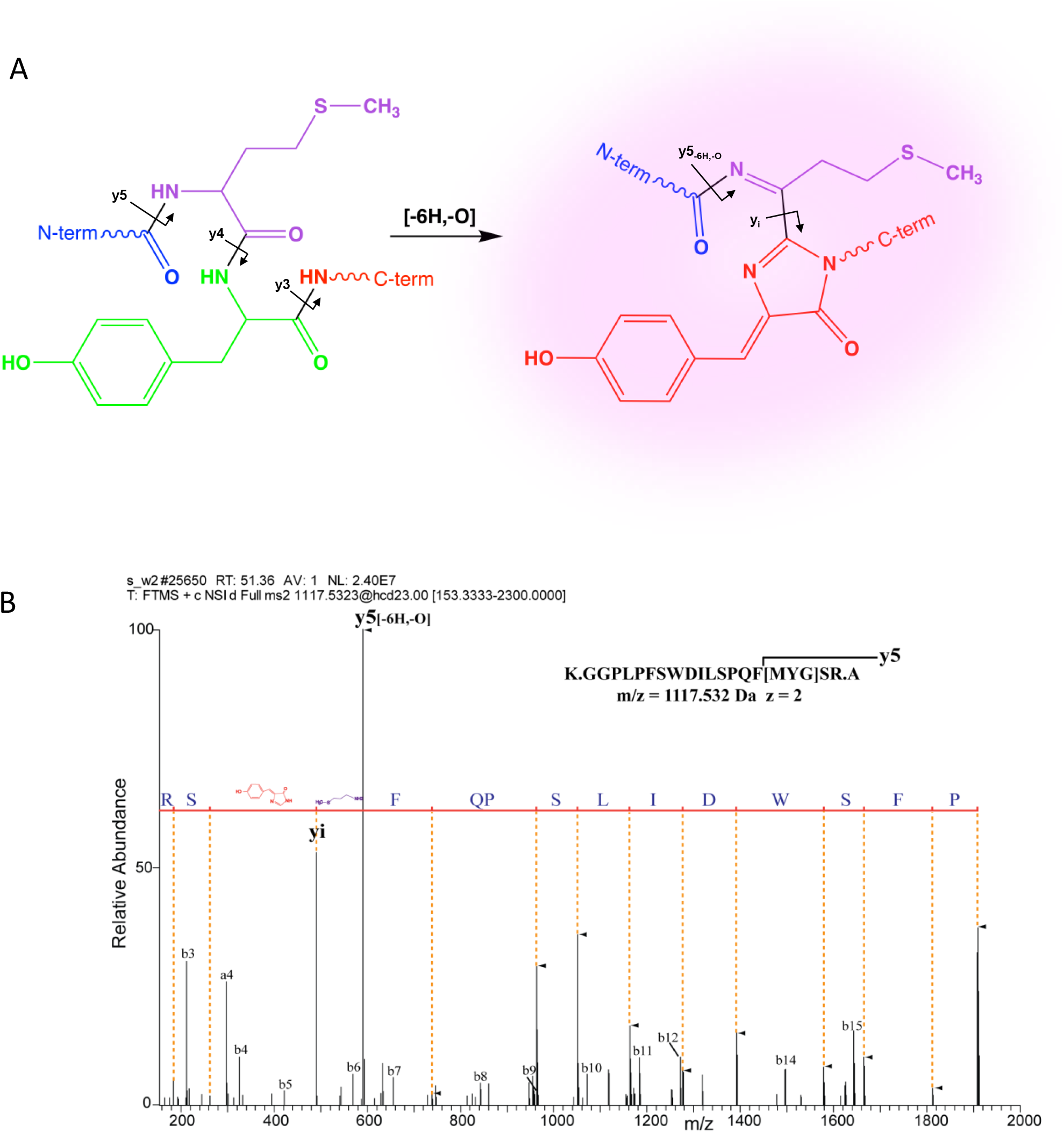
Chemical structure and MS/MS fragmentation of the red-type chromophore containing peptide from mScarlet-I fluorescent protein. **A**: Chemical structures of “dark” (linear) and cyclic (fluorescent) chromophore …MYG… in the GGPLPFSWDILSPQMYGSR peptide (adapted from (*52*)). Arrows indicate peptide bonds cleaved upon MS/MS fragmentation; y_i_ fragment is produced by cleavage of the cyclic chromophore adjacent to a 4-(*p*-hydroxybenzylidene)-5-imidazolinone moiety (in red). **B**: MS/MS spectrum of GGPLPFSWDILSPQMYGSR peptide with the cyclic chromophore. Y-ions comprising the chromophore (starting from y_5_) were detected with a mass shift of Δm= - 22.047 Da compared to corresponding fragments of the linear peptide and are designated with filled triangles. Fragments that do not contain the chromophore (*b*-ions and *y*_*1*_, *y*_*2*_ -ions) have *m/z* expected for the linear form of the peptide.

As expected, in MS/MS spectra we observed the mass shift Δm= - 20.031 Da for y-ions containing cyclic green chromophore, as compared to its linear form. One notable exception was a peptide containing the …TYG… chromophore in EGFP. There corresponding y-ions lost C2H4O moiety (Δm = - 44.026 Da), that most likely originates from the N-terminal threonine residue of the chromophore tripeptide (Supplementary Figure S6B).

Importantly, the ratio of XIC peak intensities of cyclic and linear chromophore-containing peptides corroborated the observed fluorescence. In the red FP mScarlet-I (protein 2, Supplementary Table S1) 72% of GGPLPFSWDILSPQMYGSR peptide (chromophore underlined) contained cyclic chromophore. In contrast, its non-fluorescent mutant M190 (*51*) expressed under the same conditions contained 97% of this peptide with linear chromophore (Supplementary Table S6). Slow maturing FP dsRed-Express protein (protein 9, Supplementary Table S1) contained less than 0.1% of the mature cyclic form (Supplementary Table S6, Supplementary Figure S7), whereas ca. 70% and 29% of its chromophore occurred in the linear and intermediate forms, respectively.

We therefore concluded that the proportion of “dark” and fluorescent fractions together with the total molar abundance of the full-length FP-fusion, can be determined within the same LC-MS/MS analysis and enable quantitative monitoring of FP maturation kinetics.

### Absolute quantification of FP-fusions to study protein expression in cell-free systems

Cell-free expression of FP-fusions is an attractive system for studying the mechanism and kinetics of protein biogenesis (*53-55*) (*56*) that takes advantage of facile and scalable readout by fluorescence spectroscopy. However, physicochemical and structural properties of FPs can compromise the correlation between measured fluorescence and actual concentration of their fusions (*16*). We demonstrate that combining fluorescence spectroscopy with molar quantification of FP-fusions by LC-MS/MS overcomes these analytical challenges. Furthermore, it enables the implementation of a sequential kinetic model to describe the interplay between transcription, translation, and folding.

As a test protein, we chose the human intrinsically disordered protein G3BP1 (Uniprot Q13283). It is a 52 kDa mRNA binding protein essential for liquid-liquid phase separation and formation of stress granules (*57, 58*). The glutamic acid-rich intrinsically disordered region (IDR) of the G3BP1 sequence between amino acid residues 142 - 225 acts as a negative regulator, whose net charge determines the saturation concentration for phase separation with mRNA (*58*). Using fluorescence spectroscopy and quantitative mass spectrometry we compared the yield and kinetics of expression of G3BP1 and its mutant, where glutamic acid residues were removed from the IDR (*58*) changing its formal net charge at pH 7.4 from -17 to +11 (**Figure 4A**, Supplementary Figure S8). For this purpose, the red FP mScarlet-I was fused to the N-terminus of G3BP1 and of its mutant (these FP-fusions are further termed G(WT)-mS and G(mut)-mS for the G3BP1 and the mutant, respectively). The corresponding vectors were expressed in *E.coli* based cell-free expression system PURExpress (further referred to as PURE) and *S. frugiperda* cell extract TnT^®^T7 (further referred to as TnT) (*59, 60*). At chosen time points the expression was quenched and the reaction mixture separated by 1D SDS PAGE. The Coomassie stained band matching the apparent molecular weight (M_r_) of the full length G(WT)-mS or G(mut)-mS proteins was excised, *in-gel* digested and fusion proteins quantified by LC-MS/MS using mScarlet-I peptide proxies from the q-FP8 standard (**Figure 1B**, Supplementary Figure S2). In a parallel experiment, we used a plate reader to record the total fluorescence of the reaction mixture at the channel corresponding to mScarlet-I fluorescence (**Figure 4B**). We chose to include a gel separation step prior LC-MS/MS to consistently compare the levels of fully expressed proteins. Partially expressed fragments with truncated C-terminus could bias LC-MS/MS quantification (*38*), but in these experiments they did not contribute to the total fluorescence (Supplementary Figure S9).

**Figure 4.**
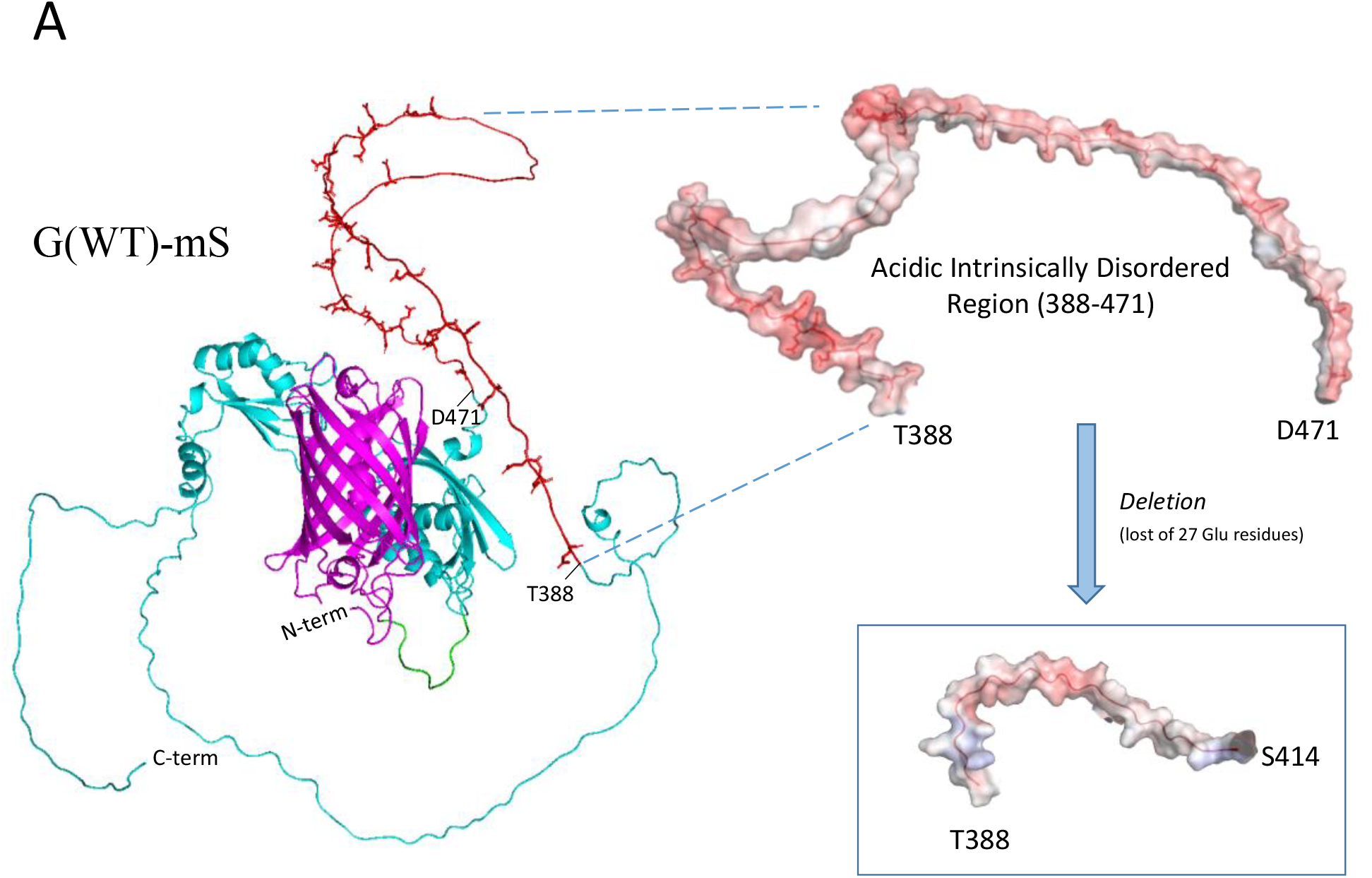
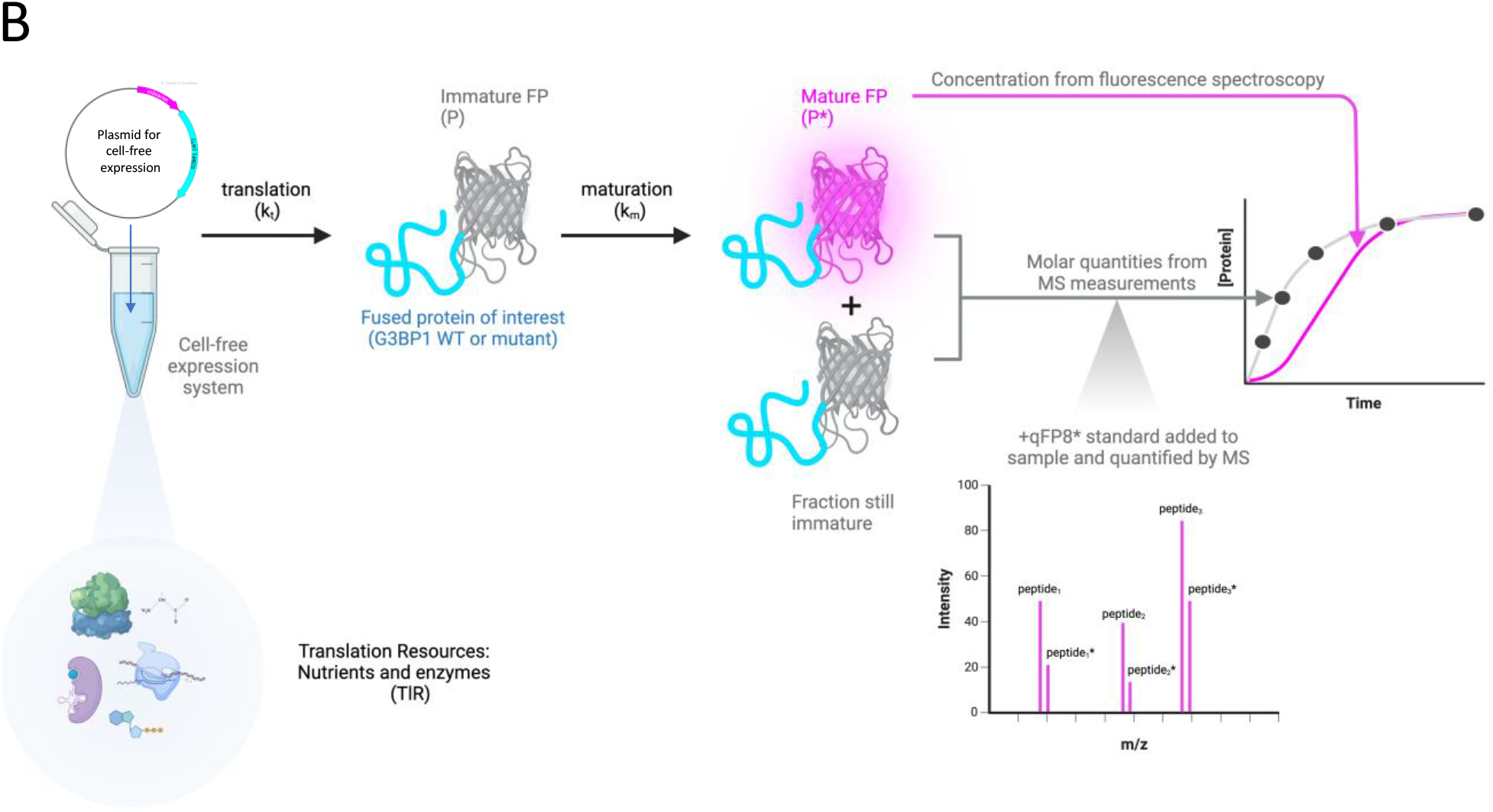
Workflow to study the expression kinetics of G3BP1 and its mutant in cell free protein expression systems. **A**: Predicted 3D structure of G3BP1 fused to mScarlet (G(WT)-mS) and its intrinsically disordered region (IDR) in WT and IDR deletion mutant; mScarlet-I is shown in pink, G3BP1 in magenta, with IDR (side chains of glutamic acid residues extended; red) and a short spacer sequence (green). The inset shows surface charge of the acidic IDR in G(WT)-mS (388-471 aa) before and after removal of glutamic acid enriched sequence stretch (*58*). **B:** Workflow to study translation and maturation kinetics by LC-MS/MS and fluorescence spectroscopy.

G(WT)-mS and G(mut)-mS proteins were quantified by LC-MS/MS using three mScarlet-I peptide proxies in the qFP-8 **(Figure 1**) with inter-peptides coefficient of variation (CV) of less than 15%. Its yield determined by mass spectrometry after 4 hours of protein expression differed significantly between PURE and TnT systems. In PURE the concentration for both proteins exceeded 3 μM, whereas it was *ca*.20-fold lower in TnT (Supplementary Dataset S3). Consistent with the previous reports that eukaryotic cell-free systems could add post-translational modifications to expressed proteins (*61-63*), we observed that G3BP1 expressed in TnT system was phosphorylated at serine residues flanking the acidic IDR, as was also observed in living cells (*57, 64, 65*) (Supplementary Figure S10, Supplementary Table S7). Since phosphorylation could promote G3BP1 dimerization (*58*) and affect protein fluorescence (*66*), we choose the PURE system to study its expression kinetics.

Kinetic curves of both G(WT)-mS and G(mut)-mS showed substantial (ca 35%) difference between protein concentrations measured by mass spectrometry (black line) and by spectroscopy (magenta line) (**Figure 5 A, B**, Supplementary Datasets S3 and S4). A considerable fraction of the FP-fusion failed to make the cyclic fluorescent chromophore even at the end time point (4 hours) of expression and thus remained “dark” (**Figure 3A**). This prompted us to quantify the fraction of fusion protein having the mature cyclic chromophore in its FP tag by mass spectrometry (brown line) from the areas of XIC peaks of precursor ions of cyclic and linear forms of the chromophore-containing peptide (Supplementary Methods; Supplementary Figure S6 and **Figure 3B**)

**Figure 5.**
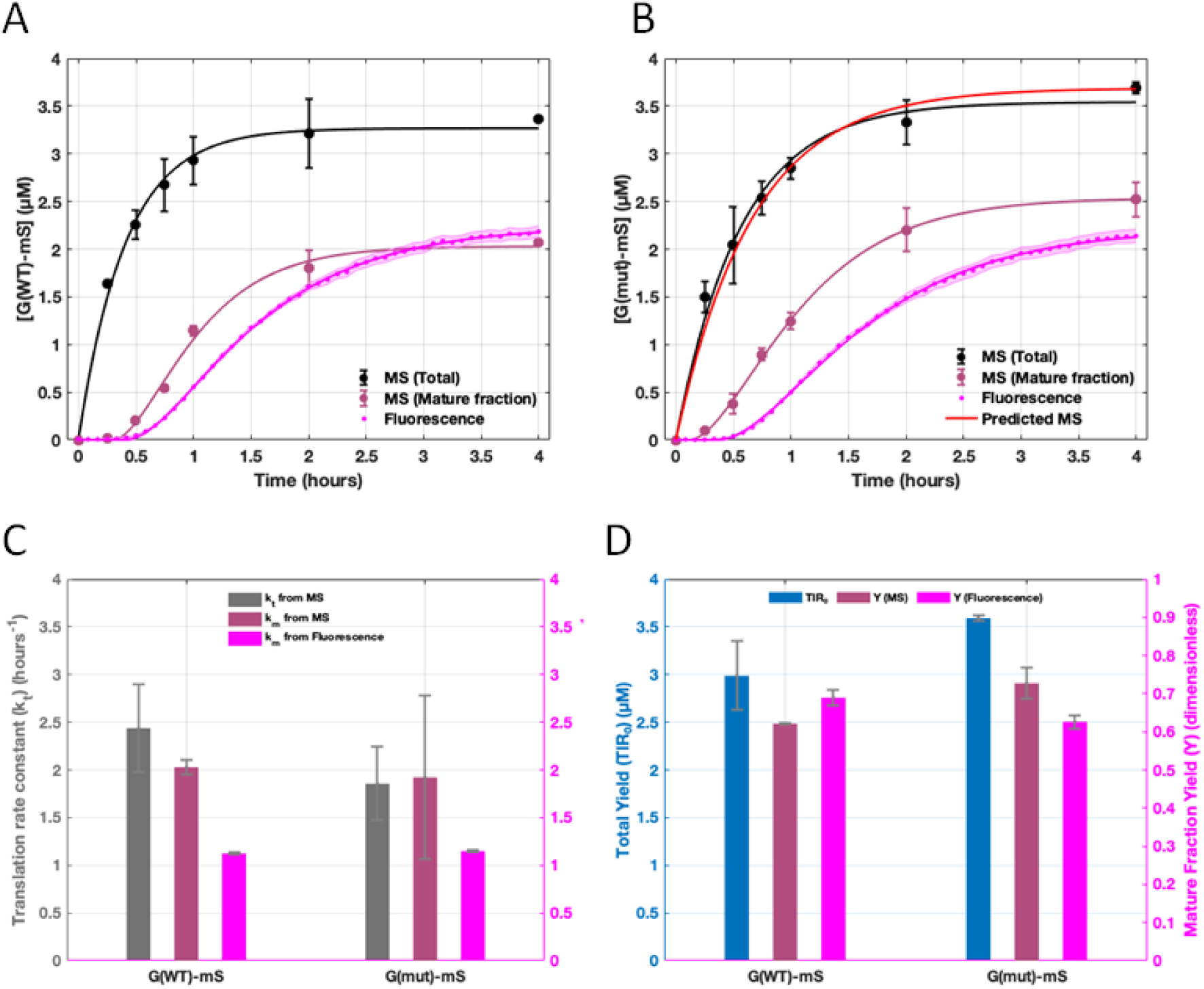
Kinetic traces, constants and yield of the G3BP1 fusions expressed in PURE. Kinetic traces for mScarlet-I fusions of G(WT)-mS (panel **A**) and G(mut)-mS (panel **B**) monitored by mass spectrometry (black) and fluorescence spectroscopy (magenta). Concentration of the fusion having a Scarlet cyclic chromophore determined by mass spectrometry is in brown. Total amount of expressed protein predicted by the kinetic model from fluorescence curve and single mass spectrometry measurement is in red. Translation and maturation rate constants (panel **C**) and overall yields (panel **D**) were calculated by fitting mass spectrometry and fluorescence measurements.

Comparison of the brown and black lines indicated that chromophore maturation was significantly slower than protein translation. Notably, chromophore maturation (brown line) was faster than the increase in detected fluorescence (magenta line). However, by the end point, the entire protein fraction with the mature chromophore became fluorescent. Both the mutant and WT proteins were produced at similar molar concentration and their expression was not considerably affected by the removal of highly acidic IDR.

### Kinetic model for protein expression and maturation of the fluorescent chromophore

Since mass spectrometry quantifies the total amount of fusion protein and fluorescence spectroscopy only quantifies its fraction having FP tag with mature chromophore, we constructed a first-order sequential kinetic model to estimate the rate constants of fusion translation and of chromophore maturation *k*_*t*_ and *k*_*m*_, respectively. A cell free expression system has a finite amount of translation resources (*TlR*). Our model is based on the consideration that an initial translation resource [*TlR*]_0_ is fully converted into protein product, and comprises two consecutive events (**Figures 4B**): *TlR* is translated into the protein with immature chromophore *P* whose fraction (*Y*) becomes the final mature protein *P*_*m*_:

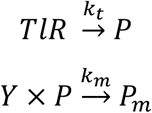

The differential equations associated with the model and their respective solution steps are presented in the Supplementary Methods. From the analytical solutions the mass spectrometry data for total protein [*P*_*T*_] was fit to:

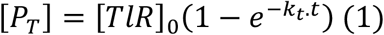

which plateaued at [*TlR*]_0_at equilibrium (*i.e*. t=∞).

The mature fraction as detected by mass spectrometry or by fluorescence spectroscopy was fit to the analytical solution of [*P*_*m*_]:

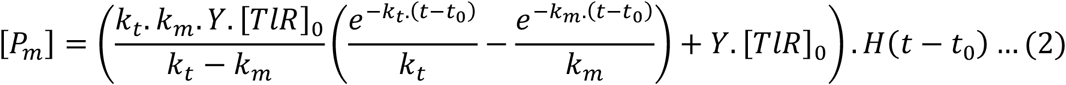

which plateaued at *Y* × [*TlR*]_0_ at equilibrium (*i.e*. t=∞).

For the protein fraction with mature chromophore obtained by mass spectrometry or by fluorescence spectroscopy, a Heaviside step function H was added to fit the data from a time point _*t*0_ from when the signal began to grow to account for hidden processes preceding maturation (**Figure 5A and B**). Having obtained the protein translation rate constant *k*_*t*_ by fitting the mass spectrometry curve with equation (1) it was then used as a known parameter in equation (2). The equation (2) was subsequently used to fit the mature fraction data from mass spectrometry or from the fluorescence spectroscopy to obtain two respective maturation rate constants *k*_*m*_ (**Figure 5C)**.

From the fits for both WT and mutant fusions, the protein translation rate constant *k*_*t*_ calculated for G(WT)-mS was 2.44 ± 0.46 hr^-1^ and its maturation rate constant *k*_*m*_ obtained from fluorescence spectroscopy was 1.12 ± 0.01 hr^-1^, which corresponded to a maturation half-time of 37.1 ± 0.4 minutes (Supplementary Dataset S5). This value is close to, on average, 31.4 minutes reported for the mScarlet-I chromophore (*23, 67, 68*). We speculate that this less than 20% difference in maturation half-times could reflect the impact of unusual highly acidic sequence in G(WT)-mS.

Next, we wanted to test the predictive potential of our model. We asked whether we could simulate a total protein curve for a fusion using previously published maturation half-time estimates. To this end, we took experimentally obtained fluorescence curves and a single data point from mass spectrometry at the 4 hour mark for the G(mut)-mS. The procedure was applied in the reverse order: we used the mScarlet-I maturation half-time value of 31.4± 2.5 minutes (averaged from the values in ref.(*23, 67, 68*)), fit the fluorescence data for the G(mut)-mS and calculated a theoretical translation rate constant (*k*_*t*_). The obtained theoretical *k*_*t*_ of 1.86 ± 0.38 hr^-1^ together with the single mass spectrometric 4^th^ hour data point as [*TlR*]_0_ were subsequently used to predict a total protein curve for G(mut)-mS, which agreed very well with the curve obtained by mass spectrometry (**Figure 5B)**.

Notably, the maturation rate constants for both fusions calculated using mass spectrometry data for mature chromophore fraction (brown curves) were higher than calculated from fluorescence measurements (**Figure 5C**). The corresponding maturation half-times (16.5 ± 0.28 and 21.7 ± 9.59 min for G(WT)-mS and G(mut)-mS, respectively) lesser than the values from the literature (*23, 67, 68*).This suggested that there may be an additional step preceding the formation of the fluorescent fraction of the fusions. To test this assumption, we constructed an extended model (details are in Supplementary Methods) according to the schematic:

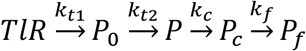

where the total nutrient resource *TlR* is converted into a precursor, *P*_0_, which accounts for the delay in the fluorescence and mature fraction curves. Then it proceeds to the immature fraction P, as in the original model, and then to a fraction *P*_*c*_ having “dark” chromophore, which is then converted to the final fluorescent form *via* a possible folding step, *P*_*f*_. Numerical fits of the same data to the corresponding differential equations are in Supplementary Methods, Supplementary Figure S11 and Supplementary Dataset S5. From the resulting rate constants, an effective rate constant was derived using the formalism:

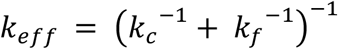

which combines the chromophore maturation and folding steps. The effective rate constant was then compared to the respective maturation rate constant *k*_*m*_ from the original model. The percentage difference between *k*_*eff*_ and *k*_*m*_ was below 10% for both G(WT)-mS and G(mut)-mS, supporting the hypothesis of the sequential maturation process through an intermediate state with a correctly assembled chromophore, but lacking fluorescence (Supplementary Figure S11C).

Overall, the data showed that WT and mutant forms of the intrinsically disordered protein G3BP1 have very close yields of expression (differing by less than 15%) and comparable translation rates (**Figure 5, C**,**D**). This additionally confirms that the change in net charge does not have significant impact on the expression yield and kinetics of the protein. The slightly increased yield of G(mut)-mS might be attributed to its shorter length.

Taken together, our model allows the determination of maturation rate constants for FP-reporters in cell free protein expression systems. It could predict protein expression curves based on a facile fluorescence readout combined with a single protein quantification experiment by mass spectrometry. The model can be instrumental for high throughput screens aiming at or relying upon expression of isoforms, mutants or homologous proteins tagged with the same FP. Finally, the mass-action based reactant-intermediate(s)-product type sequential model can be used to fit any experimental data that follows a similar reaction scheme with an intermediate (here, immature FP-reporter). Although over the course of the reaction the intermediate state is generally invisible the model allows us to infer and track its dynamics (Supplementary Figure S12).

## CONCLUSION AND PERSPECTIVES

We have designed and validated a simple and robust analytical approach for determining the molar abundance of fluorescent protein fusions. It relies on highly expressed isotopically labelled chimeric protein qFP-8 that enables absolute quantification of FP-fusions with 6 major prototypic green and red FPs, 2 self-labelling tags and at least 70 related proteins sharing quantotypic peptides. There is no need to synthesize individual peptide standards or use purified FPs as calibrants and the protocol can be applied to any fused protein with no further adjustments. The same analysis quantifies the molar fraction of mature cyclic chromophore, which is directly relevant to visible fluorescence. We envision that this technically straightforward proteomics method will support diverse applications, including quantifying expressed FP-fusions and assessing their maturation rate in cells and tissues or using FP-fusions as quantitative reporters, among others.

We further combined mass spectrometry-based quantification with fluorescence spectroscopy to monitor the kinetics of fusions expression in cell free systems. Effectively, we were able to delineate the kinetics of protein translation, chromophore maturation and formation of genuine fluorescent protein and integrate these multifaceted kinetic data into a single mathematic model. Together with computational prediction of protein structures this could advance our understanding of the interplay between protein sequence, translation, folding and structure.

Finally, we envision that FP-fusions quantification at the single cell level will expand analytical scope of system and synthetic biology by aligning the spatial precision of fluorescent microscopy with exact molar quantities of imaged proteins. It is also intriguing to explore if FP-fusions with precisely known molar quantities could serve as a generic internal standard (*42*) to determine the molar abundance of endogenous proteins detected in the same microscopy experiment and contribute to understanding molecular composition of labile protein condensates (*69*).

## EXPERIMENTAL

**Abbreviations:** BSA, bovine serum albumin; DDA, data dependent acquisition; FP, fluorescent protein; FastCAT, Fast-track QconCAT analytical protocol; FT MS, Fourier Transform mass spectrometry; FUGIS, fully unlabelled generic internal standard; HCD, higher energy collision induced dissociation; LC-MS/MS, liquid chromatography - tandem mass spectrometry; MS Western, mass spectrometry driven Western blotting; PRM, parallel reaction monitoring; SD, standard deviation; XIC, extracted ions chromatogram.

**Chemicals and key resources** are in Supplementary Methods.

### Preparation of chimeric protein standards, recombinant FPs and FP-fusions

Proteins were expressed in *E. coli*, FP-fusions – in insect cells (*Trichoplusia ni*); qFP-8 was metabolically labelled as described in (*35, 42*). Cells were lysed, aliquots of whole lysates separated by 1D SDS PAGE and visualized by Coomassie staining. The positions of full-length FP-fusions at the gel were determined by fluorescence gel imaging and by Western blotting using a primary antibody directed against the HRV 3C-site. For quantification of FP-fusions in method validation experiment, gel bands corresponding to the fusion, standard chimeric proteins qFP-8 and FUGIS, and 1pmol of the reference protein BSA were co-digested *in-gel* with trypsin. For detection of chromophore-covering peptides, depending on the FP sequence, corresponding FP bands were *in-gel* digested by trypsin or by Asp-N protease. For quantification of FP expression in a single cell, HeLa and HCT116 cell stably transfected with FPs were counted prior lysis and then processed as above.

The resulting peptide mixtures were extracted, dried down and subjected to LC-MS/MS analysis.

Details of the sample preparation for mass spectrometric analysis are described in Supplementary methods.

### Protein expression kinetics study in cell-free systems

Plasmids coding for G3BP1 and the mutant N-terminally fused with mScarlet-I were designed and optimized for insect and bacterial expression systems as described in Supplementary Methods. The plasmids were then used in PURExpress® and TnT® T7 cell-free Protein Expression Systems accordingly to the manufacturer protocol. For absolute quantification by mass spectrometry using qFP-8 chimeric standard, 10 µL aliquots were withdrawn with 15-30-minute interval from the master solution, quenched with 10 µL Laemmli buffer and separated by SDS PAGE. Gel regions at around 80kDa and 73kDa corresponding to G(WT)-mS and G(mut)-mS were *in-gel* digested with trypsin; qFP-8 standard and reference BSA were separately co-digested *in-gel* and aliquots spiked into digests of the model proteins prior LC-MS/MS analysis.

In parallel, the expression of mScarlet tagged G3BP1 variants was tracked on a TECAN Spark 20M plate reading fluorimeter. After initiation of expression fluorescent signal was measured during 4 hours at 569/594 nm excitation/emission wavelength. Obtained arbitrary fluorescence units were converted to protein concentration using calibration curves built with purified mScarlet-I protein.

### Mass spectrometry analysis and absolute quantification

Peptide mixtures were analyzed by LC-MS/MS on a nanoUPLC Ultimate 3000 interfaced to an Orbitrap HF hybrid mass spectrometer; MS3 experiment was carried on a LTQ Orbitrap Velos mass spectrometer (all Thermo Scientific, Bremen). Data were acquired in DDA mode and in DDA with inclusion list of m/z of precursor (see Supplementary Methods). To avoid carryover, 2-3 blank runs were performed after each sample analysis.

Spectra were processed with FragPipe software suit v.17.1 (*70*) or matched by Mascot software (v.2.2.04, Matrix Science). Intensities of native and isotopically labelled peptide precursors were extracted from FragPipe output by in-house scripts. Absolute quantification using isotopically labelled chimeric standard qFP-8 and FUGIS was performed as described in (*35, 42*).

**Modelling procedures** are described in Supplementary Methods

## ACKNOWLEDGEMENTS

We are grateful to Dr.R.Barsacchi (Technology Development Studio, MPI-CBG), Dr.M.Sarov (Genome Engineering Facility, MPI-CBG) for providing transfected cells; Prof.A.Hyman (MPI-CBG), Prof.A.Honigmann (TUD), Prof. S. Diez (TUD), Prof. S. Alberti (TUD) and Dr.L.Vogeley for providing FP-fusion constructs; members of Shevchenko laboratory for useful discussion.

## DATA AVAILABILITY

Source files with LC-MS/MS spectra in *.raw format are available at Edmond repository (*67*) URL: https://doi.org/10.17617/3.UXWVQE

