## Supplementary Figures and Tables for "Absolute Quantification of Fluorescent Protein Fusions by Proteomics"

#### Supplementary Dataset S1

List of FP-Fusions

| FP-fision code | Fused Tag | shuttle vector construct name[1] | V(PEPC)[2] | D(PEPC)[2] | FP-fusion sequence | MW, kDa | proprietor |
|---|---|---|---|---|---|---|---|
| | | Expression in *T.ni* insect cells[3] | | | | | |
| #A | mKate2 | TH1510-(pOCC205)_pOEM1-TwinStrep-fly_Sp | V3970 | D4549 | MGSAWSHPQFEKGGGSGGGSGGSAWSHP | 158 | *Prof. A.Hyman, MPI-CBG* |
| #B | SNAP-tag | TH1968-(pOCC177)_pOEM1-N-HIS-MBP-PS-F | V5670 | D5727 | MGSSHHHHHHSSGRMKIEEGKLVIWING | 119 | *Prof. A.Hyman, MPI-CBG* |
| #H | SNAP-tag | pOCC176-ZO2 (HIS6-MBP-3C-SNAP-TEV-ZO2) | V5851 | D3399 | MGSSHHHHHHSSGRMKIEEGKLVIWING | 200 | *Prof.A.Honigman, TUD* |
| #C | HaloTag | OCC352-hs_Claudin2-stop(HALO) | V5713 | D5753 | MGSSHHHHHHSSGRLEVLFQGPMAEIGT | 62 | *Prof.A.Honigman, TUD* |
| #D | mNeonGreen | pDL372_286_KIF5B short-C-mNeonGreen-3C- | V5784 | D5821 | MAAAMADLAECNIKVMCRFRPLNESEVN | 134 | *Prof. A.Hyman, MPI-CBG* |
| #E | mScarlet-I | pDL375_FLKif16b-mScarlet-MBP | V5785 | D5822 | MAAAMASVKVAVRVRPMNRREKDLEAKF | 223 | *Prof. S.Diez, TUD* |
| #F | Dendra2 | TH_Tau40_Dendra2_PS_6xhis | V5849 | D2918 | MAAAAEPRQEFEVMEDHAGTYGLGDRKD | 75 | *Prof. A.Hyman, MPI-CBG* |
| #G | mEGFP | pOCC119-ZO2 (HIS6-MBP-ZO2-mGFP) | V5850 | D3307 | MGHHHHHHSSGRMKIEEGKLVIWINGDK | 207 | *Prof.A.Honigman, TUD* |
| #I | mChery | pOCC172-ZO2 (HIS6-MBP-3C-Cherry-ZO2) | V5852 | D3450 | MGSSHHHHHHSSGRMKIEEGKLVIWING | 207 | *Prof.A.Honigman, TUD* |
| #J | mEGFP, Snap-tag | SNAP-nmy2-mGFP | V5728 | D5766 | MGMDKDCEMKRTTLDSPLGKLELSGCEQ | 280 | *Dr. L.Vogeley, MPI-CBG* |
| | | Cell-free expression experiment[4] | | | | | |
| G(WT)-mS _TNT | mScarlet-I | pOCC189-mScarletI-G3BP1(WT) L515 | | | MVSKGEAVIKEFMRFKVHMEGSMNGHEF | 80 | *this work* |
| G(WT)-mS _PURE | mScarlet-I | pEXP5NT-mScarletI-G3BP1(WT) | | | MVSKGEAVIKEFMRFKVHMEGSMNGHEF | 80 | *this work* |
| G(mut)-mS _PURE | mScarlet-I | pEXP5NT-mScarletI-G3BP1(ΔE1ΔE2) | | | MVSKGEAVIKEFMRFKVHMEGSMNGHEF | 73 | *this work* |

1- as given by proprietor

2 - Accession Number in MPI-CBG Protein Biochemistry Facility Database for viruses (V) and plasmids(D)

3 - designed based on pOEM1 as described <<https://doi.org/10.1186/s12896-019-0512-z>>

4- designed in this study

#### Supplementary Dataset S2

Absolute Quantities of FP-Fusions obtained by ms-based Top3-Hi *(ref.41)* and MBAQ *(ref.40)* approaches

| Approach | FP-fusion amount, pmol[1] | | | | | | | | |
|---|---|---|---|---|---|---|---|---|---|
| | #A | #B | #C | #D | #E | #F | #G | #H | #I |
| Top3-Hi (with BSA as standard protein) | 1.239±0.118 | 0.442±0.077 | 1.107±0.132 | 0.356±0.017 | 0.142±0.029 | 0.020±0.000 | 2.279±0.293 | 2.254±0.238 | 3.052±0.272 |
| Top3-Hi (with qFP-8 as standard protein) | 1.106±0.112 | 0.418±0.067 | 1.009±0.057 | 0.349±0.036 | 0.132±0.022 | 0.022±0.001 | 2.257±0.183 | 2.003±0.207 | 3.089±0.201 |
| MBAQ[2] | 1.102±0.069 | 0.395±0.057 | 1.233±0.126 | 0.336±0.020 | 0.162±0.058 | 0.015±0.002 | 1.755±0.180 | 2.189±0.206 | 2.704±0.789 |
| MBAQ for FP-tag[3] | 0.205±0.008 | 0.130±0.016 | 0.492±0.067 | *0.072±0.051* | *0.050±0.020* | | *0.040±0.019* | 0.922±0.206 | *0.052±0.042* |

[1]with Stdev

[2] “best3” peptide references are selected from full length sequence of FP-fusions

[3] “best3” peptide references are selected from FP-tag part of the FP-fusion sequence. Values in *Italic* indicates that that calculations are made with “best 2” peptides.
No reference peptide with concordant abundance were detected for fusion #F.

### Supplementary Dataset S3

**Concentration of the model FP-fusions in PURExpress cell-free expression system obtained by mass spectrometry**

**G(WT)-mS experiment**

| | | G(WT)-mS total, µM | | | | | with mature chromophore, µM | | | | | |
|---|---|---|---|---|---|---|---|---|---|---|---|---|
| Time point, min | Replicate 1 | Replicate 2 | Replicate 3 | Replicate 4 | Mean W | Std Err W | Mature Rep 1 | Mature Rep 2 | Mature Rep 3 | Mature Rep 4 | Mean Mat | Std Err Mature |
| 0 | 0 | 0 | 0 | 0 | 0 | 0 | 0 | 0 | 0 | 0 | 0 | 0 |
| 15 | 1.633851632 | 1.616275865 | 1.732293398 | 1.556644632 | 1.634766382 | 0.036466038 | 0.010290042 | 0.018434025 | 0.01690059 | 0.021770714 | 0.016849 | 0.002411 |
| 30 | 2.542953967 | 1.994811837 | 2.482816364 | 2.012111464 | 2.258173408 | 0.147611602 | 0.177643265 | 0.210719125 | 0.205329847 | 0.230208326 | 0.205975 | 0.010851 |
| 45 | 3.218203298 | 2.210308002 | 3.061927189 | 2.19667465 | 2.671778285 | 0.272255203 | 0.573742326 | 0.484441873 | 0.607034308 | 0.521926591 | 0.546786 | 0.027174 |
| 60 | 3.374522523 | 2.537288043 | 3.35938704 | 2.452249934 | 2.930861885 | 0.252394918 | 1.232133249 | 1.052335759 | 1.249886776 | 1.038579813 | 1.143234 | 0.056637 |
| 120 | 3.886266498 | 2.59903173 | 3.809215848 | 2.557320707 | 3.212958696 | 0.366927954 | 2.142554329 | 1.506758929 | 2.120365961 | 1.45141771 | 1.805274 | 0.188716 |
| 240 | 3.342711774 | | 3.385760195 | | 3.364235984 | 0.02152421 | 2.041803576 | | 2.093283139 | | 2.067543 | 0.02574 |

**G(mut)-mS experiment**

| | | G(mut)-mS total, µM | | | | | with mature chromophore, µM | | | |
|---|---|---|---|---|---|---|---|---|---|---|
| Time point, min | Replicate 1 | Replicate 2 | Replicate 3 | Replicate 4 | Mean Mut | Std Err Mut | Mature Rep 1 | Mature Rep 2 | Mean Mature | Std Err Mature |
| 0 | 0 | 0 | 0 | 0 | 0 | 0 | 0 | 0 | 0 | 0 |
| 15 | 1.73619583 | 1.225726647 | 1.815454365 | 1.210941289 | 1.497079533 | 0.161773149 | 0.110397316 | 0.093881801 | 0.102139558 | 0.008257758 |
| 30 | 2.781744812 | 1.37609366 | 2.703251286 | 1.321069925 | 2.045539921 | 0.402864427 | 0.480109065 | 0.27235974 | 0.376234403 | 0.103874663 |
| 45 | 2.6994863 | 2.21730557 | 2.96639411 | 2.264070518 | 2.536814125 | 0.179693255 | 0.962869674 | 0.832808067 | 0.89783887 | 0.065030804 |
| 60 | 3.021522196 | 2.671827094 | 3.063980153 | 2.633340204 | 2.847667412 | 0.113237456 | 1.33581111 | 1.157376914 | 1.246594012 | 0.089217098 |
| 120 | 3.685916547 | 2.960692598 | 3.782527026 | 2.90540775 | 3.33363598 | 0.232391723 | 2.426746577 | 1.972990031 | 2.199868304 | 0.226878273 |
| 240 | 3.552523524 | 3.801477068 | 3.649777484 | 3.750675986 | 3.688613516 | 0.055240878 | 2.343503794 | 2.702796823 | 2.523150308 | 0.179646515 |

**Concentration of the model FP-fusions in TnT®T7 (P. frugiperda) cell-free expression system at 4h time point obtained by mass spectrometry, µM**

| G(WT)-mS | G(mut)-mS |
|---|---|
| 0.17 ± 0.03 | 0.19±0.02 |

#### Supplementary Dataset S4

**Concentration of the model FP-fusions in PURExpress cell-free expression system obtained by fluorecsent spectroscopy**

| | G(WT)-mS, µM | | | G(mut)-mS, µM | | |
|---|---|---|---|---|---|---|
| Time, h | [P*] rep 1 | [P*] rep 2 | [P*] rep 3 | [P*] rep 1 | [P*] rep 2 | [P*] rep 3 |
| 0 | 0.0058 | 0.015 | 0.0035 | 0.0007 | 0.004 | 0.0054 |
| 0.08349417 | 0.0058 | 0.0117 | 0.0014 | 0.0008 | 0.0066 | 0.0085 |
| 0.16682472 | 0.0064 | 0.0079 | 0.0053 | 0 | 0.0035 | 0.0032 |
| 0.25015417 | 0.0079 | 0.0192 | 0.0008 | 0.0001 | 0.0094 | 0.0025 |
| 0.33348667 | 0.0077 | 0.0243 | 0.0048 | 0.0017 | 0.0119 | 0.0184 |
| 0.41681528 | 0.0233 | 0.0308 | 0.0158 | 0.0153 | 0.008 | 0.0248 |
| 0.50014333 | 0.0451 | 0.0576 | 0.0365 | 0.0384 | 0.0322 | 0.0442 |
| 0.5834775 | 0.0885 | 0.1001 | 0.0752 | 0.0773 | 0.065 | 0.0878 |
| 0.66680556 | 0.149 | 0.1634 | 0.1359 | 0.1327 | 0.1196 | 0.1452 |
| 0.75013361 | 0.2275 | 0.2501 | 0.2125 | 0.2096 | 0.1912 | 0.2213 |
| 0.83346667 | 0.3256 | 0.3463 | 0.312 | 0.3002 | 0.2798 | 0.3076 |
| 0.91679528 | 0.4292 | 0.4529 | 0.4149 | 0.4021 | 0.3864 | 0.423 |
| 1.00012472 | 0.5524 | 0.5791 | 0.5321 | 0.5023 | 0.4695 | 0.5264 |
| 1.083455 | 0.6561 | 0.6852 | 0.6288 | 0.6082 | 0.5771 | 0.6372 |
| 1.16678528 | 0.77 | 0.7982 | 0.7489 | 0.7053 | 0.6708 | 0.7313 |
| 1.25011361 | 0.8781 | 0.9244 | 0.8416 | 0.8116 | 0.7632 | 0.847 |
| 1.33344361 | 0.978 | 1.0029 | 0.9505 | 0.8894 | 0.8607 | 0.9188 |
| 1.41677222 | 1.0786 | 1.1077 | 1.0517 | 0.9878 | 0.954 | 1.0193 |
| 1.50010444 | 1.1752 | 1.204 | 1.149 | 1.0793 | 1.0374 | 1.1055 |
| 1.58343694 | 1.2416 | 1.2755 | 1.2129 | 1.1422 | 1.107 | 1.1769 |
| 1.66676444 | 1.3276 | 1.3633 | 1.2924 | 1.2098 | 1.1643 | 1.2501 |
| 1.750095 | 1.4005 | 1.4417 | 1.3666 | 1.2929 | 1.2495 | 1.3322 |
| 1.83342528 | 1.4724 | 1.5206 | 1.4301 | 1.3552 | 1.3021 | 1.3969 |
| 1.91675333 | 1.5193 | 1.5681 | 1.4754 | 1.4225 | 1.3639 | 1.4718 |
| 2.00008583 | 1.5992 | 1.6643 | 1.5507 | 1.4853 | 1.4176 | 1.5524 |
| 2.08341583 | 1.637 | 1.7018 | 1.5798 | 1.542 | 1.47 | 1.608 |
| 2.16674417 | 1.6789 | 1.7569 | 1.6156 | 1.5716 | 1.4922 | 1.6452 |
| 2.25007694 | 1.7374 | 1.8105 | 1.6727 | 1.6377 | 1.5607 | 1.7157 |
| 2.33340556 | 1.7599 | 1.8408 | 1.6857 | 1.6712 | 1.576 | 1.7552 |
| 2.41673639 | 1.8301 | 1.9118 | 1.7604 | 1.7145 | 1.6227 | 1.8022 |
| 2.50006889 | 1.8589 | 1.9407 | 1.7867 | 1.7474 | 1.6588 | 1.836 |
| 2.58339556 | 1.896 | 1.9867 | 1.8218 | 1.7774 | 1.6881 | 1.8666 |
| 2.66672583 | 1.9109 | 1.9959 | 1.8394 | 1.8237 | 1.7374 | 1.909 |
| 2.75005583 | 1.9501 | 2.0359 | 1.8773 | 1.8711 | 1.7779 | 1.9617 |
| 2.83338444 | 1.9727 | 2.0526 | 1.9011 | 1.8881 | 1.7992 | 1.969 |
| 2.916715 | 2.0037 | 2.0938 | 1.9264 | 1.9165 | 1.8185 | 2.0041 |
| 3.00004778 | 2.0131 | 2.1005 | 1.9395 | 1.9556 | 1.867 | 2.0468 |
| 3.08337556 | 2.0583 | 2.1368 | 1.9894 | 1.9653 | 1.8741 | 2.0494 |
| 3.16670667 | 2.0796 | 2.1686 | 2.0093 | 1.9766 | 1.8903 | 2.0621 |
| 3.25003361 | 2.0663 | 2.16 | 1.9894 | 2.0065 | 1.9093 | 2.0942 |
| 3.33336556 | 2.0859 | 2.1715 | 2.0091 | 2.0309 | 1.9315 | 2.1204 |
| 3.41669222 | 2.1167 | 2.2016 | 2.0343 | 2.0376 | 1.9396 | 2.1279 |
| 3.50002194 | 2.1245 | 2.2144 | 2.05 | 2.0699 | 1.9769 | 2.1588 |
| 3.58335056 | 2.1349 | 2.2286 | 2.0583 | 2.0844 | 1.9876 | 2.1741 |
| 3.66668111 | 2.1315 | 2.2173 | 2.0534 | 2.11 | 2.017 | 2.1933 |
| 3.75001028 | 2.1598 | 2.252 | 2.0761 | 2.1129 | 2.01 | 2.2055 |
| 3.83334028 | 2.1505 | 2.2401 | 2.0768 | 2.131 | 2.0381 | 2.2189 |
| 3.91667167 | 2.1604 | 2.2475 | 2.0823 | 2.1331 | 2.0324 | 2.2278 |
| 4.00000306 | 2.1824 | 2.2676 | 2.1012 | 2.1429 | 2.0381 | 2.2337 |

### Supplementary Dataset S5

#### Model Parameters

##### Original model parameters

| Sample | TlR_0 (µM) | Y1 | Y2 | kt (1/hr) | km1 (1/hr) | km2 (1/hr) |
|---|---|---|---|---|---|---|
| G(WT)-mS | 2.99±0.36 | 0.59±0.01 | 0.76±0.02 | 3.47±0.46 | 3.45±0.08 | 1.02±0.01 |
| G(mut)-mS | 3.59±0.03 | 0.73±0.04 | 0.63±0.02 | 1.86±0.39 | 1.92±0.86 | 1.15±0.01 |

| **Legend** | **Explanation** |
|---|---|
| TlR_0 | Initial translation resource |
| Y1 | Yield from mature fraction |
| Y2 | Yield from fluorescence |
| kt | Translation rate constant |
| km1 | Maturation rate constant from mature fraction |
| km2 | Maturation rate constant from fluorescence |

##### Extended model parameters

| Sample | kt1 (1/hr) | kt2 (1/hr) | kc (1/hr) | kf (1/hr) | TlR_0 (µM) | X | Y | km_eff (1/hr) | km_true (1/hr) | Difference (%) |
|---|---|---|---|---|---|---|---|---|---|---|
| G(WT)-mS | 3.98±0.5 | 2.44±0.02 | 2.35±0.02 | 1.95±0.19 | 3.27±0.37 | 0.64±0.0001 | 0.67±0.11 | 1.07±0.06 | 1.02±0.01 | 4.96 |
| G(mut)-mS | 1.76±0.01 | 2.22±1.43 | 4.21±0.03 | 1.81±0.17 | 3.55±0.02 | 0.71±0.03 | 0.61±0.0001 | 1.26±0.08 | 1.15±0.01 | 10.03 |

| **Legend** | **Explanation** |
|---|---|
| TlR_0 | Initial translation resource |
| kt1 | Rate constant for TlR_0 to P_0 |
| kt2 | Rate constant for P_0 to P |
| kc | Rate constant for P to P_c |
| kf | Rate constant for P_c to P_f |
| X | Fraction of P converting to P_c |
| Y | Fraction of P_c converting to P_f |
| km_eff | Effective maturation rate constant |
| km_true | Maturation rate constant from original model |
| Difference | Percentage difference between km_eff and km_true |
