## Supplementary Methods for "Absolute Quantification of Fluorescent Protein Fusions by Proteomics"

1. Key resources table
2. Preparation of chimeric protein standards and model proteins
   *2.1 Protein expression*
   *2.2 Cell lysis*
   *2.3 SDS PAGE electrophoresis*
   *2.4 In-gel fluorescence imaging and Western Blotting for SDS-PAGE-separated model FP-fusions*
3. Sample preparation for mass spectrometric analysis
   *3.1 for peptide mapping of FPs and FP-fusions*
   *3.2 for ms-based absolute quantification of FP-fusions using qFP-8 standard*
   *3.3 for ms-detection of chromophore-containing peptides*
   *3.4 for ms-based quantification of FP expression in cells*
   *3.5 for ms-based quantification of protein expression kinetics in cell free systems*
4. Mass Spectrometric analysis and data processing
   *4.1 LC-MS/MS analysis*
   *4.2 MS data processing and absolute quantification*
5. Plasmid preparation for cell-free expression
   *5.1 Plasmid Design*
   *5.2 Restriction Digestion-Ligation of DNA*
   *5.3 Agarose Gel Electrophoresis*
   *5.4 DNA Isolation and Gel Extraction*
   *5.5 Transformation of DNA into Competent Cells for storage and amplification*
6. Quantitative Fluorescence Measurement for cell-free expressed mScarlet-I tagged G3BP1
   *6.1 Preparation of mScarlet-I standard*
   *6.2 Fluorescence assay*
   *6.3 Calibration of fluorescence data*
7. Protocols for generating 3D structures for mScarlet-I tagged G3BP1 and its mutant with deleted IDR
8. Kinetic Model
   *8.1 Original model*
   *8.2 Extended model*
9. Kinetic curves fitting and simulations

### 1. Key resources table

| Reagents or resources | Source | Identifier |
|---|---|---|
| **Antibodies** | | |
| HRV3C Polyclonal Antibody | Invitrogen | PA1-118 |
| IRDye® 800CW Goat anti-Rabbit IgG Secondary Antibody | Licorbio | P/N: 926-32211 |
| **Bacterial strains** | | |
| BL21(DE3) competent *E.coli* | New England Biolabs | C2527I |
| 5-alpha Competent *E. coli* | New England Biolabs | C2987H |
| T7 Express Competent *E. coli* | New England Biolabs | C2566H |
| **Chemicals, peptides, and recombinant proteins** | | |
| Trypsin, sequencing grade | Promega | V5111 |
| Trypsin/Lys-C Mix, Mass Spec Grade | Promega | V5073 |
| AspN | Promega | V162A |
| C4Life Standard Mix | Promega | CS30240 |
| 100x Halt™ Protease Inhibitors Cocktail | ThermoFischer Scientific | 87786 |
| BSA standard | ThermoFischer Scientific | 23209 |
| Methanol, Optima™ LC/MS-grade | FisherScientific | 10636545 |
| Acetonitrile, Optima™ LC/MS-grade | FisherScientific | 10055454 |
| Water for chromatography LC-grade | Merck | 1.15333 |
| Formic acid 98-100% | Merck | 1.00264.0100 |
| Acetic Acid | Karl Roth | 6755.2 |
| Ammonium bicarbonate | Sigma-Aldrich | A6141 |
| Isopropanol | Honeywell Research Chemicals | 34965 |
| DL-dithiotheitol (DTT) | Sigma-Aldrich | D5545-1G |
| Jodacetamid | Sigma-Aldrich | I1149-5G |
| Coomassie Brilliant Blue R250 | Serva | 17525.02 |
| Tris-Glycine/SDS electrophoresis buffer (10x) | Serva | 42529.01 |
| Laemmli Sample Buffer (x2) for SDS-PAGE | Serva | 42526.01 |
| Prestained Protein Standards SeeBlue™ Plus2 | Invitrogen | LC5925 |
| Precision Plus Protein™ Unstained Protein Standards | BioRad Laboratories | 1610363 |
| 2-Mercaptoethanol | Sigma-Aldrich | M3148-25ML |
| LB medium | In-house prepared | N/A |
| Phosphate-buffered saline (PBS) | In house prepared | N/A |
| Dulbeccos Modified Eagle Medium (DMEM) | Thermo Fischer Scientific | 31966 |
| Terrific Broth medium | In-house prepared | N/A |
| Fetal Bovine Serum (FBS) | Merck | S0615 |
| Geneticin | ThermoFischer Scientific | 10131-019 |
| x10 rCutSmart™ Buffer | New England Biolabs | B6004S |
| T4 DNA ligase buffer | New England Biolabs | B0202S |
| T4 ligase | New England Biolabs | M0202S |
| Gel Loading Dye, Purple (6x) | New England Biolabs | B7025 |
| Quick-Load© 1 kb DNA Ladder | New England Biolabs | N0468S |
| NEB® 5-alpha | New England Biolabs | C2987H |
| Kanamycine | Sigma-Aldrich | K4000 |
| Ampicillin | Sigma-Aldrich | A9518 |
| Chloramphenicol | Calbiochem | 220551 |
| Isopropyl ß-D-1-thiogalactopyranoside (IPTG) | Fermentas | R 0392 |

| | | |
|---|---|---|
| Agarose | Invitrogen | 16500-500 |
| TAE buffer | In house | N/A |
| GelRed Nucleic Acid Stain 10,000X Water | Sigma-Aldrich | SCT123 |
| Gel Loading Dye, Purple (6x) | New England Biolabs | B7024S |
| Quick-Load® Purple 1 kb Plus DNA Ladder | New England Biolabs | N0550S |
| LB broth | In house | N/A |
| LB agar | In house | N/A |
| ESF 921 serum-free medium | OET | 500300 |
| 2x Laemmli protein sample buffer | Bio-Rad Laboratories | 1610737 |
| Sodium dodecylsulfate (SDS) | Serva | 20765 |
| Nonidet™ P-40 Alternative | Sigma-Aldrich | 74385 |
| Tween-20 | Karl Roth | 9127.1 |
| Imidazole | MP Biomedicals | 102033 |
| Recombinant proteins used in this study are listed in Supplementary Table S2 | This study | N/A |
| **Commercial assays** | | |
| TnT® T7 Insect Cell Extract Protein Expression System | Promega | L1101 |
| PURExpress® In Vitro Protein Synthesis Kit | New England Biolabs | E6800S |
| QIAprep Spin Miniprep Kit (50) | Qiagen | 27104 |
| QIAGEN Plasmid Maxi Kit (10) | Qiagen | 12162 |
| QiaQuick Gel Extraction Kit | Qiagen | 28704 |
| **Cell lines** | | |
| HeLa Kyoto | https://www.cellosaurus.org/CVCL_1922 | RRID:CVCL_1922 |
| HCT116 cells | Gift from propietor Dr.S.Sarov (MPI-CBG) | N/A |
| Tni cells (*Trichoplusia ni)* | https://www.cellosaurus.org/CVCL_C412 (cat# 94-002 from Expression Systems) | RRID:CVCL_C412 |
| **Oligonucleotides** | | |
| Primers used for cloning in cell-free experiment are listed in SM Chapter 5.1 | This study | N/A |
| **Recombinant DNA** | | |
| pET11Kan expression vector | NovoPro | V014870 |
| pEGFP-C1 expression vector | NovoPro | V012024 |
| pEXP5-NT expression vector | Invitrogen | V960-05 |
| pOCC expression vectors | PEPC of MPI CBG | N/A |
| Shuttle constructs used in this study are listed in Supplementary Table S2 | This study | N/A |
| **Software** | | |
| Mascot v.2.2.04 | MatrixScience | N/A |
| Scaffold v.5.3.3 | ProteomeSoftware | N/A |
| FragPipe v.17.1 and v. 20.0 | https://github.com/Nesvilab/FragPipe | N/A |
| Alphafold3 | https://alphafoldserver.com | N/A |
| Xcalibur v.3.0 | ThermoFischerScientific | N/A |
| Glob Quant | doi/10.1021/acs.jproteome.1c00596 | N/A |
| **Other** | | |
| TGX™ Precast Gels | Bio-Rad Laboratories | 4561096 |
| Petri dishes, 145x20mm | Greiner Bio-One | 639165 |
| 384 well flat bottom microplates | Greiner Bio-One | 784900 |
| Mini-PROTEAN® Tetra Vertical Electrophoresis Cell, 2-gel | Bio-Rad Laboratories | 1658005 |
| IKA KS 260 basic orbital shaker | IKA | Z341827 |
| Thermomixer Comfort | Eppendorf | |
| Incubator-Thermostat | Memmert | N/A |
| Cellstar tubes | Greiner | N188 271-N |
| Protein LowBind Tubes 1.5ml | Eppendorf | 022431081 |
| His-Trap FF Crude column | GE Healthcare | 17-5247-01 |
| Typhoon FLA9500 Fluorescence imager | Amersham | N/A |

| | | |
|---|---|---|
| Amersham™ Protran® Western-Blotting-Membranen, Nitrozow to referellulose | Sigma-Aldrich | GE10600002 |
| Semi-Phor TE70 Semi Dry transfer unit | Hoefer | TE70XP |
| Odyssey imager | LI-COR Bioscience | N/A |
| LUNA-II™ Automated Cell Counter | Logos Biosystems | L40001 |
| LM20 Microfluidizer® | Microfluidics | N/A |
| Q Exactive Orbitrap HF mass spectrometer | ThermoFischer Scientific | N/A |
| LTQ Orbitrap Velos mass spectrometer | ThermoFischer Scientific | N/A |
| Ultimate 3000 RSLnano System | ThermoFischer Scientific | N/A |
| µPAC™ HPLC column | ThermoFischer Scientific | COL-NANO050G1B |
| Acclam™ PepMap™ 100 75 µm x 2cm trapping column | ThermoFischer Scientific | 164750 |

### 2. Preparation of chimeric protein standards and model proteins

*2.1 Protein expression*

Model proteins and chimeric protein standards (Suppl Tables S1 and S2) were expressed in bacterial and insect cells at the Protein Biochemistry facility of the MPI-CBG.
Chimeric protein standards qFP-8 and FUGIS were expressed and $^{13}C_6{}^{15}N_4$-Arg and $^{13}C_6$ Lys metabolically labelled in *E.coli* cells as described in(*1, 2*) .
FPs were expressed in *E.coli* BL21(DE3) strain from pET11 backbone as HIS6 fusions.
Cells were cultivated in LB medium supplemented with 50 µg/ml of kanamycin at 37°C to $OD_{600}$=0.3 and induced with 0.2 mM IPTG for 2 hours at 37°C. Cells were then pelleted from 1 ml of cell culture ($OD_{600}$=3) by centrifugation at 500 g for 5 minutes, washed with 1x PBS and stored frozen at -70 °C.
FP-fusion proteins were expressed in *Trichoplusnia ni* cells. Shuttle constructs based on the pOEM1 backbone and ORFs encoding for fusion of proteins to fluorophores, and the corresponding baculoviruses were prepared as described(*3*) . *Trichoplusnia ni* cells at a concentration of $1\times10^6$ cells/mL were infected with 1% (v/v) baculovirus and cultivated in suspension for 72 h at 27°C in ESF 921 serum-free medium. Cells were then pelleted from 1 ml of cell culture (1 M cells/ml) by centrifugation at 500 g for 5 minutes, washed with 1x PBS and stored frozen at -70 °C.
HeLa cells stably expressing eGFP were provided by TDS Facility (Technology Development Studio), MPI-CBG; HCT116 cells stably expressing Venus and TagRFP – by Dr. Mihail Sarov, MPI-CBG. Briefly, HeLa Kyoto cells stably expressing eGFP were generated by transfection with ORF coding eGFP (QKFJN) cloned to pEGFP-C1 expression vectors. HCT116 cells stably expressing Venus or TagRFP were generated by Crispr Cas9 mediated homology directed repair (HDR) using RNP targeting the AAVS1 locus and repair template comprising traffic light reporter with both FPs. The cells were cultured on DMEM supplemented with 10% FBS and 100 µg/ml Geneticin, then washed with 1xPBS and incubated with 2 ml of Trypsin for 5 minutes at 37C degrees. Cell suspension was then transferred to a 15 ml Cellstar tubes, added 13 ml of serum containing medium to inactivate the trypsin and centrifuged at 138 g for 5 minutes. Cells in suspension were counted using LUNA-II Automated Cell Counter. The supernatant was discarded, cells washed twice and re-suspended in PBS.

*2.2. Cell lysis*

*E. coli* and *T.ni* cells expressing recombinant model proteins were pelleted as in #2.1 and lysed in 200 µl of 1% SDS acqueous solution containing 100x Halt™ Protease Inhibitors Cocktail by subsequent passing 10 times through a syringe and repeatable vortex. The samples were then centrifuged for 15 min at 14000g, supernatant collected, aliquoted and stored at -20°C.

0.5Mio HeLa and HCT116 cell stably transfected with FPs were re-suspended in 100 µl of 1x PBS buffer containing 0.2% NP40 and 100x Halt$^{tm}$ Protease Inhibitors Cocktail and processed as described above.

*2.3. SDS gel electrophoresis*

Sample aliquots (lysates of cells expressing model proteins or chimeric protein standards, cell-free extracts expressing mScarlet tagged G3P1 or its mutant, or of the reference protein BSA) were mixed with Laemmli Sample Buffer for SDS-PAGE supplemented with 0.1% of 2-mercaptoethanol (1:1 v/v) and separated on Mini-PROTEAN® Tetra Vertical Electrophoresis Cell using TGX$^{TM}$ Precast Gels according to the manufacturer protocol. An aliquot of Molecular Weight standards was loaded onto the first sample well. No reducing agents or heating were applied to the samples prepared for fluorescence assay. Gels for following-up mass spectrometric analyses were visualized by Coomassie staining as following: after a quick rinse with distilled water, the gel was incubated in staining solution (methanol : water : acetic acid = 5:4:1 v/v/v and Coomassie RGB 0.2%) in fresh single-use Petri dish for 2h at RT on a shaking platform. Then the solution was discarded, the gel de-stained by two changes of destaining mix (methanol : water : acetic acid = 45:45:2 v/v/v) for 2h and left overnight in water on an orbital shaker.

*2.4 In-gel fluorescence imaging and Western Blotting for SDS-PAGE-separated model FP-fusions*

Fluorescence in-gel imaging of FP-fusions following SDS-PAGE separation was performed on Typhoon FLA9500 Fluorescence imager. For FP-fusions ##A-I, in-gel fluorescence imaging and Western blotting were performed consecutively on the same gel. Aliquots of lysates of insect cells expressing fusions ##A-I were separated by SDS PAGE under non-reducing conditions and without heating. The volumes of aliquots were adjusted to fluorescence properties of the corresponding FPs and expression level of their fusions. Fluorescent signal was recorded using non-fixed wet fresh gel on Cy2, Cy3 and Cy5 channels using Typhoon FLA 9500 control software accordingly to the manufacturer protocol; Cy2 Laser 473nm BPB1 (530DF20), Cy3 Laser 532nm BPG1 (570DF20), Cy5 Laser 635nm LPR (665LP). The gel was then blotted onto 0.45 µm Western-Blotting membrane using SemiPhor semi-dry transfer unit for 1 h at 130 mA. The blot was blocked with 5% (w/v) nonfat dry milk and incubated overnight with the anti-HRV13 3C rabbit polyclonal antibodies (1:1000). After 3× washing with 0.1% Tween-20/PBS, the blot was incubated with IRDye800 Goat anti rabbit antibodies (1:1000). Bands were detected on Odyssey imaging system.

### 3. Sample preparation for mass spectrometric analysis

*3.1 Sample preparation for peptide mapping of FPs and FP-fusions:*

Aliquots of lysates of *E.coli* cells expressing FPs or of *T.ni* cells expressing FP-fusions (10 µl of stock, section 2.2) were separated by SDS-PAGE, the corresponding gel region *in-gel* digested with trypsin and analysed by LC-MS/MS.

*3.2 Sample preparation for ms-based absolute quantification of FP-fusions using chimeric standard proteins qFP-8 and FUGIS*

Aliquots of lysates of cells expressing FP-fusions (equivalent of 0.1µl of stock), qFP-8 (equivalent of 1µl of stock, section 2.2), FUGIS and 1 pmol of reference BSA were separated by SDS-PAGE and their Coomassie stained gel band excised. The position of the full-length FP-fusion band was determined as in #2.4. To correlate the intensities of peptide peaks of the corresponding FPs with their isotopically labelled proxies, FP-fusions and standards were first analysed separately. Their gel bands were individually *in-gel* digested with trypsin, analysed by LC-MS/MS and the intensities of the corresponding peptide peaks are compared. Then the adjusted accordingly volumes of cell lysates were separated by SDS-PAGE for quantitative experiment. The bands of FP-fusion, qFP-8, FUGIS and of 1pmol of reference BSA were excised, crashed with scalpel and mixed together in 1.5ml protein low-bind tubes. The protein mixture was *in-gel* reduced with DTT, alkylated with iodacetamide (IAA) and digested with trypsin overnight (ca 12h) at 37ºC as described (*4*). The resulting peptide mixture was extracted by two changes of 5% formic acid (FA) and acetonitrile at 37ºC in a thermomixer at 600 rpm, extracts pulled together and dried down in a vacuum centrifuge. Peptide pellets were dissolved in 50 or 100 µl of 5% FA and 5 µl of peptide mixture were taken for LC-MS/MS analysis. Each experiment was performed in four replicates.

*3.3 Sample preparation for ms-detection of chromophore-containing peptides*

Aliquots of lysates of *E.coli* cell expressing recombinant FPs were separated by 1D SDS PAGE (sections ##2.2, 2.3), the FP-corresponding gel regions were excised and digested *in-gel* overnight with trypsin or Asp-N protease. Resulting peptides were extracted as in #3.1.

For dsRed express and EGFP, proteins were processed from cell lysates of *E.coli* cells expressing recombinant dsRed-express and EGFP. Proteins were reduced with DTT and alkylated with IAA in an aliquot of cell lysate and precipitated as following: 10 µl of lysates were incubated with 90 µl of isopropanol for 10 min at 80ºC and spun down for 7 min at 13000g. The supernatant was discarded, fresh portion of 100 µl isopropanol added to the pellet and the procedure repeated. Then the pellet was dried in a vacuum centrifuge for 5 min. 0.5 µg Trypsin/Lys C mix was added directly on top of the pellet and digested overnight at 37ºC in a shaker. The resulting peptides were dried down and dissolved prior mass spectrometric analysis in 50 µl of 5% formic acid.

*3.4 Sample preparation for ms-based absolute quantification of FP expression in stably transfected cells*

20µl aliquots of the lysate of HeLa and HCT116 cells stably transfected with FPs, or 2µl of the *E.coli* lysate expressing FPs, were separated by SDS-PAGE, gel region corresponding to the FP excised and co-digested *in-gel* with qFP-8 standard and reference BSA as in #3.1. An aliquot of resulting peptide mix equivalent to 5K or 10K cells was then taken for analysis by mass spectrometry, or further diluted for the experiment in Suppl.Figure S5. Each experiment was performed in four replicates.

*3.5 Sample preparation for ms-based quantification of protein expression kinetics in cell free systems*

10µl aliquots of cell-free extracts expressing model proteins withdrawn at the fixed time points were mixed with the equal volume of Laemmli SDS sample loading buffer (no reducing agent added), separated by 1D SDS-PAGE and visualised by Coomassie. Gel regions of mScarlet tagged G3BP1 variants were excised and in-gel digested overnight by trypsin with no reduction and alkylation steps. The standard was prepared separately by co-digesting x20 amount of SDS gel-separated BSA and qFP-8. The resulting peptide mix was extracted as in #3.1. Quality and stability of the digested standard was monitored by LC-MS/MS comparing normalized intensities of the reference BSA peptides. The standard was then dissolved in 50µl of aquatic 5% formic acid and 1µl was spiked into samples prior mass spectrometric analysis (*5*). Each experiment was performed in four replicates.

**4. Mass spectrometric analysis and data processing**

*4.1 LC-MS/MS analysis*

LC-MS/MS analysis was performed on a UltiMate$^{tm}$ 3000 RSLCnano System interfaced on-line to a Q Exactive HF hybrid Quadrupole Orbitrap mass spectrometer or on a LTQ Orbitrap Velos mass spectrometer (for $MS^3$ experiments). The nano-LC system was equipped with Acclam$^{tm}$ PepMap$^{tm}$ 100 75 µm x 2cm trapping column and 50cm µPAC$^{tm}$ analytical column. Peptides were separated using 60, 80 or 120 min linear gradient (Table); solvent A - aqueous solution of 0.1% FA, solvent B - 0.1% FA in acetonitrile. For absolute quantification of FP-fusions and method validation experiments spectra were acquired in DDA (Data Dependent Acquisition) mode using Top20 method, precursor *m/z* range was 350 to 1600 Th; target mass resolution $Rs_{m/z\ 200}$ of 120000 (Full Width of Half Maximum, FWHM) and of 15000 (FWHM) for precursor and fragments respectively; dynamic exclusion time was set on 15s. The lock-mass function was set to recalibrate MS1 scans using the background ion of $(Si(CH3)_2O)_6$ at *m/z* 445.1200. For the protein expression kinetics experiments, we employed data-dependent acquisition (DDA) with inclusion list of m/z of precursor of mScarlet peptide proxies. Spectra were acquired in profile mode with $Rs_{m/z\ 200}$ 240000 (for MS) and 120000 (for MS/MS) and isolation window of 8 Th. HCD fragmentation was trigged by the precursor mass of the isotopically labelled peptide proxies. The amount of material loaded on the column was 200 to 400 fmoles. All analyses were performed in four replicates.

$MS^3$ experiment for mScarlet-I peptide GGPLPFSWDILSPQMYGSR comprising mature chromophore tri-peptide was performed on a LTQ Orbitrap Velos mass spectrometer. Peptides were separated using 80 min linear gradient. Doubly-charged precursor ion with *m/z* 1125.0280 was detected at $Rs_{m/z\ 200}$ 60000; fragmentation for the precursor and its singly-charged fragment with m/z 490.3300 was acquired under the following settings: isolation window 3Da, normalized collision energy 35, CID mode.

To avoid carryover, 2-3 blank runs were performed after each sample analysis, the last blank was recorded and also search against customized database.

Instrument performance was monitored using QCloud quality control system: https://qcloud2.crg.eu (ref.(*6*)).

Chromatographic gradients

| | 60 min gradient | | | 80 min gradient | | | 120 min gradient | | |
|---|---|---|---|---|---|---|---|---|---|
| step | time, min | flow, µl/min | B, % | time, min | flow, µl/min | B, % | time, min | flow, µl/min | B, % |
| 1 | 0 | 0.5 | 0 | 0 | 0.4 | 0 | 0 | 0.4 | 0 |
| 2 | 5 | 0.5 | 2 | 45 | 0.4 | 30 | 45 | 0.4 | 30 |
| 3 | 45 | 0.5 | 30 | 58 | 0.4 | 50 | 58 | 0.4 | 50 |
| 4 | 49 | 0.5 | 100 | 60 | 0.4 | 100 | 60 | 0.4 | 100 |
| 5 | 50 | 0.5 | 100 | 65 | 0.4 | 100 | 65 | 0.4 | 100 |
| 6 | 50 | 0.5 | 0 | 65 | 0.4 | 0 | 65 | 0.4 | 0 |
| 7 | 60 | 0.5 | 0 | 80 | 0.4 | 0 | 80 | 0.4 | 0 |

*4.2 MS data processing and absolute quantification*

For peptide mapping of FPs and FP-fusions, acquired spectra were matched by Mascot software against their sequences with the mass accuracy 5ppm and 0.05Da for precursor and fragments, respectively. Posttranslational modifications with the masses -20.026215 Da and -22.041865 Da corresponding to the loss of [-6H-O] and [-4H-O] respectively, were used for matching peptides comprising cyclic chromophore forms. Fragmentation spectra were visualized in Scaffold software.

For quantitative experiments, acquired spectra were processed by FragPipe software suit v.17.1 (*7*) under the following settings: mass tolerance 5 ppm and 0.05 Da for precursor and fragments, respectively; de-isotoping option – on; enzyme – trypsin; allowed_missed_cleavage

– 2; digest_mass_range 300 – 5000 Da; peptide_length 5 – 50 amino acids; variable modifications: 01 = 15.994900 M 3, 07 = 6.020129 KR 2, 09 = 10.008269 R 2, 10 = 31.989829 M 1, fixed modification 57.02146 C; searched for b-, y-, and a- ions. Spectra were matched against customized database comprising sequences of *E.coli* proteins (SwissProt protein collection), chimeric standards, FP-fusions and fluorescent proteins used in this study, and decoy database.

Molar abundances of model FP-fusions or of FPs expressed in stably transfected cells were quantified using qFP8 chimeric protein standard as described (*1*) (**Fig 1B**, Supplementary Fig S2). Briefly, non-normalized extracted ion chromatograms (XIC) of precursors of peptide proxies were extracted from ionquant-1.7.17 output (a part of FragPipe v.17.1) and sorted by in-house developed scripts. Then the amount of chimeric protein standard was related to the known amount of the reference protein BSA using areas of XICs peaks of reference peptide. Next, areas of XIC peaks of precursor *m/z* of native FP peptides were compared to the areas of the corresponding isotopically labelled peptides produced by digestion of the chimeric protein. Each peptide was quantified independently; to calculate the molar abundance of proteins the molar abundances of corresponding peptides were averaged.

Projected molar abundance and concentration of FPs in a single cell was re-calculated using counted cell number and radius measured for HeLA and HCT116 cells as 7.5 µm and 5 µm respectively; the reference values for *E.coli* cells were taken from the literature (https://bionumbers.hms.harvard.edu; https://tipbiosystems.com/wp-content/uploads/2020/05/AN102-E.coli-Cell-Count_2019_04_25.pdf).

Absolute quantification of FP-fusions with FUGIS standard was carried out by GlobQuant software as described (*2*) using peptide intensities of combined ion output provided by MSFragger v. 3.8 (a part of Fragpipe v. 20.0). The amount of FUGIS standard was related to the reference protein BSA. Best3 peptides whose XIC areas were having coefficient of variation (CV) <30% were selected from four most abundant precursors in FP-fusion digests. Where specified, Best2 peptides were taken. For quantification of FP-fusions relying to their fused FPs, Best3 peptides were selected from the peptides matching FP part of the sequence. Molar abundances of FP-fusions was calculated by comparing of averaged intensities of Best3 peptides with the median of FUGIS peptides.

Absolute quantification of FP-fusions by Top3-Hi method was performed as described by *(40)* using areas of XIC of their three most abundant peptide peaks and of qFP-8 as reference protein.

To quantify the fraction of mScarlet comprising maturated cyclic chromophore, XICs for the precursors of chromophore-containing peptide GGPLPFSWDILSPQMYGSR with linear and cyclic MYG-tripeptide were manually extracted in XCalibur Qual Browser. XIC of the cyclic form was then normalized to their sum and related to the total amount of the corresponding mScarlet fusion.

### 5. Plasmid preparation for cell-free expression

#### *5.1 Plasmid Design*

A pOCC plasmid backbone was provided by the Protein Biochemistry facility of the MPI-CBG, for expression in the TnT extract. The pOCC vectors were engineered to include AscI (5'-GG/CGCGCC-3') and NotI (5'-GC/GGCCGC-3') restriction sites downstream of the T7 promoter for modular restriction cloning with various inserts. Each vector contains an antibiotic resistance gene (AmpR) and an *E. coli*-specific origin of replication (ORI) for selection and plasmid amplification, respectively. The vector contained the sequence for the fluorescent protein reporter mScarlet-I. The wild type G3BP1 and the deletion mutant G3BP1(delE1/E2)

were obtained from pOCC189-G3BP1(WT) L515 and pOCC189-G3BP1(ΔE1ΔE2) L-667 (*8*). Restriction cloning of the G3BP1 gene using the NotI-HF and AscI enzymes and ligation with T4 DNA ligase (both from NEB) resulted in the creation of plasmids G3BP1-mScarlet-I and G3BP1(delE1/E2)-mScarlet-I (Figure 1(a)).
For the constructs compatible with the PURE system, the wild type and mutant G3BP1 coding sequences were codon optimized for *E.coli* and synthesized by Twist Biosciences (www.twistbioscience.com). The oligomers were amplified with SSO-001 (TCCGCTGCTGGTTCTGGCGCGGCCGCGATGGTTATGGAAAAACCTTCCCCG) and SSO-002 (CTTCTATTACGGCGCGCCTTGACGTGGAGCCAGGCC) primers to make overhang for Gibson assembly. The oligomer was cloned into a modified pEXP5NT backbone, bearing mScarlet-I (Figure 1(b)) by Gibson assembly. All four constructs were confirmed by sequencing.

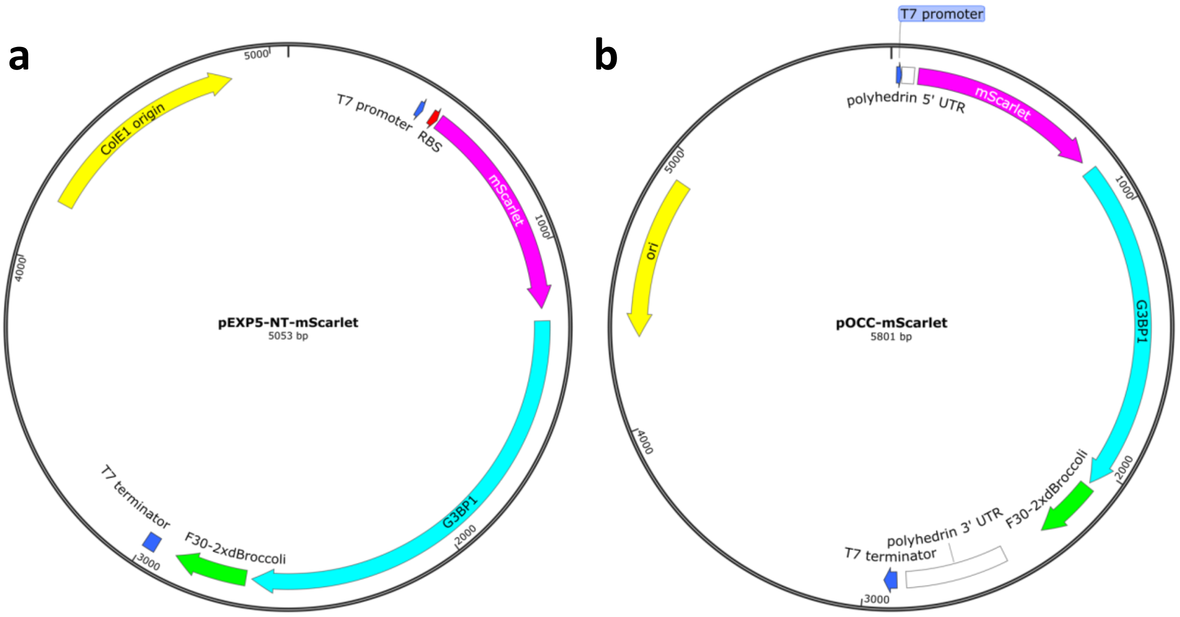


Plasmid construct maps for the a) pEXP5-NT and b) pOCC vectors used for insect cell and bacterial system expression coding for G3BP1 with an N-terminal mScarlet-I tag. The Broccoli aptamer came with the construct for mRNA quantification and was not relevant to this project.

List of primers used in cell-free expression experiment

| Primer | Sequence (5'->3') | ? |
|---|---|---|
| SSO-001 | TCCGCTGCTGGTTCTGGCGCGGCCGCGATGGTTATGGAAAAACCTTCCCCG | 1 |
| SSO-002 | CTTCTATTACGGCGCGCCTTGACGTGGAGCCAGGCC | 1 |
| SSO-007 | GCGGCCGCGCCAGAACCAGCAGCGGAGCCAGCAGATCCCTTGTACAGTTCATCCATACCA | 2 |
| Tphi fwd | CGAAAGGAAGCTGAGTTG | 2 |
| SSO-004 2xBroc AscI F | AAGGCGCGCCGTAATAGAAGTTGCCATGTGTATGTGGGAGAC | 3 |
| SSO-005 2xBroc EcoRI R | GCAGCCAACTCAGCTTCCTTTCGGGAATTCTTAGCAGCCGTTGCCATG | 3 |
| SSO-006 2xBroc seq R | CGTCTCCCACATACACATGGCAA | Seq |
| SSO-009 mScarlet-Ec seq F | TATGAGCGCAGCGAAGGCCGTC | seq |

<<@Archie: what is the header of the last column?>>

*5.2 Restriction Digestion-Ligation of DNA*

For each sample, 1-5 µg of total DNA was digested in a total volume of 50 µl containing 5 µl of CutSmart buffer (10x). 1 µl of restriction enzyme per 1000 ng of DNA (maximum 2.5 µl per sample) was added. Digests were incubated for 20-60 minutes at 37°C and heat-inactivated for 20 minutes at 65°C if not used immediately for gel electrophoresis.
Ligation of digested DNA fragments was performed in a 20 µl total volume containing 1 µl (400 units) T4 ligase and 2 µl T4 Ligation buffer. A total weight of 50-80 ng of insert with a 2-3-fold molar excess over the vector was used. Ligation was either performed at room temperature for 20 minutes or at 16°C overnight, followed by heat inactivation of the ligase at 65°C for 10 minutes according to the manufacturer's recommendations.

*5.3 Agarose Gel Electrophoresis*
Agarose gel electrophoresis was conducted for quality verification and fragment separation of DNA and RNA using 0.7-1.0% agarose gels at 100-125 V for 1 hour with 1x TAE buffer as both the running buffer and for casting the gels. For this, 0.28–1 g of agarose was melted in 40-100 ml of 1x TAE buffer by microwaving and poured into the casting chamber (4.5–10 cm x 10 cm). Gels contained 0.01% "GelRed Nucleic Acid Stain 10,000X Water" and were examined under UV light at 312 nm in a photo chamber. The DNA samples for cell-free expression were prepared for loading by adding 1 part of "Gel Loading Dye Purple" to 6 parts of sample. For size comparison, "Quick-Load© 1 kb DNA Ladder" was loaded into the gel and run in parallel to the samples.

*5.4 DNA Isolation and Gel Extraction*
Plasmid DNA isolation was performed using the QIAprep Spin Miniprep Kit and Plasmid Maxi Kit, following the manufacturer's protocol. Bacteria were cultured overnight in 5-9 ml of antibiotic-containing media (50 µg/ml Kanamycin or 100 µg/ml Ampicillin) for miniprep, and in 500 ml for maxiprep.
Gel extraction of DNA fragments was performed using the QiaQuick Gel Extraction Kit, following the manufacturer's protocol.

*5.5 Transformation of DNA into Competent Cells for storage and amplification*
1-5 µl of plasmid DNA was added to 50 µl NEB® 5-alpha chemically competent cells in an Eppendorf tube and incubated for 30 minutes on ice. The cells were then heat-shocked at 42°C for 30 seconds and immediately placed on ice for 2 minutes. 950 µl of SOC media was added to the cells, which were then incubated at 37°C and 300 rpm in a thermo shaker for 1 hour. 100 µl of cells were plated on a pre-warmed LB agar plate containing the appropriate antibiotic (Kanamycin or Ampicillin) and incubated at 37°C overnight. Single colonies were picked and transferred to a fresh agar plate for storage. Additionally, 5-8 ml of antibiotic (50 µg/ml Kanamycin or 100 µg/ml Ampicillin)-containing LB media in a 15 ml cultivation tube was inoculated with single isolated colonies for plasmid amplification. Plates and tubes were incubated at 37°C overnight, with the tubes shaken at 220 rpm. Water was used as a negative control instead of plasmid DNA.

### 6. Quantitative Fluorescence Measurement for cell-free expressed G3BP1 tagged with mScarlet

*6.1 Preparation of mScarlet-I standard*
*E.coli* T7express strain, bearing pRARE helper plasmid, was used for the expression of pure mScarlet-I tagged with $His_6$ for chromatographic isolation using a His-Trap FF Crude column. The strain was transformed with the expression plasmid and a helper plasmid, and the

transformants were selected on LB agar plates supplemented with 90 μg/ml Kanamycin and 17 μg/ml Chloramphenicol.

A 250 ml Terrific Broth (TB) culture was inoculated with a single colony from the selection plate and incubated overnight at 37°C with agitation at 220 rpm. This pre-culture was subsequently diluted 1:1000 in 1 liter of TB medium containing 90 μg/ml Kanamycin and 17 μg/ml Chloramphenicol and incubated at 37°C with shaking at 220 rpm until an optical density at 600 nm of 0.5 was reached. The incubation temperature was then reduced to 18°C for 1 hour at 220 rpm. Protein expression was induced by the addition of 0.2 mM Isopropyl β-D-1-thiogalactopyranoside (IPTG) from a 20 mM stock solution in water, and the culture was incubated overnight at 18°C with shaking at 220 rpm.

The cells were harvested by centrifugation at 2,000 rpm and 4°C for 15 minutes. The resulting pellet was resuspended in lysis buffer (2x PBS, 20 mM Imidazole, 1 mM DTT, pH 7.4) and lysed using an LM20 Microfluidizer at 15,000 bar on ice in two consecutive runs. The lysate was clarified by centrifugation at 16,000 rpm and 4°C for 30 minutes, followed by filtration through a 0.45 μm sterile filter. The clarified supernatant was loaded onto two equilibrated 5 ml HisTrap FF crude columns at a flow rate of 2.5 ml/min.

The columns were washed sequentially at a flow rate of 5 ml/min with 10 column volumes of low-salt wash buffer (2x PBS, 20 mM Imidazole, 1 mM DTT), 10 CV of high-salt wash buffer (6x PBS, 1 mM DTT), and again 10 column volumes of the low-salt wash buffer. Elution of the bound protein was performed using an elution buffer (2x PBS, 250 mM Imidazole, 1 mM DTT) and collected in 2 ml fractions. The eluted fractions were analyzed by SDS-PAGE followed by Coomassie staining, and fractions containing pure mScarlet-I were pooled based on relative concentration and purity.

The pooled protein solution was dialyzed against 2x PBS with 5% glycerol at 4°C overnight, concentrated, aliquoted and subsequently stored at -80°C following flash freezing in liquid nitrogen. A small volume of the highly concentrated stock was then diluted in PURE extract to perform the calibration.

*6.2 Fluorescence assay*

The expression of mScarlet-I tagged G3BP1 in PURE was tracked on a TECAN Spark 20M plate reading fluorimeter. The protein concentration was tracked with the mScarlet-I fluorescence with excitation/emission at 569/594 nm with identical bandwidth and gain as before.

The instrument was pre-incubated at 37 °C. In a 10 μl aliquot, 4 μl of the PURE A component and 3 μl of the PURE B component was added with 10 ng/μL plasmid, loaded in triplicate onto the 384 well plate and sealed with the adhesive cover. The reactions were run for 4 hours each.

*6.3 Calibration of fluorescence data*

Fluorescence units for the reactions were converted to concentration with calibration curves obtained from purified mScarlet-I protein. The protein was aliquoted at a concentration range that covered the highest fluorescent read out for expressed fusion proteins in the PURE reaction systems. All aliquots had the same concentration of PURE as used in the experiments with mScarlet-I tagged G3BP1. The calibration curve was fit to a straight line, using MATLAB.

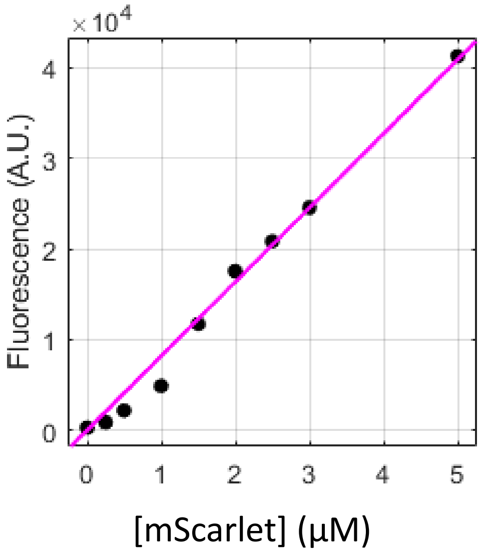

Calibration curves for mScarlet-I fluorescence in PURE system.

**7. Protocols for generating 3D structures for G(WT)-mS and G(mut)-mS**

To create 3D structures of the fusion proteins analyzed in this study, Protein Data Bank (PDB) files were generated using AlphaFold3 by submitting the sequences of both the wild-type (WT) and mutant G3BP1–mScarlet-I constructs to https://alphafoldserver.com. The resulting structures were loaded into PyMOL and verified by structural alignment with reference PDB files for mScarlet-I (PDB ID: 5LK4) obtained from the RCSB Protein Data Bank (https://rcsb.org), and with AlphaFold-predicted G3BP1 structures from the AlphaFold database. For intrinsically disordered protein (IDP) regions, one of five randomly suggested conformations from AlphaFold3 was selected. Since the folded fluorescent protein aligned well and the disordered section is expected to not have a structure, no energy minimization was done.

The verified structures were then visually edited using PyMOL command-line instructions. To define specific regions—such as separating the IDP domain from the fluorescent protein (FP) region, or highlighting acidic regions—the following selection syntax was used:

*select <region_name>, resi <start_residue>-<end_residue>*

Color assignments were made using:

*color <color_name>, <region_name>*

To highlight specific residues, such as all glutamic acids within a region, the following selection command was applied:

*select glutamic_acids, resn GLU and resi <start_residue>-<end_residue>*

These residues were visualized as red sticks using:

*hide everything, glutamic_acids*
*show sticks, glutamic_acids*
*color red, glutamic_acids*

For electrostatic surface rendering, the APBS plugin was installed if not already present. It was accessed via *Plugins → APBS Electrostatics*, and a PQR file was generated (saved as run01). The default color bar was removed using:

*for ramp in cmd.get_names_of_type('ramp'): cmd.disable(ramp)*

Electrostatic surface transparency was adjusted to 0.3 using:

*set transparency, 0.3*

(Note: This was applied after selecting the *run01* object and not other structure representations.)

Phosphorylated structures were created using ChimeraX by loading the PDB files, removing hydrogen atoms from the hydroxyl groups of serine residues undergoing phosphorylation, and manually adding phosphate groups via the *Build Structure* tool under *Structure Editing* in the *Tools* menu. The structures were for visualization purpose only and therefore not energy minimised.

All structural representations were exported as PNG images.

### 8. Kinetic Model

#### *8.1 Original model*

The fluorescence and mass spectrometric progress curves were fit to the analytical solutions of three coupled differential equations. Protein expression in cell-free expression systems can be simply described by the conversion of transcription and translation nutrient resource (TlR) to full length immature protein (P), a fractional yield (Y) of which converts to a fluorescent form ($P_m$) after maturation. The remaining fraction (1-Y) goes to an inactive non-fluorescent state ($P_i$) which is lost from the scope of our analysis. For simplicity we keep the rate constant for both generation of mature and inactive fractions to be $k_m$ and the overall protein production rate to be $k_t$. With this narrative, the pathways can be modelled thus:

$$TlR \overset{k_t}{\rightarrow} P$$

$$Y.P \overset{k_m}{\rightarrow} P_m$$

$$(1-Y).P \overset{k_m}{\rightarrow} P_i$$

The rate of nutrient consumption as a function of protein production rate $k_t$ can be modelled as first-order decay with an overall expression rate constant $k_t$:

$$\frac{d[TlR]}{dt} = -k_t.[TlR] \qquad (1)$$

The relationship between immature protein (P) production rate with maturation rate constant $k_m$ can also be described as a balance between production rate from using TlR and depletion rate into $P_m$ and $P_i$:

$$\frac{d[P]}{dt} = k_t.[TlR] - k_m.Y.[P] - k_m.(1-Y).[P] = k_t.[TlR] - k_m.[P] \tag{2}$$

Finally, the maturation rate from immature protein to fluorescent protein is simply a first-order growth depending on the concentration of fractional concentration of P that goes into the successful maturation pathway:

$$\frac{d[P_m]}{dt} = k_m.Y.[P] \tag{3a}$$

And

$$\frac{d[P_i]}{dt} = k_m.(1-Y).[P] \tag{3b}$$

We solve equation (1) by integrating both sides, with initial nutrient concentration $[TlR]_0$ at zero time:

$$\int_{[TlR]_0}^{[TlR]} \frac{d[TlR]}{[TlR]} = -k_t \int_0^t dt$$

or,

$$[TlR] = [TlR]_0.e^{-k_t.t} \tag{4}$$

In order to solve the immature protein production rate $\frac{d[P]}{dt}$ we first substitute the relation for [TlR] from equation (4) into equation (2):

$$\frac{d[P]}{dt} = k_t.[TlR] - k_m.[P]$$

or,

$$\frac{d[P]}{dt} = k_t.[TlR]_0.e^{-k_t.t} - k_m.[P]$$

To solve this, we have to take an indefinite integral both sides, which requires the following steps:

$$\frac{d[P]}{dt} + k_m.[P] = k_t.[TlR]_0.e^{-k_t.t}$$

Multiplying by $e^{k_m.t}$ on both sides:

$$e^{k_m.t}\frac{d[P]}{dt} + k_m.e^{k_m.t}.[P] = k_t.[TlR]_0.e^{-k_t.t}.e^{k_m.t}$$

The left-hand side can be written as the derivative of the product of $e^{k_m.t}$ and $[P]$

$$\frac{d([P].e^{k_m.t})}{dt} = k_t.[TlR]_0.e^{(k_m-k_t).t}$$

Now we integrate both sides:

$$\int d([P].e^{k_m.t}) = k_t.[TlR]_0 \int e^{(k_m-k_t).t}.dt$$

or,

$$[P].e^{k_m.t} = \frac{k_t.[TlR]_0.e^{(k_m-k_t).t}}{k_m-k_t} + constant$$

or,

$$[P] = constant.e^{-k_m.t} - \frac{k_t.[TlR]_0.e^{-k_t.t}}{k_t-k_m} \quad (5)$$

In order to get the value of the constant we set the boundary condition that the concentration of immature protein and zero time is zero:
$Given, [P]_{t=0} = 0$

$$0 = constant.e^{-k_m.0} - \frac{k_t.[TlR]_0.e^{-k_t.0}}{k_t-k_m}$$

or,

$$constant = \frac{k_t.[TlR]_0}{k_t-k_m}$$

Substituting the value for the constant back into equation (5)

$$\therefore [P] = \frac{k_t.[TlR]_0}{k_t-k_m}(e^{-k_m.t} - e^{-k_t.t}) \quad (6)$$

Solving for the mature protein production rate (3) by substituting (6) in (3):

$$\frac{d[P_m]}{dt} = k_m.Y.[P]$$

or,

$$\frac{d[P_m]}{dt} = k_m.Y.\frac{k_t.[TlR]_0}{k_t-k_m}(e^{-k_m.t} - e^{-k_t.t})$$

Solving by taking an indefinite integral on both sides:

$$\int d[P_m] = \frac{k_t.k_m.Y.[TlR]_0}{k_t-k_m}\left(\int e^{-k_m.t}.dt - \int e^{-k_t.t}.dt\right)$$

or,

$$[P_m] = \frac{k_t.k_m.Y.[TlR]_0}{k_t-k_m}\left(-\frac{e^{-k_m.t}}{k_m} + \frac{e^{-k_t.t}}{k_t}\right) + constant$$

Setting the boundary condition that the concentration of mature protein at zero time is zero, we get the value for the constant to be:

$$Given, [P_m]_{t=0} = 0; constant = -\frac{k_t.k_m.Y.[TlR]_0}{k_t-k_m}.\frac{k_m-k_t}{k_t.k_m}$$

$$= Y.[TlR]_0$$

$$\therefore [P_m] = \frac{k_t.k_m.Y.[TlR]_0}{k_t-k_m}\left(\frac{e^{-k_t.t}}{k_t} - \frac{e^{-k_m.t}}{k_m}\right) + Y.[TlR]_0 \quad (7)$$

Note that at infinite time:

$$[P_m]_{t=\infty} = \frac{k_t.k_m.Y.[TlR]_0}{k_t - k_m}\left(\frac{e^{-k_t.\infty}}{k_t} - \frac{e^{-k_m.\infty}}{k_m}\right) + Y.[TlR]_0 = Y.[TlR]_0$$

This implies that when the reaction reaches equilibrium only a fraction Y of the total nutrient resource is converted to mature protein $P_m$, in line with the observations.

The mass spectrometer observes total protein as well as mature fraction. Modelling the mature fraction data is exactly the same as the fluorescence data hence equation (7) applies. In order to obtain the analytical solution to the rate equation for total protein we consider that the total protein content at any time is a sum of the immature, mature and inactive protein content:

$$[P_T] = [P] + [P_m] + [P_i]$$

Which applies to their rates as well:

$$\frac{d[P_T]}{dt} = \frac{d[P]}{dt} + \frac{d[P_m]}{dt} + \frac{d[P_i]}{dt}$$

From equations (2), (3a) and (3b);

$$\frac{d[P_T]}{dt} = k_t.[TlR] - k_m.Y.[P] - k_m.(1 - Y).[P] + k_m.Y.[P] + k_m.(1 - Y).[P]$$

or,

$$\frac{d[P_T]}{dt} = k_t.[TlR]$$

Substituting in equation (4):

$$\frac{d[P_T]}{dt} = k_t.[TlR]_0.e^{-k_t.t}$$

Integrating both sides with boundary conditions $P_T = 0$ at $t = 0$:

$$\int_0^{[P_T]} d[P_T] = k_t.[TlR]_0.\int_0^t e^{-k_t.t}dt$$

We get the expression for total protein:

$$[P_T] = [TlR]_0(1 - e^{-k_t.t}) \qquad (8)$$

Which is used to fit the mass spectrometry data for total protein concentration over time.

Note that if we wish to model the transcription processes the following two differential equations need to be considered for an mRNA (R) production rate and a transcription resource (TsR) usage rate with transcription rate constant $k_r$:

$$\frac{d[R]}{dt} = k_r.[TsR].[DNA]$$

And,

$$\frac{d[TsR]}{dt} = -k_r.[TsR]$$

Which has the solution of the form same as equation (4).

$$[TsR] = [TsR]_0.e^{-k_r.t}$$

Which gives the original mRNA production rate as:

$$\frac{d[R]}{dt} = k_r.[TsR]_0.[DNA].e^{-k_r.t}$$

Which has the solution given there is no mRNA at zero time:

$$[R] = [TsR]_0.[DNA].(1 - e^{-k_t.t})$$

Let us consider all transcription processes reach a peak by t = $t_0$ where mRNA becomes constant:

$$[R]_{max} = [TsR]_0.[DNA].(1 - e^{-k_t.t_0})$$

The true rate equation for P should be, with all parameters, same as equation (2) except for TlR' which is translation resources without the mRNA:

$$\frac{d[P]}{dt} = k_t.[TlR'].[R] - k_m.[P]$$

Which, by the time translation kicks off around $t_0$, mRNA can be considered to be constant:

$$\frac{d[P]}{dt} = k_t.[TlR'].[R]_{max} - k_m.[P]$$

or,

$$\frac{d[P]}{dt} = k_t.[TlR'].[TsR]_0.[DNA].(1 - e^{-k_t.t_0}) - k_m.[P]$$

Which can be reduced to include the expression for the mRNA, rendered a constant, to get the original equation (2), with translation resource TlR that includes mRNA:

$$\frac{d[P]}{dt} = k_t.[TlR] - k_m.[P]$$

Note that the analytical form of $P_i$ is unimportant to the scope of this work and therefore was left unsolved.

*8.2. Extended Model*

Considering that the mature fraction of the protein expression product consistently showed faster kinetics than that of the fluorescent product, we explored an extended version of the original model. In this model, translation resources (TlR) progress to an immature state P via an intermediate precursor $P_0$, introduced to account for delays in the formation of mature and fluorescent fractions. Recall that this delay was accounted for by the time constant $t_0$ in the original model with the Heaviside function.

In the original model, the immature state transitions directly to the mature fluorescent state ($P_m$). Here, we introduce an intermediate state $P_c$ which represents a state where the chromophore is supposed to have formed and therefore detectable by mass spectrometry but not yet folded and therefore undetectable to the fluorimeter. This is then followed by a properly folded mature state $P_f$. This is represented by the following schematic with corresponding rate constants:

$$TlR \xrightarrow{k_{t1}} P_0 \xrightarrow{k_{t2}} P \xrightarrow{k_c} P_c \xrightarrow{k_f} P_f$$

Considering that the products detected by mass spectrometry and fluorescence are fractions of their respective precursors, we define the observed chromophore-formed product ($P_{co}$) as originating from a fraction X of its precursor P. Similarly, the observed fully folded product ($P_{fo}$) originates from a fraction Y of its precursor $P_c$. The remaining fractions, (1–X) of P and (1–Y) of $P_c$, lead to unobserved products ($P_{cu}$ and $P_{fu}$) and are not considered further. Notably, the observed chromophore-formed product ($P_{co}$) alone does not account for the entire observed fully folded product ($P_{fo}$), as the fluorescent signal is consistently higher than the mature chromophore signal. Thus, the model retains dependence on the total amount of chromophore-formed product ($P_c$) rather than solely on the observable fraction ($P_{co}$).

These pathways can be summarized thus:

$$X.P \overset{k_c}{\rightarrow} P_{co}$$

$$Y.P_c \overset{k_f}{\rightarrow} P_{fo}$$

$$(1-X).P \overset{k_c}{\rightarrow} P_{cu}$$

$$(1-Y).P_c \overset{k_f}{\rightarrow} P_{fu}$$

The differential equations describing these pathways are outlined below. The decay of translation resources (TlR) follows first-order kinetics, with the solution carried over from the original model:

$$\frac{d[TlR]}{dt} = -k_t.[TlR]$$

$$[TlR] = [TlR]_0.e^{-k_t.t}$$

P0 formation results from the balance between its supply from TlR and its depletion via conversion to P:

$$\frac{d[P_0]}{dt} = k_{t1}.[TlR]_0.e^{-k_t.t} - k_{t2}.[P_0]$$

P formation is governed by the supply from P0 and its loss through conversion to Pc:

$$\frac{d[P]}{dt} = k_{t2}.[P_0] - k_c.[P]$$

Pf formation depends solely on the conversion from Pc:

$$\frac{d[P_f]}{dt} = k_f.[P_c]$$

The formation rate of the observable chromophore-containing product ($P_{co}$) is determined by the conversion of a fraction X of P:

$$\frac{d[P_{co}]}{dt} = k_c.X.[P]$$

The rate of formation of the observable fully folded product ($P_{fo}$) is governed by the conversion of a fraction Y of $P_c$:

$$\frac{d[P_{fo}]}{dt} = k_f.Y.[P_c]$$

These differential equations were numerically fit to the data using MATLAB using the lsqnonlin function, wherein the total protein detected by mass spectrometry is fit to

$$[P_T] = [P_0] + [P] + [P_c] + [P_f]$$

And the mature fraction detected by mass spectrometry is fit to $[P_{co}]$ and the fluorescent fraction detected by fluorescence is to $[P_{fo}]$.

Compared to the original model, where the immature product P directly matures into the fluorescent product $P_m$ at a rate $k_m$:

$$P \overset{k_m}{\longrightarrow} P_m$$

the extended model separates maturation into two sequential steps: conversion of P into a chromophore-formed intermediate $P_c$ at rate $k_c$, followed by folding into the fully fluorescent product $P_f$ at rate $k_f$:

$$P \overset{k_c}{\rightarrow} P_c \overset{k_f}{\rightarrow} P_f$$

Thus, the effective time constant for the two-step maturation process in the extended model ($\tau_{eff}$) is equivalent to the single-step maturation time constant ($\tau_m$) in the original model:

$$\tau_{eff} = \tau_m = \tau_c + \tau_f$$

Accordingly, the corresponding rate constants—being inverses of the time constants for first-order reactions—are related as follows:

$$\frac{1}{k_{eff}} = \frac{1}{k_m} = \frac{1}{k_c} + \frac{1}{k_f}$$

The percentage difference between the effective rate constant ($k_{eff}$) and the original single-step rate constant ($k_m$) was calculated using:

$$Difference = \frac{|k_m - k_{eff}|}{k_m} \times 100\ \%$$

**9. Fitting and simulations**

All fits for the original model were performed on MATLAB using the fit function with a non-linear least squares method following the 'Trust-Region' algorithm. Equation (7) was used to fit the kinetic curves for total protein measured by mass spectrometry. A Heaviside step function was used to account for the delay due to transcription. The final fitting form was:

$$[P] = [TlR]_0\left(1 - e^{-k_t.(t-t_0)}\right).H(t - t_0) \qquad (9)$$

The $k_t$ value obtained from this fit was then substituted into equation (7) which was used to fit the kinetic curves for mature protein measured by fluorescence spectroscopy. A Heaviside step function was added for the same purpose as before. The final fitting form was:

$$[P_m] = \left(\frac{k_t.k_m.Y.[TlR]_0}{k_t - k_m}\left(\frac{e^{-k_t.(t-t_0)}}{k_t} - \frac{e^{-k_m.(t-t_0)}}{k_m}\right) + Y.[TlR]_0\right).H(t - t_0) \qquad (10)$$

Simulations of this model was done by filling in the obtained parameters to the analytical solutions and plotting.

For fits to the extended model, the differential equations were solved using the ode45 function in MATLAB and numerically fit to the data using the lsqnonlin function. Simulations of this model was done by solving the differential equations with ode45 using the parameters obtained by fitting, and plotting.
