## Supplementary Dataset for "Absolute Quantification of Fluorescent Protein Fusions by Proteomics"

### List of Supplementary Figures and Tables

**Figure S1.** Characterization of the chimeric protein standard qFP-8 by LC-MS

**Figure S2.** Workflow for absolute quantification of FPs and FP-fusions with qFP-8 chimeric protein standard

**Figure S3.** Characterization of FP-fusions ##A-I by Western Blot and Fluorescent Gel Imaging

**Figure S4**. Peptide mapping of FP-fusions by mass spectrometry

**Figure S5.** Absolute quantification of EGFP in stably transfected HeLa cells with qFP-8 standard

**Figure S6.** Fragmentation spectra of chromophore-containing peptides from red- and green-type FPs

**Figure S7.** Chromophore-containing peptides detected in red-type FP dsRed-express by mass spectrometry

**Figure S8**. Removal of glutamic acid residues from intrinsically disordered region changed formal net charge of the mScarlet-tagged protein G3BP1

**Figure S9**. Analysis of short products of expression

**Figure S10**. Detection of serine phosphorylation in G(WT)-mS expressed in TnT cell-free system by mass spectrometry

**Figure S11**. Extended model

**Figure S12.** Simulations of hidden variables using the original and extended models

**Table S1.** List of Fluorescent Proteins, Self-labelling Tags and their Fusions

**Table S2.** Peptide proxies included in qFP-8 chimeric standard protein

**Table S3.** MS-based approaches for absolute quantification of proteins using peptide references and spiked protein standards

**Table S4.** Amount of FPs-fusions ##A-J quantified using peptide proxies of the qFP-8 chimeric standard

**Table S5.** Examples of FP amounts in stably transfected cells quantified using qFP-8 standard

**Table S6.** Abundance of chromophore-containing peptides in red FP mScarlet and dsRed-express

**Table S7.** Phosphorylation status of the G3BP1 peptide SSSPAPADIAQTVQEDLR detected by mass spectrometry in G(WT)-mS and G(mut)-mS expressed in TnT and PURE cell free expression systems

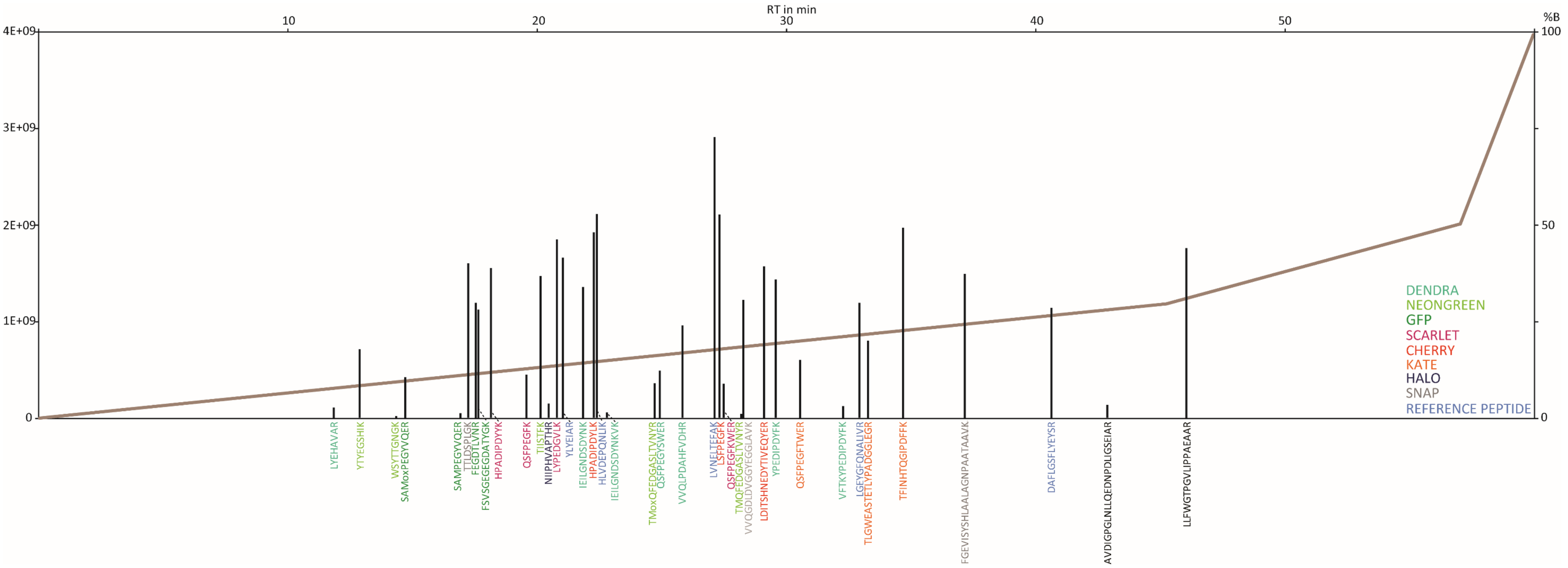


**Supplementary Figure S1.** Characterization of the chimeric protein standard qFP-8 by LC-MS.

150fmol of tryptic digest of the qFP-8 was injected. FP proxy peptides are colour-coded (see legend on the right-hand side). Vertical axis at the left-hand side shows the raw intensity of the peptide signal. The intensities of the +2/+3 charge states of the same peptide are summed. The intensity of Met-containing peptides is the sum of intensities native and mono-oxidized forms. The vertical axis at the right hand side shows % of acetonitrile (solvent B) in the chromatographic gradient; the gradient profile is shown as a continuous black line. The top horizontal axis shows the retention time.

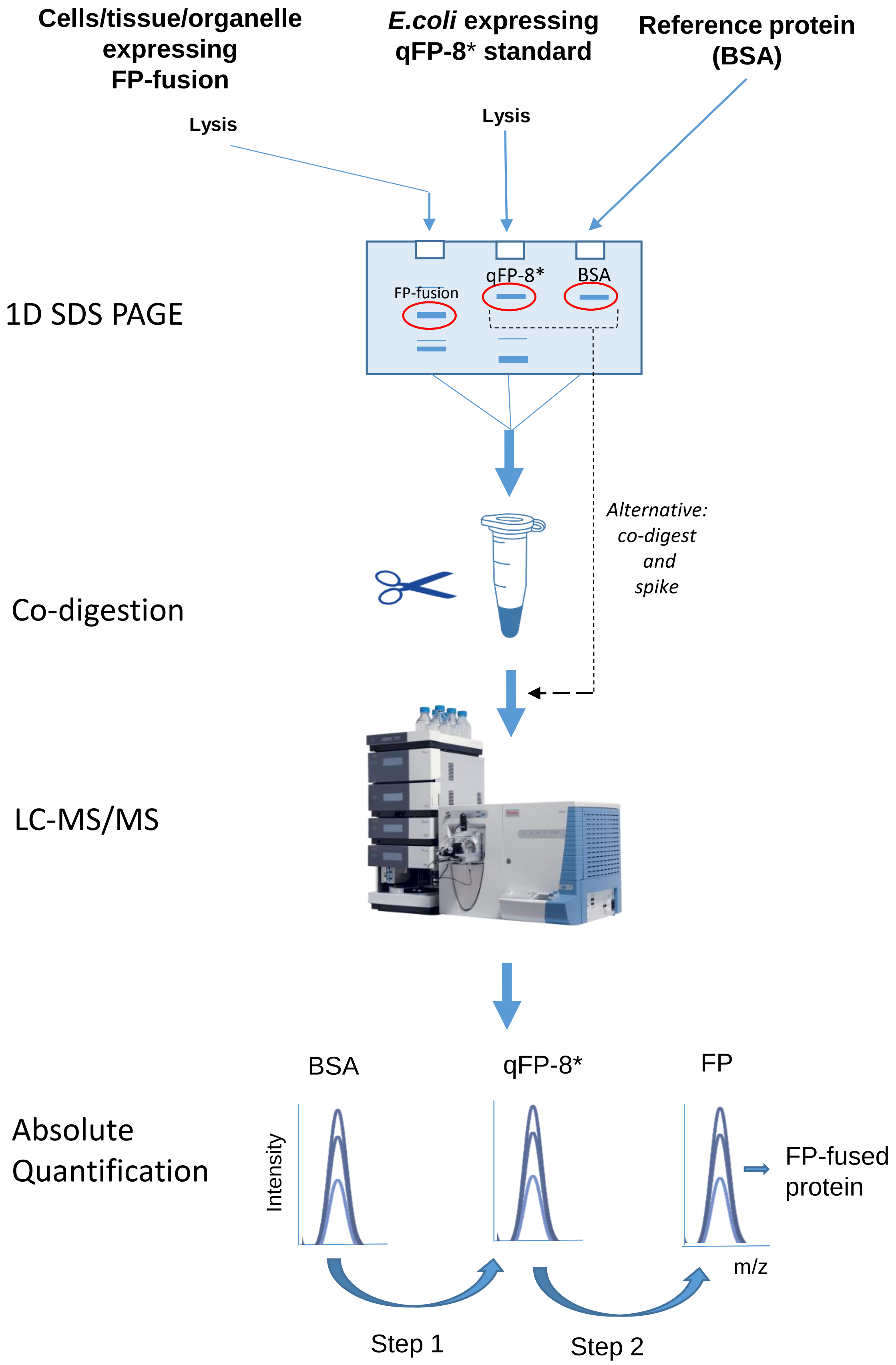


**Supplementary Figure S2.** Workflow for absolute quantification of FPs and FP-fusions with qFP-8 chimeric protein standard. "*" stays for $^{13}C_6{}^{15}N_4$-Arg and $^{13}C_6$ – Lys in metabolically labelled proteins. Dashed arrow: alternative sample preparation step in which qFP-8 and BSA are digested separately and spiked into the protein digest before LC-MS/MS analysis. This helps in scaling up the experiments.

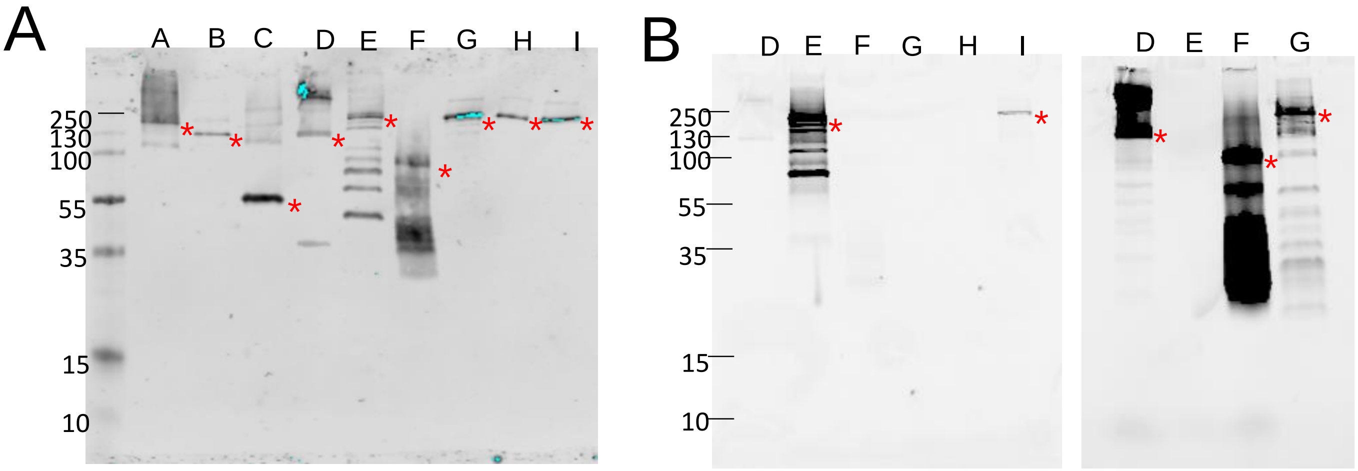


**Supplementary Figure S3.** Characterization of FP-fusions ##A-I by Western Blot and Fluorescent Gel Imaging.

Aliquots of lysates of insect cells expressing fusions ##A-I were separated by SDS PAGE; fluorescent gel imaging and Western blotting were performed using the same gel. #A-#I stays for the FP-fusion (Suppl. Dataset S1); bands corresponding to the position of full-length FP-fusions are designated with asterisk (*); molecular weight markers shown on the left. **A**: Western blot performed against 3C-cleavage site; **B:** fluorescent gel imaging acquired on Cy3 (left) and Cy5 (right) channels as described in Materials and Methods. Fusion A (fused FP – far-red mKate2) – fluorescence cannot be measured with available wave lengths.

Additional bands originate from incomplete translation and multimerization under non-reducing conditions. Intensive fluorescence and Western blot signals detected for fusion F below its full length sequence position indicated that full-length protein is only a minor fraction in the sample. Dendra-containing C-terminal fragments below 60kDa are x9 more abundant (quantified 160fmol) then its correctly expressed fusion (quantified 160 fmol and 19 fmol respectively). The case of fusion #F underlines the importance of gel separation to remove products of incomplete expression products before quantitative analysis of labelled proteins. In-solution analysis of fluorescence could bias the amount of fusion #F.

Fusion #A

```
   1 MGSAWSHPQF EKGGGSGGGS GGSAWSHPQF EKLEVLFQGP AAAMDSSSGS
  51 QGNGSFMDQN SLGILNMDNL KVFGDLSSFS KSGRRSGTGR DSPAVHERLR
 101 EERLRALETL EKPRPSRSSQ AKSTALQMRL STNISELVTD DTIDLNSSSS
 151 FALVTPATKG TNISFEPAEI TGRSTLCQAG KQRRREKPSL SVAEILKSSF
 201 VEKARLLERK LQDASQAEAS YHLSRGSSSS LSDSFNCSRS LPRTADMSEL
 251 SLNGGGGFSF GSASLAAEGQ DESFAPAELM QSKLVLGEIS WAQEFTAMPT
 301 ATLSTQALKK IAAPTSEIET SVSNSVLAGD PDFSLGNYFQ TRSENIWNIV
 351 SNSSPNRSRT NQPLLEPESN VTLDSVGEKT PQPDNKTYTK TDAITGSNLG
 401 RNLMRKMQQD RIETALKSRN GLAAKETKRP PSSSEILSLS AIDKALRDID
 451 LNSDTSTVEV VNHLWEHGRG NNYDDGENKE NQSSNSHAER TSCGSNKLTD
 501 TMSFTDSVLN STDFRHLQQS ISRKPLSPLA DHPQITISRA DTDPVETEAE
 551 ADIDEWPSTP VKEPSRRVRR TKISPRAASP SSSDGVRPLT CTEDENDEED
 601 EDKTPVNKKR VSIATMGLIP RKASSLSSTR LDGCDVAVAS STERDFGISP
 651 NLAKNLSPLS SPRSCLSSPL LDSTTSSDRR QLMSGAARKA NSSPAGSEAS
 701 STSGFTASGR RGLGTSSVAV PRSISRCSNS SEFSHRDGKL LPLKVTHTTL
 751 CWGSTKLRTD VRKSMQVKNT ADKRLVIRLG IQGPGFQLVG TDSSTITLQA
 801 MECRSVVINF CPTVCGAAIG ALSFYAPPGA HNSNQPGLEI PLYGYGGSAS
 851 ITINGLLKGP VGASFLTIGD VCELSAGPLS ANLKFHNKGP LTAFVGISVD
 901 STVLMKLRLS EAFEVRPSRL ILPPNSERCV QILFRPNRED LKNILKKTAQ
 951 VLTLANMRIF CGDEPNRQRM RSLVQRMSSR QREKLTSPML DSIWGEFPDE
1001 QPVRNIGQLN EPPEFILDLV TVVRIIDVAL TLNRDFDESS ESSLMFLPEA
1051 DETVLFRTIC VANSPTPTQS HLLEPVDELL EADTTTLTPV EQNWCVQPGL
1101 LEFGVGCSPL ATLSVTNSFC SRQLIEVTCN VPDLVRISPS ECYVSPDGGH
1151 AEIDVRLLRS PRQDLLEREP LIVVSMENER ISVPISFKFG APGSAGSAAG
1201 SGMVSELIKE NMHMKLYMEG TVNNHHFKCT SEGEGKPYEG TQTMRIKAVE
1251 GGPLPFAFDI LATSFMYGSK TFINHTQGIP DFFKQSFPEG FTWERVTTYE
1301 DGGVLTATQD TSLQDGCLIY NVKIRGVNFP SNGPVMQKKT LGWEASTETL
1351 YPADGGLEGR ADMALKLVGG GHLICNLKTT YRSKKPAKNL KMPGVYYVDR
1401 RLERIKEADK ETYVEQHEVA VARYCDLPSK LGHRLEVLFQ GPGSSHHHHH
1451 HSG
```

Fusion #B

```
   1 MGSSHHHHHH SSGRMKIEEG KLVIWINGDK GYNGLAEVGK KFEKDTGIKV
  51 TVEHPDKLEE KFPQVAATGD GPDIIFWAHD RFGGYAQSGL LAEITPDKAF
 101 QDKLYPFTWD AVRYNGKLIA YPIAVEALSL IYNKDLLPNP PKTWEEIPAL
 151 DKELKAKGKS ALMFNLQEPY FTWPLIAADG GYAFKYENGK YDIKDVGVDN
 201 AGAKAGLTFL VDLIKNKHMN ADTDYSIAEA AFNKGETAMT INGPWAWSNI
 251 DTSKVNYGVT VLPTFKGQPS KPFVGVLSAG INAASPNKEL AKEFLENYLL
 301 TDEGLEAVNK DKPLGAVALK SYEEELVKDP RIAATMENAQ KGEIMPNIPQ
 351 MSAFWYAVRT AVINAASGRQ TVDEALKDAQ TNSSSNNNNN NNNNNSSGRL
 401 EVLFQGPAAA MASNDYTQQA TQSYGAYPTQ PGQGYSQQSS QPYGQQSYSG
 451 YSQSTDTSGY GQSSYSSYGQ SQNTGYGTQS TPQGYGSTGG YGSSQSSQSS
 501 YGQQSSYPGY GQQPAPSSTS GSYGSSSQSS SYGQPQSGSY SQQPSYGGQQ
 551 QSYGQQQSYN PPQGYGQQNQ YNSSSGGGGG GGGGGNYGQD QSSMSSGGGS
 601 GGGYGNQDQS GGGGSGGYGQ QDRGGRGRGG SGGGGGGGGG GYNRSSGGYE
 651 PRGCGGGRGG RGGMGGSDRG GFNKFGGPRD QGSRHDSEQD NSDNNTIFVQ
 701 GLGENVTIES VADYFKQIGI IKTNKKTGQP MINLYTDRET GKLKGEATVS
 751 FDDPPSAKAA IDWFDGKEFS GNPIKVSFAT RRADFNRGGG NGRGGRGRGG
 801 PMGRGGYGGG GSGGGGRGGF PSGGGGGGGQ QRAGDWKCPN PTCENMNFSW
 851 RNECNQCKAP KPDGPGGGPG GSHMGGNYGD DRRGGRGGYD RGGYRGRGGD
 901 RGGFRGGRGG GDRGGFGPGK MDSRGEHRQD RRERPYGAPG SSSGRENLYF
 951 QGMDKDCEMK RTTLDSPLGK LELSGCEQGL HRIIFLGKGT SAADAVEVPA
1001 PAAVLGGPEP LMQATAWLNA YFHQPEAIEE FPVPALHHPV FQQESFTRQV
1051 LWKLLKVVKF GEVISYSHLA ALAGNPAATA AVKTALSGNP VPILIPCHRV
1101 VQGDLDVGGY EGGLAVKEWL LAHEGHRLGK PGLG
```

Fusion #C

```
   1 MGSSHHHHHH SSGRLEVLFQ GPMAEIGTGF PFDPHYVEVL GERMHYVDVG
  51 PRDGTPVLFL HGNPTSSYVW RNIIPHVAPT HRCIAPDLIG MGKSDKPDLG
 101 YFFDDHVRFM DAFIEALGLE EVVLVIHDWG SALGFHWAKR NPERVKGIAF
 151 MEFIRPIPTW DEWPEFARET FQAFRTTDVG RKLIIDQNVF IEGTLPMGVV
 201 RPLTEVEMDH YREPFLNPVD REPLWRFPNE LPIAGEPANI VALVEEYMDW
 251 LHQSPVPKLL FWGTPGVLIP PAEAARLAKS LPNCKAVDIG PGLNLLQEDN
 301 PDLIGSEIAR WLSTLEISGG SAGSAAGSGA AAMASLGLQL VGYILGLLGL
 351 LGTLVAMLLP SWKTSSYVGA SIVTAVGFSK GLWMECATHS TGITQCDIYS
 401 TLLGLPADIQ AAQAMMVTSS AISSLACIIS VVGMRCTVFC QESRAKDRVA
 451 VAGGVFFILG GLLGFIPVAW NLHGILRDFY SPLVPDSMKF EIGEALYLGI
 501 ISSLFSLIAG IILCFSCSSQ RNRSNYYDAY QAQPLATRSS PRPGQPPKVK
 551 SEFNSYSLTG YV
```

Fusion #D

```
   1 MAAAMADLAE CNIKVMCRFR PLNESEVNRG DKYIAKFQGE DTVVIASKPY
  51 AFDRVFQSST SQEQVYNDCA KKIVKDVLEG YNGTIFAYGQ TSSGKTHTME
 101 GKLHDPEGMG IIPRIVQDIF NYIYSMDENL EFHIKVSYFE IYLDKIRDLL
 151 DVSKTNLSVH EDKNRVPYVK GCTERFVCSP DEVMDTIDEG KSNRHVAVTN
 201 MNEHSSRSHS IFLINVKQEN TQTEQKLSGK LYLVDLAGSE KVSKTGAEGA
 251 VLDEAKNINK SLSALGNVIS ALAEGSTYVP YRDSKMTRIL QDSLGGNCRT
 301 TIVICCSPSS YNESETKSTL LFGQRAKTIK NTVCVNVELT AEQWKKKYEK
 351 EKEKNKILRN TIQWLENELN RWRNGETVPI DEQFDKEKAN LEAFTVDKDI
 401 TLTNDKPATA IGVIGNFTDA ERRKCEEEIA KLYKQLDDKD EEINQQSQLV
 451 EKLKTQMLDQ EELLASTRRD QDNMQAELNR LQAENDASKE EVKEVLQALE
 501 ELAVNYDQKS QEVEDKTKEY ELLSDELNQK SATLASIDAE LQKLKEMTNH
 551 QKKRAAEMMA SLLKDLAEIG IAVGNNDVKQ PEGTGMIDEE FTVARLYISK
 601 MKSEVKTMVK RCKQLESTQT ESNKKMEENE KELAACQLRI SQHEAKIKSL
 651 TEYLQNVEQK KRQLEESVDA LSEELVQLRA QEKVHEMEKE HLNKVQTANE
 701 VKQAVEQQIQ SHRETHQKQI SSLRDEVEAK AKLITDLQDQ NQKMMLEQER
 751 LRVEHEKLKA TDQEKSRKLH ELTVMQDRRE QARQDLKGLE ETVAKELQTL
 801 HNLRKLFVQD LATRVKKSAE IDSDDTGGSA AQKQKISFLE NNLEQLTKVH
 851 KQLVRDNADL RCELPKLEKR LRATAERVKA LESALKEAKE NASRDRKRYQ
 901 QEVDRIKEAG APGSAGSAAG SGMVSKGEED NMASLPATHE LHIFGSINGV
 951 DFDMVGQGTG NPNDGYEELN LKSTKGDLQF SPWILVPHIG YGFHQYLPYP
1001 DGMSPFQAAM VDGSGYQVHR TMQFEDGASL TVNYRYTYEG SHIKGEAQVK
1051 GTGFPADGPV MTNSLTAADW CRSKKTYPND KTIISTFKWS YTTGNGKRYR
1101 STARTTYTFA KPMAANYLKN QPMYVFRKTE LKHSKTELNF KEWQKAFTDV
1151 MGMDELYKLE VLFQGPGSSH HHHHSG
```

Fusion #E

```
   1 MAAAMASVKV AVRVRPMNRR EKDLEAKFII QMEKSKTTIT NLKIPEGGTG
  51 DSGRERTKTF TYDFSFYSAD TKSPDYVSQE MVFKTLGTDV VKSAFEGYNA
 101 CVFAYGQTGS GKSYTMMGNS GDSGLIPRIC EGLFSRINET TRWDEASFRT
 151 EVSYLEIYNE RVRDLLRRKS SKTFNLRVRE HPKEGPYVED LSKHLVQNYG
 201 DVEELMDAGN INRTTAATGM NDVSSRSHAI FTIKFTQAKF DSEMPCETVS
 251 KIHLVDLAGS ERADATGATG VRLKEGGNIN KSLVTLGNVI SALADLSQDA
 301 ANTLAKKKQV FVPYRDSVLT WLLKDSLGGN SKTIMIATIS PADVNYGETL
 351 STLRYANRAK NIINKPTINE DANVKLIREL RAEIARLKTL LAQGNQIALL
 401 DSPTALSMEE KLQQNEARVQ ELTKEWTNKW NETQNILKEQ TLALRKEGIG
 451 VVLDSELPHL IGIDDDLLST GIILYHLKEG QTYVGRDDAS TEQDIVLHGL
 501 DLESEHCIFE NIGGTVTLIP LSGSQCSVNG VQIVEATHLN QGAVILLGRT
 551 NMFRFNHPKE AAKLREKRKS GLLSSFSLSM TDLSKSRENL SAVMLYNPGL
 601 EFERQQREEL EKLESKRKLI EEMEEKQKSD KAELERMQQE VETQRKETEI
 651 VQLQIRKQEE SLKRRSFHIE NKLKDLLAEK EKFEEERLRE QQEIELQKKR
 701 QEEETFLRVQ EELQRLKELN NNEKAEKFQI FQELDQLQKE KDEQYAKLEL
 751 EKKRLEEQEK EQVMLVAHLE EQLREKQEMI QLLRRGEVQW VEEEKRDLEG
 801 IRESLLRVKE ARAGGDEDGE ELEKAQLRFF EFKRRQLVKL VNLEKDLVQQ
 851 KDILKKEVQE EQEILECLKC EHDKESRLLE KHDESVTDVT EVPQDFEKIK
 901 PVEYRLQYKE RQLQYLLQNH LPTLLEEKQR AFEILDRGPL SLDNTLYQVE
 951 KEMEEKEEQL AQYQANANQL QKLQATFEFT ANIARQEEKV RKKEKEILES
1001 REKQQREALE RALARLERRH SALQRHSTLG TEIEEQRQKL ASLNSGSREQ
1051 SGLQASLEAE QEALEKDQER LEYEIQQLKQ KIYEVDGVQK DHHGTLEGKV
1101 ASSSLPVSAE KSHLVPLMDA RINAYIEEEV QRRLQDLHRV ISEGCSTSAD
1151 TMKDNEKLHN GTIQRKLKYE RMVSRSLGAN PDDLKDPIKI SIPRYVLCGQ
1201 GKDAHFEFEV KITVLDETWT VFRRYSRFRE MHKTLKLKYA ELAALEFPPK
1251 KLFGNKDERV IAERRSHLEK YLRDFFSVML QSATSPLHIN KVGLTLSKHT
1301 ICEFSPFFKK GVFDYSSHGT GGAPGSAGSA AGSGMVSKGE AVIKEFMRFK
1351 VHMEGSMNGH EFEIEGEGEG RPYEGTQTAK LKVTKGGPLP FSWDILSPQF
1401 MYGSRAFIKH PADIPDYYKQ SFPEGFKWER VMNFEDGGAV TVTQDTSLED
1451 GTLIYKVKLR GTNFPPDGPV MQKKTMGWEA STERLYPEDG VLKGDIKMAL
1501 RLKDGGRYLA DFKTTYKAKK PVQMPGAYNV DRKLDITSHN EDYTVVEQYE
1551 RSEGRHSTGG MDELYKNAPL EVLFQGPGSS MKIEEGKLVI WINGDKGYNG
1601 LAEVGKKFEK DTGIKVTVEH PDKLEEKFPQ VAATGDGPDI IFWAHDRFGG
1651 YAQSGLLAEI TPDKAFQDKL YPFTWDAVRY NGKLIAYPIA VEALSLIYNK
1701 DLLPNPPKTW EEIPALDKEL KAKGKSALMF NLQEPYFTWP LIAADGGYAF
1751 KYENGKYDIK DVGVDNAGAK AGLTFLVDLI KNKHMNADTD YSIAEAAFNK
1801 GETAMTINGP WAWSNIDTSK VNYGVTVLPT FKGQPSKPFV GVLSAGINAA
1851 SPNKELAKEF LENYLLTDEG LEAVNKDKPL GAVALKSYEE ELVKDPRIAA
1901 TMENAQKGEI MPNIPQMSAF WYAVRTAVIN AASGRQTVDE ALKDAQTNSS
1951 SNNNNNNNNN NSG
```

Fusion #F

```
   1 MAAAAEPRQE FEVMEDHAGT YGLGDRKDQG GYTMHQDQEG DTDAGLKESP
  51 LQTPTEDGSE EPGSETSDAK STPTAEDVTA PLVDEGAPGK QAAAQPHTEI
 101 PEGTTAEEAG IGDTPSLEDE AAGHVTQARM VSKSKDGTGS DDKKAKGADG
 151 KTKIATPRGA APPGQKGQAN ATRIPAKTPP APKTPPSSGE PPKSGDRSGY
 201 SSPGSPGTPG SRSRTPSLPT PPTREPKKVA VVRTPPKSPS SAKSRLQTAP
 251 VPMPDLKNVK SKIGSTENLK HQPGGGKVQI INKKLDLSNV QSKCGSKDNI
 301 KHVPGGGSVQ IVYKPVDLSK VTSKCGSLGN IHHKPGGGQV EVKSEKLDFK
 351 DRVQSKIGSL DNITHVPGGG NKKIETHKLT FRENAKAKTD HGAEIVYKSP
 401 VVSGDTSPRH LSNVSSTGSI DMVDSPQLAT LADEVSASLA KQGLGAPGSA
 451 GSAAGSGMNT PGINLIKEDM RVKVHMEGNV NGHAFVIEGE GKGKPYEGTQ
 501 TANLTVKEGA PLPFSYDILT TAVHYGNRVF TKYPEDIPDY FKQSFPEGYS
 551 WERTMTFEDK GICTIRSDIS LEGDCFFQNV RFKGTNFPPN GPVMQKKTLK
 601 WEPSTEKLHV RDGLLVGNIN MALLLEGGGH YLCDFKTTYK AKKVVQLPDA
 651 HFVDHRIEIL GNDSDYNKVK LYEHAVARYS PLPSQVWLEV LFQGPGSSHH
 701 HHHSG
```

Fusion #G

```
   1 MGHHHHHHSS GRMKIEEGKL VIWINGDKGY NGLAEVGKKF EKDTGIKVTV
  51 EHPDKLEEKF PQVAATGDGP DIIFWAHDRF GGYAQSGLLA EITPDKAFQD
 101 KLYPFTWDAV RYNGKLIAYP IAVEALSLIY NKDLLPNPPK TWEEIPALDK
 151 ELKAKGKSAL MFNLQEPYFT WPLIAADGGY AFKYENGKYD IKDVGVDNAG
 201 AKAGLTFLVD LIKNKHMNAD TDYSIAEAAF NKGETAMTIN GPWAWSNIDT
 251 SKVNYGVTVL PTFKGQPSKP FVGVLSAGIN AASPNKELAK EFLENYLLTD
 301 EGLEAVNKDK PLGAVALKSY EEELVKDPRI AATMENAQKG EIMPNIPQMS
 351 AFWYAVRTAV INAASGRQTV DEALKDAQTN SSSNNNNNNN NNNSSGRLEV
 401 LFQGPAAAMP VRGDRGFPPR RELSGWLRAP GMEELIWEQY TVTLQKDSKR
 451 GFGIAVSGGR DNPHFENGET SIVISDVLPG GPADGLLQEN DRVVMVNGTP
 501 MEDVLHSFAV QQLRKSGKVA AIVVKRPRKV QVAALQASPP LDQDDRAFEV
 551 MDEFDGRSFR SGYSERSRLN SHGGRSRSWE DSPERGRPHE RARSRERDLS
 601 RDRSRGRSLE RGLDQDHART RDRSRGRSLE RGLDHDFGPS RDRDRDRSRG
 651 RSIDQDYERA YHRAYDPDYE RAYSPEYRRG ARHDARSRGP RSRSREHPHS
 701 RSPSPEPRGR PGPIGVLLMK SRANEEYGLR LGSQIFVKEM TRTGLATKDG
 751 NLHEGDIILK INGTVTENMS LTDARKLIEK SRGKLQLVVL RDSQQTLINI
 801 PSLNDSDSEI EDISEIESNR SFSPEERRHQ YSDYDYHSSS EKLKERPSSR
 851 EDTPSRLSRM GATPTPFKST GDIAGTVVPE TNKEPRYQED PPAPQPKAAP
 901 RTFLRPSPED EAIYGPNTKM VRFKKGDSVG LRLAGGNDVG IFVAGIQEGT
 951 SAEQEGLQEG DQILKVNTQD FRGLVREDAV LYLLEIPKGE MVTILAQSRA
1001 DVYRDILACG RGDSFFIRSH FECEKETPQS LAFTRGEVFR VVDTLYDGKL
1051 GNWLAVRIGN ELEKGLIPNK SRAEQMASVQ NAQRDNAGDR ADFWRMRGQR
1101 SGVKKNLRKS REDLTAVVSV STKFPAYERV LLREAGFKRP VVLFGPIADI
1151 AMEKLANELP DWFQTAKTEP KDAGSEKSTG VVRLNTVRQI IEQDKHALLD
1201 VTPKAVDLLN YTQWFPIVIF FNPDSRQGVK TMRQRLNPTS NKSSRKLFDQ
1251 ANKLKKTCAH LFTATINLNS ANDSWFGSLK DTIQHQQGEA VWVSEGKMEG
1301 MDDDPEDRMS YLTAMGADYL SCDSRLISDF EDTDGEGGAY TDNELDEPAE
1351 EPLVSSITRS SEPVQHEESI RKPSPEPRAQ MRRAASSDQL RDNSPPPAFK
1401 PEPPKAKTQN KEESYDFSKS YEYKSNPSAV AGNETPGAST KGYPPPVAAK
1451 PTFGRSILKP STPIPPQEGE EVGESSEEQD NAPKSVLGKV KIFEKMDHKA
1501 RLQRMQELQE AQNARIEIAQ KHPDIYAVPI KTHKPDPGTP QHTSSRPPEP
1551 QKAPSRPYQD TRGSYGSDAE EEEYRQQLSE HSKRGYYGQS ARYRDTELGA
1601 PGSSSGRENL YFQGMVSKGE ELFTGVVPIL VELDGDVNGH KFSVSGEGEG
1651 DATYGKLTLK FICTTGKLPV PWPTLVTTLT YGVQCFSRYP DHMKQHDFFK
1701 SAMPEGYVQE RTIFFKDDGN YKTRAEVKFE GDTLVNRIEL KGIDFKEDGN
1751 ILGHKLEYNY NSHNVYIMAD KQKNGIKVNF KIRHNIEDGS VQLADHYQQN
1801 TPIGDGPVLL PDNHYLSTQS KLSKDPNEKR DHMVLLEFVT AAGITLGMDE
1851 LYK
```

Fusion #H

```
   1 MGSSHHHHHH SSGRMKIEEG KLVIWINGDK GYNGLAEVGK KFEKDTGIKV
  51 TVEHPDKLEE KFPQVAATGD GPDIIFWAHD RFGGYAQSGL LAEITPDKAF
 101 QDKLYPFTWD AVRYNGKLIA YPIAVEALSL IYNKDLLPNP PKTWEEIPAL
 151 DKELKAKGKS ALMFNLQEPY FTWPLIAADG GYAFKYENGK YDIKDVGVDN
 201 AGAKAGLTFL VDLIKNKHMN ADTDYSIAEA AFNKGETAMT INGPWAWSNI
 251 DTSKVNYGVT VLPTFKGQPS KPFVGVLSAG INAASPNKEL AKEFLENYLL
 301 TDEGLEAVNK DKPLGAVALK SYEEELVKDP RIAATMENAQ KGEIMPNIPQ
 351 MSAFWYAVRT AVINAASGRQ TVDEALKDAQ TNSSSNNNNN NNNNNSSGRL
 401 EVLFQGPMDK DCEMKRTTLD SPLGKLELSG CEQGLHRIIF LGKGTSAADA
 451 VEVPAPAAVL GGPEPLMQAT AWLNAYFHQP EAIEEFPVPA LHHPVFQQES
 501 FTRQVLWKLL KVVKFGEVIS YSHLAALAGN PAATAAVKTA LSGNPVPILI
 551 PCHRVVQGDL DVGGYEGGLA VKEWLLAHEG HRLGKPGLGG SSSGRENLYF
 601 QGAAAMPVRG DRGFPPRREL SGWLRAPGME ELIWEQYTVT LQKDSKRGFG
 651 IAVSGGRDNP HFENGETSIV ISDVLPGGPA DGLLQENDRV VMVNGTPMED
 701 VLHSFAVQQL RKSGKVAAIV VKRPRKVQVA ALQASPPLDQ DDRAFEVMDE
 751 FDGRSFRSGY SERSRLNSHG GRSRSWEDSP ERGRPHERAR SRERDLSRDR
 801 SRGRSLERGL DQDHARTRDR SRGRSLERGL DHDFGPSRDR DRDRSRGRSI
 851 DQDYERAYHR AYDPDYERAY SPEYRRGARH DARSRGPRSR SREHPHSRSP
 901 SPEPRGRPGP IGVLLMKSRA NEEYGLRLGS QIFVKEMTRT GLATKDGNLH
 951 EGDIILKING TVTENMSLTD ARKLIEKSRG KLQLVVLRDS QQTLINIPSL
1001 NDSDSEIEDI SEIESNRSFS PEERRHQYSD YDYHSSSEKL KERPSSREDT
1051 PSRLSRMGAT PTPFKSTGDI AGTVVPETNK EPRYQEDPPA PQPKAAPRTF
1101 LRPSPEDEAI YGPNTKMVRF KKGDSVGLRL AGGNDVGIFV AGIQEGTSAE
1151 QEGLQEGDQI LKVNTQDFRG LVREDAVLYL LEIPKGEMVT ILAQSRADVY
1201 RDILACGRGD SFFIRSHFEC EKETPQSLAF TRGEVFRVVD TLYDGKLGNW
1251 LAVRIGNELE KGLIPNKSRA EQMASVQNAQ RDNAGDRADF WRMRGQRSGV
1301 KKNLRKSRED LTAVVSVSTK FPAYERVLLR EAGFKRPVVL FGPIADIAME
1351 KLANELPDWF QTAKTEPKDA GSEKSTGVVR LNTVRQIIEQ DKHALLDVTP
1401 KAVDLLNYTQ WFPIVIFFNP DSRQGVKTMR QRLNPTSNKS SRKLFDQANK
1451 LKKTCAHLFT ATINLNSAND SWFGSLKDTI QHQQGEAVWV SEGKMEGMDD
1501 DPEDRMSYLT AMGADYLSCD SRLISDFEDT DGEGGAYTDN ELDEPAEEPL
1551 VSSITRSSEP VQHEESIRKP SPEPRAQMRR AASSDQLRDN SPPPAFKPEP
1601 PKAKTQNKEE SYDFSKSYEY KSNPSAVAGN ETPGASTKGY PPPVAAKPTF
1651 GRSILKPSTP IPPQEGEEVG ESSEEQDNAP KSVLGKVKIF EKMDHKARLQ
1701 RMQELQEAQN ARIEIAQKHP DIYAVPIKTH KPDPGTPQHT SSRPPEPQKA
1751 PSRPYQDTRG SYGSDAEEEE YRQQLSEHSK RGYYGQSARY RDTELGAP
```

Fusion #I

```
   1 MGSSHHHHHH SSGRMKIEEG KLVIWINGDK GYNGLAEVGK KFEKDTGIKV
  51 TVEHPDKLEE KFPQVAATGD GPDIIFWAHD RFGGYAQSGL LAEITPDKAF
 101 QDKLYPFTWD AVRYNGKLIA YPIAVEALSL IYNKDLLPNP PKTWEEIPAL
 151 DKELKAKGKS ALMFNLQEPY FTWPLIAADG GYAFKYENGK YDIKDVGVDN
 201 AGAKAGLTFL VDLIKNKHMN ADTDYSIAEA AFNKGETAMT INGPWAWSNI
 251 DTSKVNYGVT VLPTFKGQPS KPFVGVLSAG INAASPNKEL AKEFLENYLL
 301 TDEGLEAVNK DKPLGAVALK SYEEELVKDP RIAATMENAQ KGEIMPNIPQ
 351 MSAFWYAVRT AVINAASGRQ TVDEALKDAQ TNSSSNNNNN NNNNNSSGRL
 401 EVLFQGPMVS KGEEDNMAII KEFMRFKVHM EGSVNGHEFE IEGEGEGRPY
 451 EGTQTAKLKV TKGGPLPFAW DILSPQFMYG SKAYVKHPAD IPDYLKLSFP
 501 EGFKWERVMN FEDGGVVTVT QDSSLQDGEF IYKVKLRGTN FPSDGPVMQK
 551 KTMGWEASSE RMYPEDGALK GEIKQRLKLK DGGHYDAEVK TTYKAKKPVQ
 601 LPGAYNVNIK LDITSHNEDY TIVEQYERAE GRHSTGGMDE LYKGSAGSAA
 651 GSGAAAMPVR GDRGFPPRRE LSGWLRAPGM EELIWEQYTV TLQKDSKRGF
 701 GIAVSGGRDN PHFENGETSI VISDVLPGGP ADGLLQENDR VVMVNGTPME
 751 DVLHSFAVQQ LRKSGKVAAI VVKRPRKVQV AALQASPPLD QDDRAFEVMD
 801 EFDGRSFRSG YSERSRLNSH GGRSRSWEDS PERGRPHERA RSRERDLSRD
 851 RSRGRSLERG LDQDHARTRD RSRGRSLERG LDHDFGPSRD RDRDRSRGRS
 901 IDQDYERAYH RAYDPDYERA YSPEYRRGAR HDARSRGPRS RSREHPHSRS
 951 PSPEPRGRPG PIGVLLMKSR ANEEYGLRLG SQIFVKEMTR TGLATKDGNL
1001 HEGDIILKIN GTVTENMSLT DARKLIEKSR GKLQLVVLRD SQQTLINIPS
1051 LNDSDSEIED ISEIESNRSF SPEERRHQYS DYDYHSSSEK LKERPSSRED
1101 TPSRLSRMGA TPTPFKSTGD IAGTVVPETN KEPRYQEDPP APQPKAAPRT
1151 FLRPSPEDEA IYGPNTKMVR FKKGDSVGLR LAGGNDVGIF VAGIQEGTSA
1201 EQEGLQEGDQ ILKVNTQDFR GLVREDAVLY LLEIPKGEMV TILAQSRADV
1251 YRDILACGRG DSFFIRSHFE CEKETPQSLA FTRGEVFRVV DTLYDGKLGN
1301 WLAVRIGNEL EKGLIPNKSR AEQMASVQNA QRDNAGDRAD FWRMRGQRSG
1351 VKKNLRKSRE DLTAVVSVST KFPAYERVLL REAGFKRPVV LFGPIADIAM
1401 EKLANELPDW FQTAKTEPKD AGSEKSTGVV RLNTVRQIIE QDKHALLDVT
1451 PKAVDLLNYT QWFPIVIFFN PDSRQGVKTM RQRLNPTSNK SSRKLFDQAN
1501 KLKKTCAHLF TATINLNSAN DSWFGSLKDT IQHQQGEAVW VSEGKMEGMD
1551 DDPEDRMSYL TAMGADYLSC DSRLISDFED TDGEGGAYTD NELDEPAEEP
1601 LVSSITRSSE PVQHEESIRK PSPEPRAQMR RAASSDQLRD NSPPPAFKPE
1651 PPKAKTQNKE ESYDFSKSYE YKSNPSAVAG NETPGASTKG YPPPVAAKPT
1701 FGRSILKPST PIPPQEGEEV GESSEEQDNA PKSVLGKVKI FEKMDHKARL
1751 QRMQELQEAQ NARIEIAQKH PDIYAVPIKT HKPDPGTPQH TSSRPPEPQK
1801 APSRPYQDTR GSYGSDAEEE EYRQQLSEHS KRGYYGQSAR YRDTELGAP
```

**Supplementary Figure S4**. Peptide mapping of FP-fusions by mass spectrometry. Peptides detected by ms-proteomics analysis on the tryptic digests of each FP-fusion are shown in red.

.

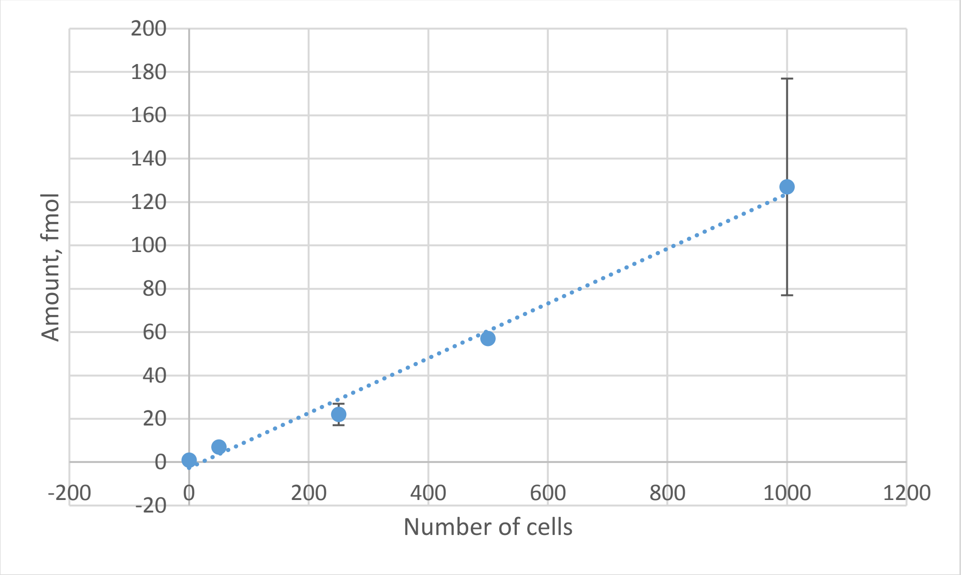


**Supplementary Figure S5.** Absolute quantification of EGFP in stably transfected HeLa cells with qFP-8 standard. Cell lysate of HeLa stably expressing mEGFP (Suppl. Table S4, #2) was separated by SDS PAGE, FP band digested with trypsin and equivalent of 1000, 500, 250 and 50 cells analyzed by mass spectrometry using qFP-8 peptide proxies.

A

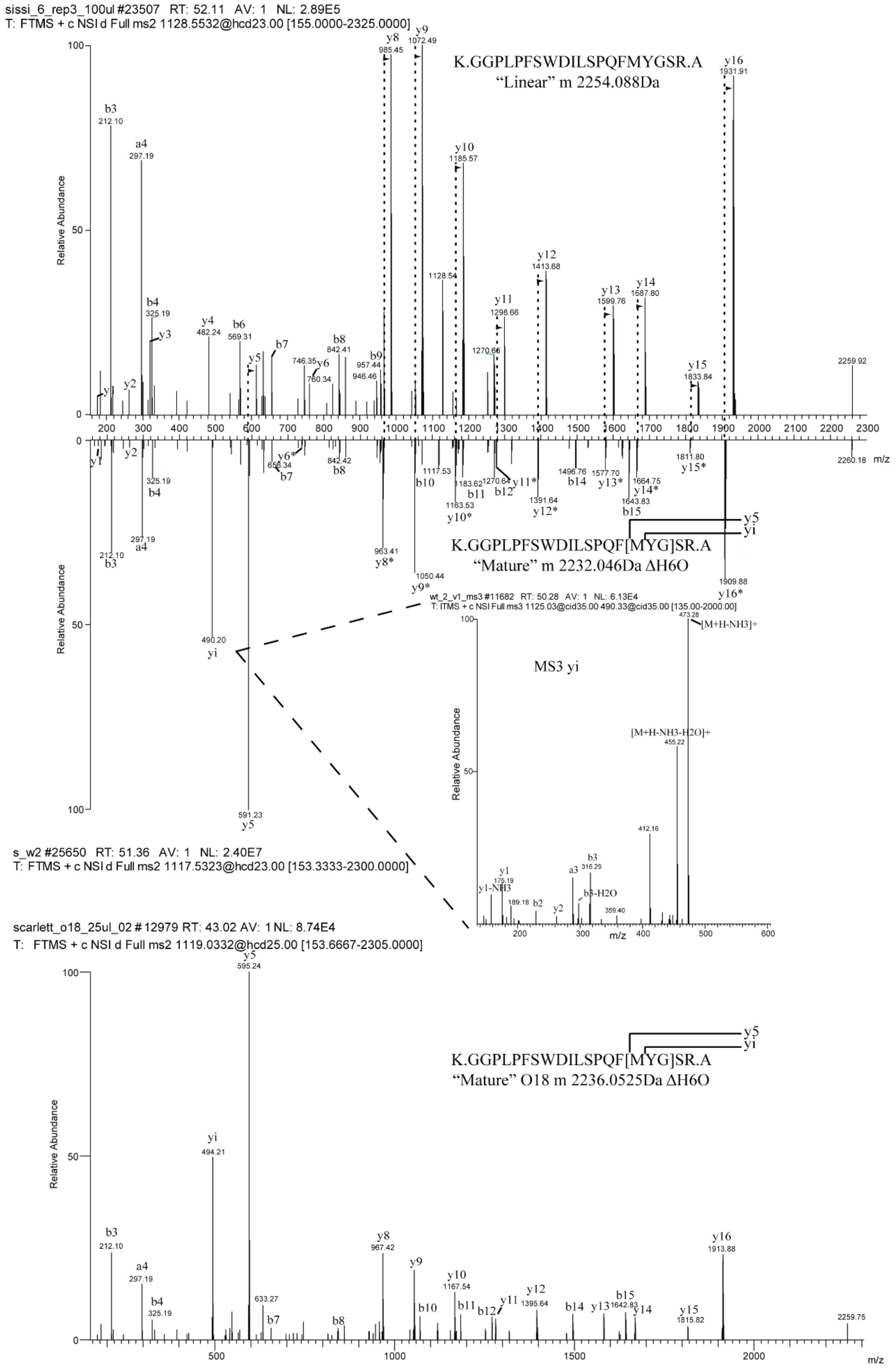
sissi_6_rep3_100ul #23507 RT: 52.11 AV: 1 NL: 2.89E5
T: FTMS + c NSI d Full ms2 1128.5532@hcd23.00 [155.0000-2325.0000]
K.GGPLPFSWDILSPQFMYGSR.A
"Linear" m 2254.088Da
Relative Abundance
m/z
K.GGPLPFSWDILSPQF[MYG]SR.A
"Mature" m 2232.046Da ΔH6O
wt_2_v1_ms3 #11682 RT: 50.28 AV: 1 NL: 6.13E4
T: ITMS + c NSI Full ms3 1125.03@cid35.00 490.33@cid35.00 [135.00-2000.00]
MS3 yi
[M+H-NH3]+
[M+H-NH3-H2O]+
s_w2 #25650 RT: 51.36 AV: 1 NL: 2.40E7
T: FTMS + c NSI d Full ms2 1117.5323@hcd23.00 [153.3333-2300.0000]
scarlett_o18_25ul_02 # 12979 RT: 43.02 AV: 1 NL: 8.74E4
T: FTMS + c NSI d Full ms2 1119.0332@hcd25.00 [153.6667-2305.0000]
K.GGPLPFSWDILSPQF[MYG]SR.A
"Mature" O18 m 2236.0525Da ΔH6O

B

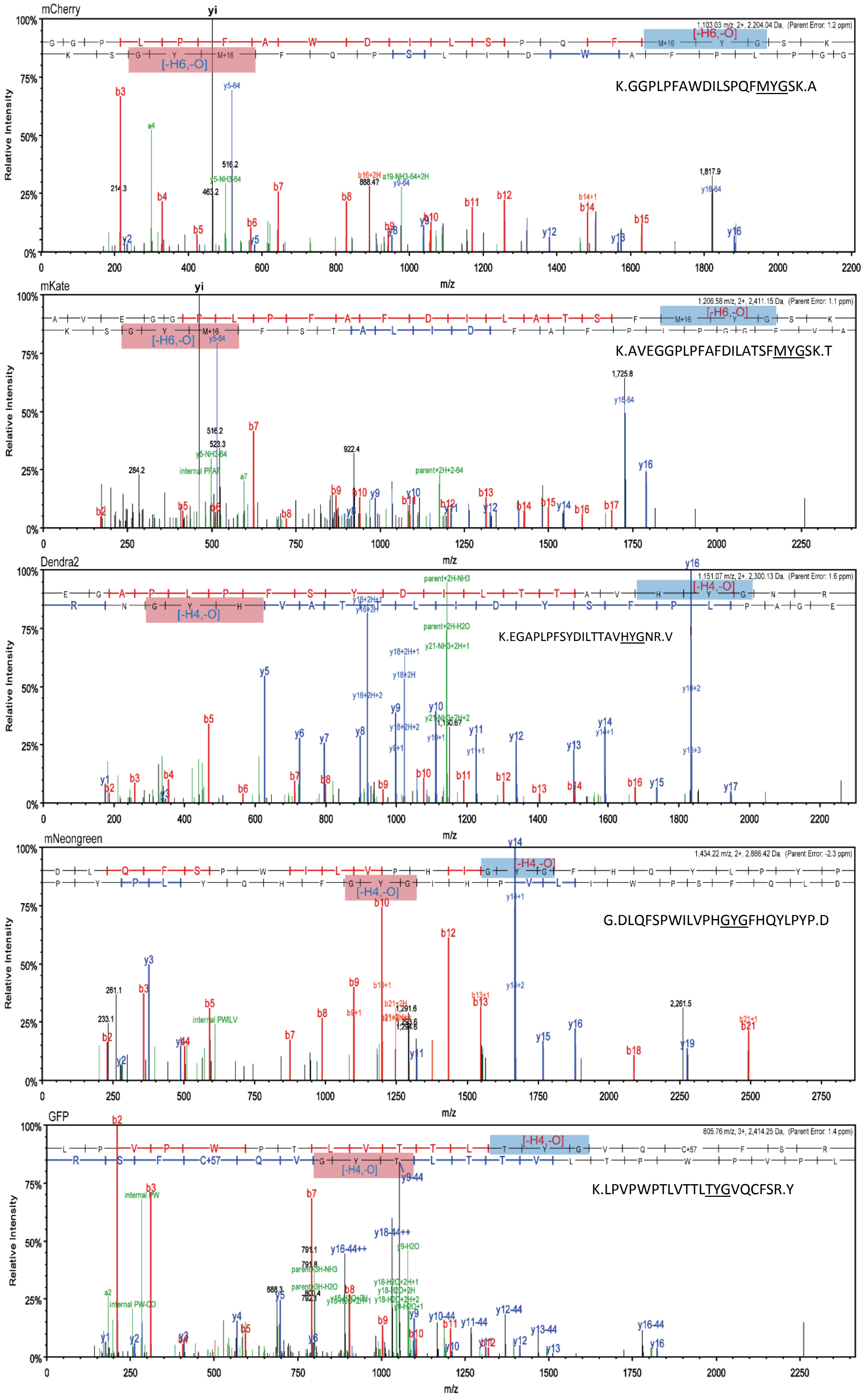

mCherry
K.GGPLPFAWDILSPQFMYGSK.A
[-H6,-O]
mKate
K.AVEGGPLPFAFDILATSFMYGSK.T
[-H6,-O]
Dendra2
K.EGAPLPFSYDILTTAVHYGNR.V
[-H4,-O]
mNeongreen
G.DLQFSPWILVPHGYGFHQYLPYP.D
[-H4,-O]
GFP
K.LPVPWPTLVTTLTYGVQCFSR.Y
[-H4,-O]
Relative Intensity
m/z

**Supplementary Figure S6.** Fragmentation spectra of chromophore-containing peptides from red- and green-type FPs

A. Fragmentation spectra of mScarlet-I peptide with linear and mature (cyclic) chromophore tripeptide …MYG… Abundant fragment $y_i$ is formed by cleavage of internal chemical bond within the chromophore cycle. The identity of yi was confirmed by MS3 fragmentation (insert) and meatabolic labeling of C-terminal arginine residue with $H_2^{18}O$ (lower panel, all y-ions show characteristic 4Da mass shift). Note that, upon digestion with trypsin in $H_2^{18}O$ water arginine residues exchange both oxygene atoms within their C-terminal carboxyl group.
B. Fragmentation spectra of peptides comprising mature (cyclic) chromophore detected in red-type (mKate2, mCherry) and green-type FPs (Dendra2, mNeonGreen, EGFP). Signature of two abundant fragments – yi (resulted from cleavage of internal bound withing cyclic chromophore) and y5 - is characteristic for red type of mature chromophore. GFP peptide with cyclic chromophore …TYG… shows characteristic series of chromophore-containing y-ions with the $\Delta m$=44.026 Da. Chromophore sequence is highlighted in blue and red for y- and b-ions, respectively.

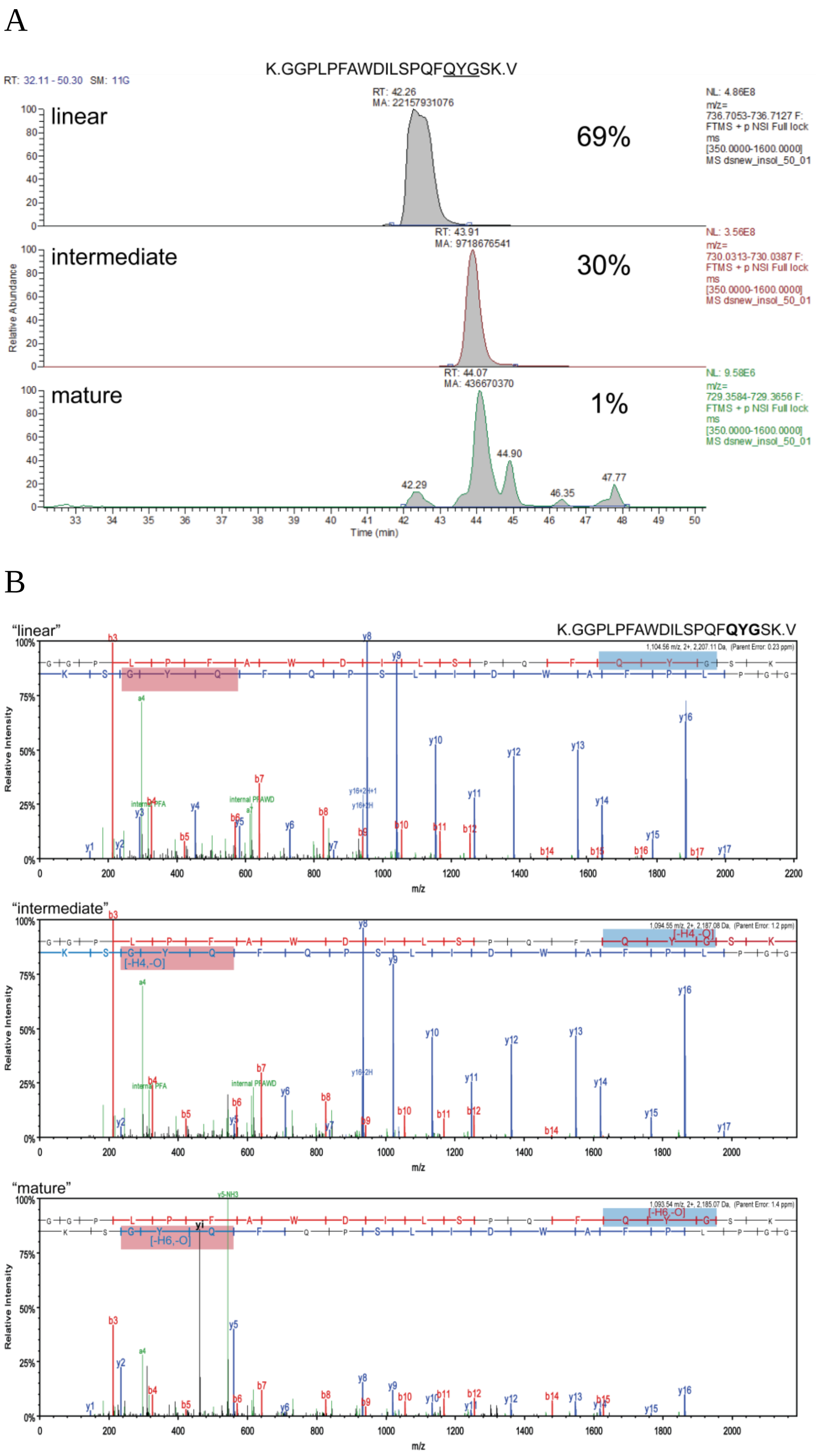


**Supplementary Figure S7.** Chromophore-containing peptides detected in red-type FP dsRed-express by mass spectrometry. Extracted Ion Chromatograms (XICs) (**Panel A**) and fragmentation spectra (**Panel B**) of dsRed-express peptide GGPLPFAWDILSPQF<u>QYG</u>SK comprising linear, intermediate and mature cyclic forms of …QYG… chromophore tripeptide.

A

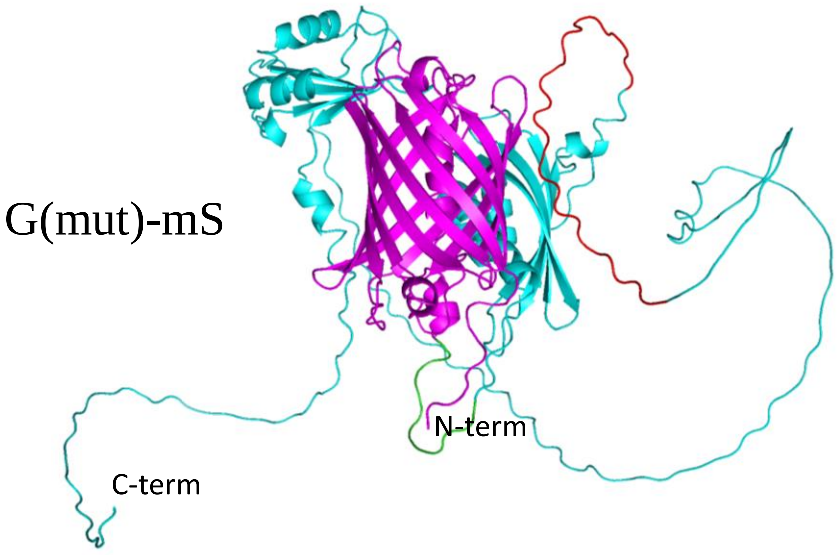


B

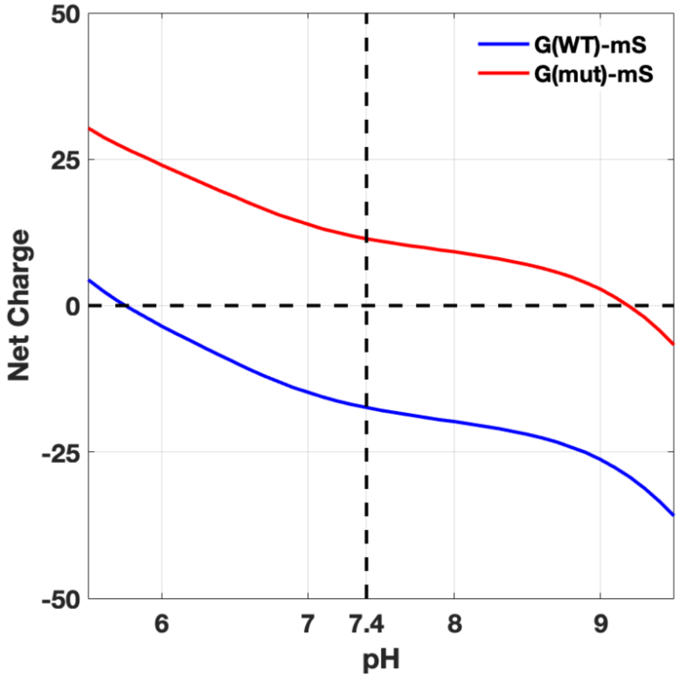


**Supplementary Figure S8**. Removal of glutamic acid residues from intrinsically disordered region changed formal net charge of the mScarlet-tagged protein G3BP1. **A**: 3D structure of the G(mut)-mS where glutamic acid residues are removed from the intrinsically disordered region (IDR). G3BP1 is shown in magenta, with the IDR (red) after removal of glutamic acid residues and short spacer sequence (green); fused red FP mScarlet-I is shown in pink. **B**: Net formal charge to pH curve for G(WT)_mS and G(mut)-mS.

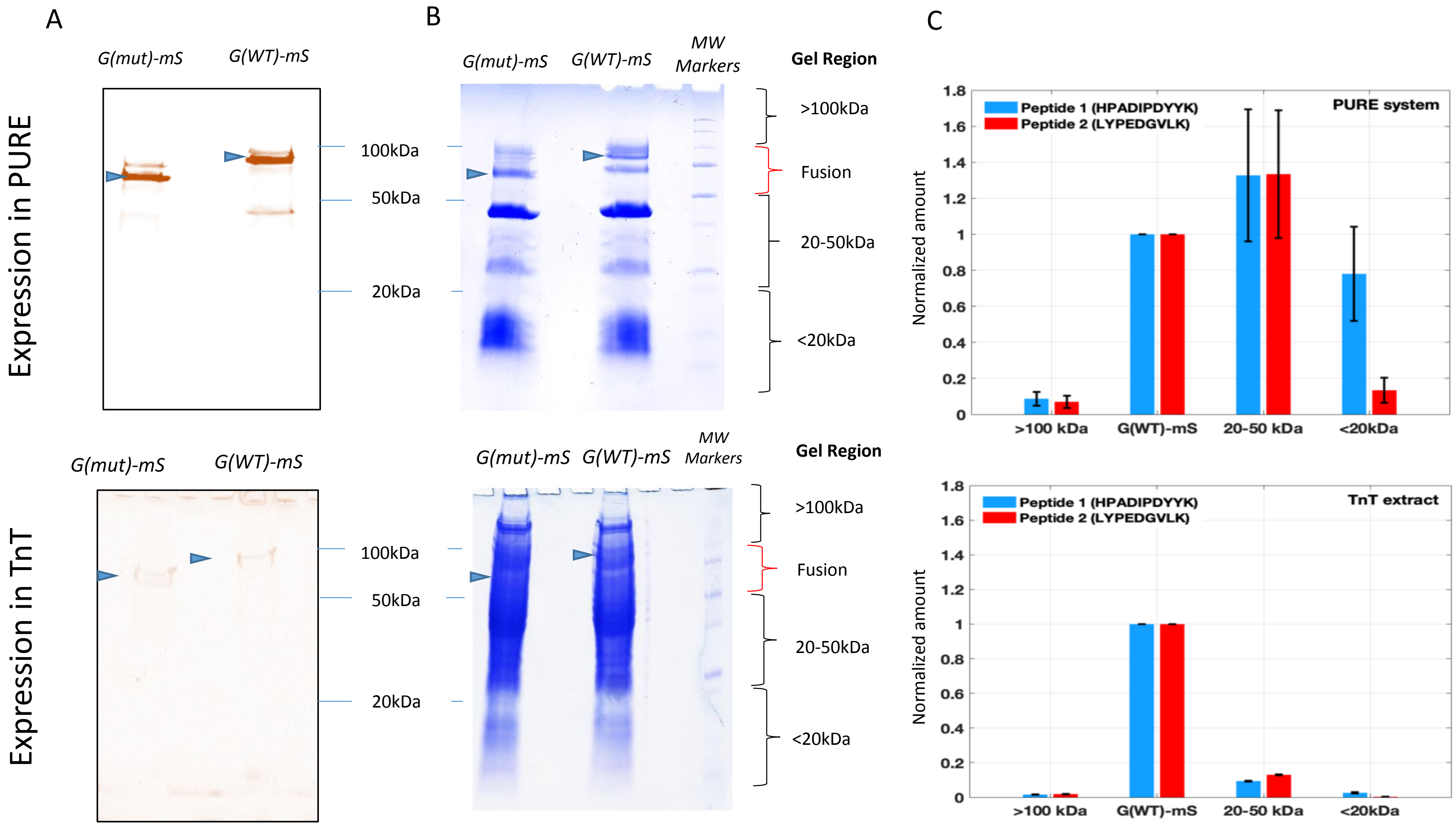


**Supplementary Figure S9.** Analysis of short products of expression. G(WT)-mS and G(mut)-mS were expressed for 4h in PURE and TnT cell free systems and aliquots of extracts separated on SDS PAGE under non-reducing conditions with no heating. **A:** fluorescence imaging of gel-separated extracts at Cy3 channel relevant to the mScarlet-I fluorescence; **B:** Corresponding SDS gels visualized by Coomassie. The position of full-length product is designated with arrow. Regions excised from the G(WT)-mS gel lane for quantification of FP peptides are indicated on the right side. **C:** Amount of mScarlet peptides 76-85 HPADIPDYYK and 151-159 LYPEDGVLK quantified using qFP-8 in G(WT)-mS at four gel regions of the G(WT)-mS gel lane marked on panel B.

A

B

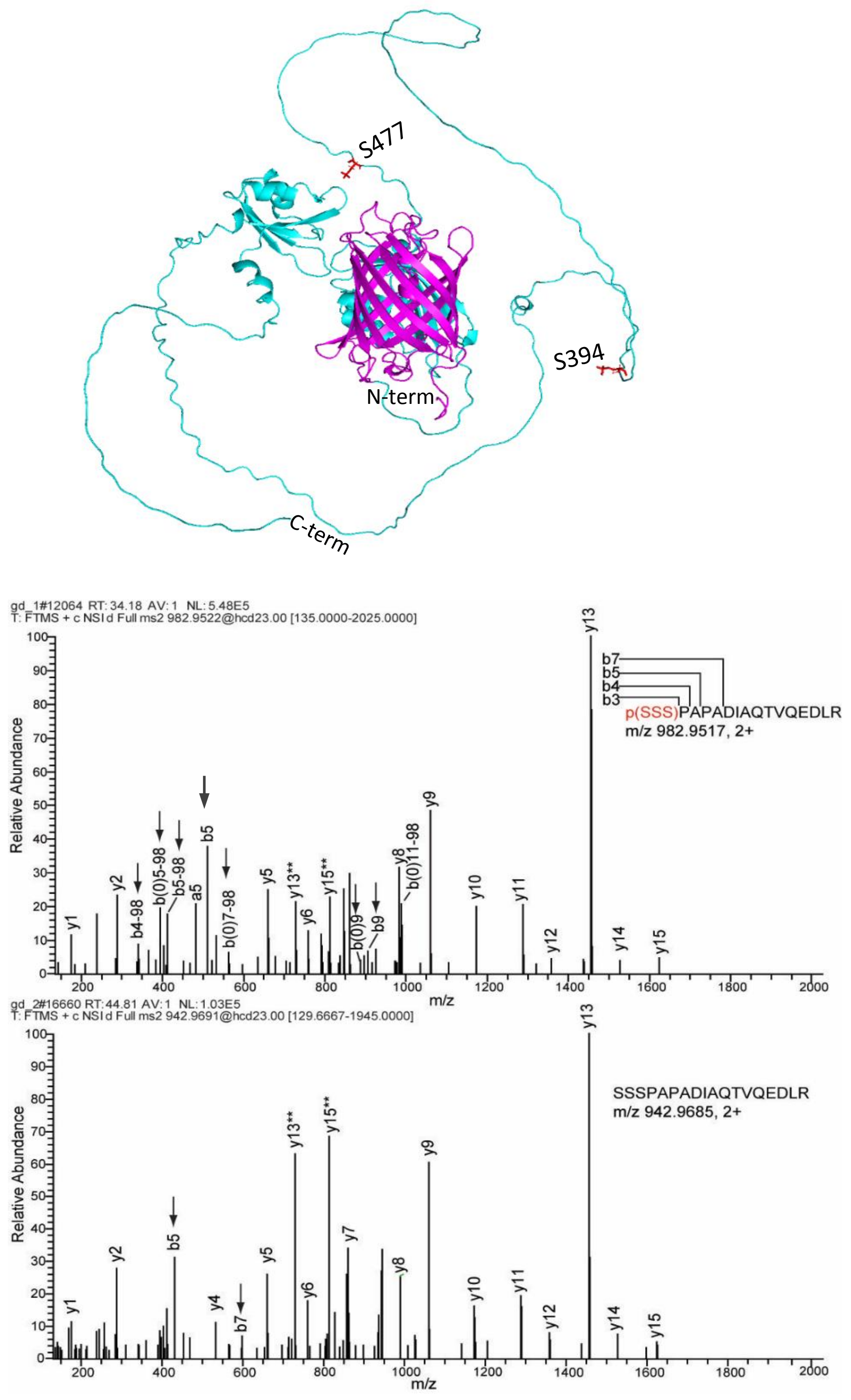


**Supplementary Figure S10**. Detection of serine phosphorylation in G(WT)-mS expressed in TnT cell-free system by mass spectrometry. **A:** 3D structure of G(WT)-mS. Position of phosphorylated serine residues S149 and S232 (393 and 477 in G(WT)-mS sequence, respectively) are shown in red. **B:** Fragmentation spectra of the mono-phosphorylated peptide (230)SSSPAPADIAQTVQEDLR(248) (upper panel) and unmodified (lower panel). Mono-phosphorylated serine cluster 230-233 is marked in red; the corresponding b-ions are designated with arrows on the spectrum.

A

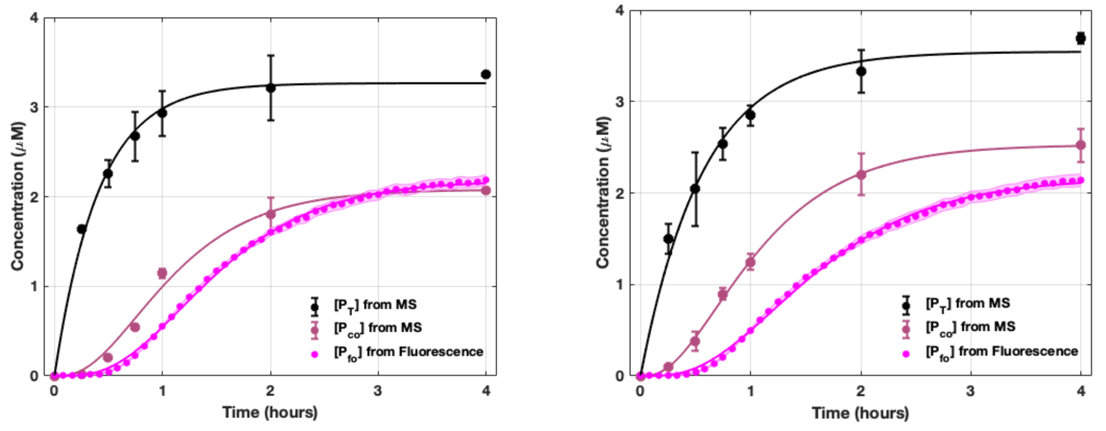


B

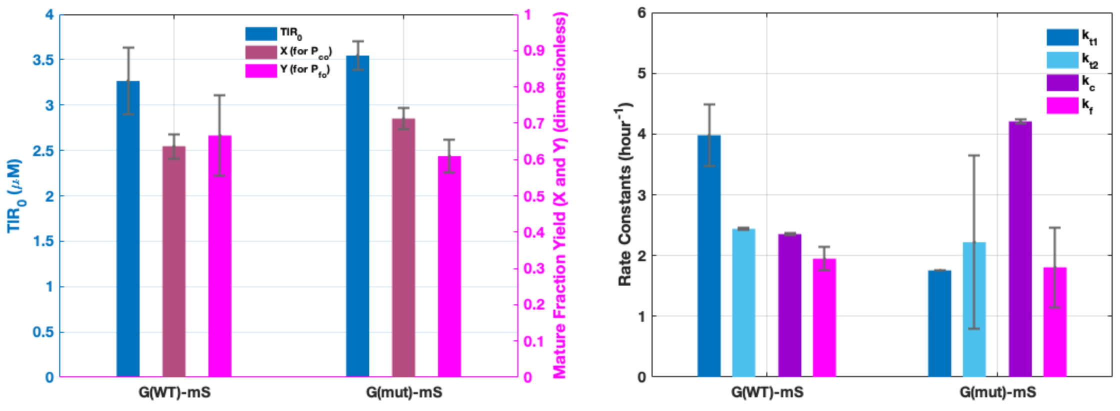


C

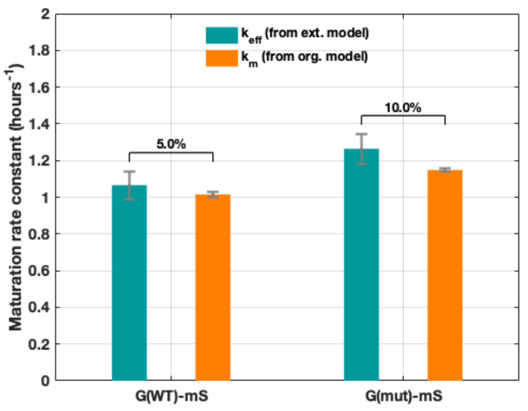


**Supplementary Figure S11.** Extended model. **A**: Fits to the extended model for G(WT)-mS (left) and G(mut)-mS (right); **B**: Fitting parameters from the extended model; **C**: Comparison between $k_{eff} = (k_c^{-1} + k_f^{-1})^{-1}$ from the extended model and $k_m$ from the original model highlighting minimal percentage differences.

A

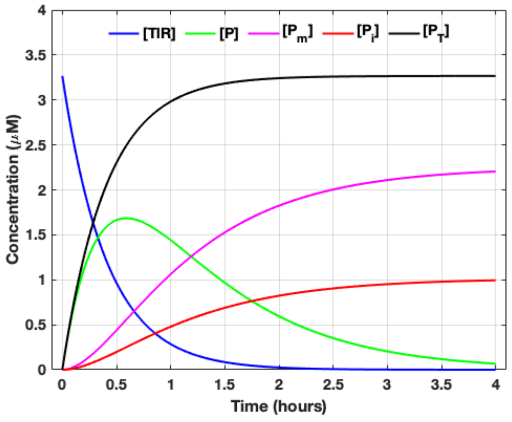


B

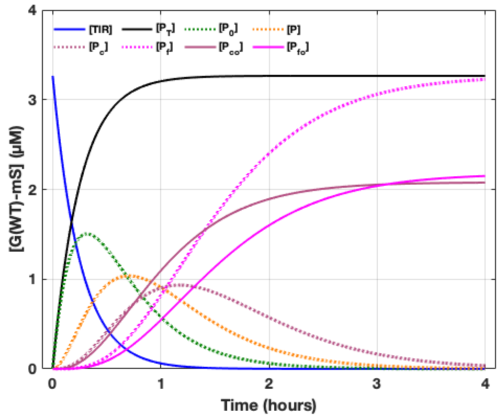


**Supplementary Figure S12**. Simulations of hidden variables using the original and extended models.

**A.** Simulations using the original model for G(WT)-mS (left) and G(mut)-mS (right) showing the kinetic traces for translation nutrient TlR, immature intermediate P, mature protein $P_m$, dark fraction that never matured, $P_i$ and the total protein $P_T$.

**B.** Simulations using the extended model for G(WT)-mS (left) and G(mut)-mS (right) showing the kinetic traces for translation nutrient TlR, precursor $P_0$, immature intermediate P, total chromophore mature fraction $P_c$, fraction of the chromophore mature fraction actually observed $P_{co}$ by mass spectrometry, total fully mature folded fraction $P_f$, fraction of the fully mature folded fraction actually observed $P_{fo}$ by fluorescence spectroscopy, and the total protein $P_T$ as observed by mass spectrometry.

**Supplementary Table S1.**

**List of Fluorescent Proteins, Self-labelling Tags and their Fusions**

| N | Name of FP or self-labelling tag | AccN[1] | FP type/ Chromophore tripeptide | MW, kDa | N of quantotypic peptides[2] | N of quantifiable FPs from the same family[3] | FP-fusion | | |
|---|---|---|---|---|---|---|---|---|---|
| | | | | | | | code | Gene name of the fused protein | MW, kDa |
| | | | | Fluorescent Proteins | | | | | |
| 1 | mKate2 | DBB08 | Basic Far-red FP MYG | 26 | 3 | *20* | #A | Spd2 | 158 |
| 2 | mScarlet-I | 6VVTK[4] | Basic RFP MYG | 26 | 4 | *3* | #E | Kif16B, MBP | 230 |
| 3 | mCherry | ZERB6 | Basic RFP MYG | 26 | 3 | *36* | #I | TJP2, MBP | 207 |
| 4 | mNeonGeen | ZRKRV | Basic yellow-green FP GYG | 26 | 3(1) | *1* | #D | KIF5B | 134 |
| 5 | mEGFP | QKFJN | Basic green FP TYG | 26 | 2(1) | *12* | #G | TJP2, MBP | 207 |
| | | | | | | | #J[6] | NMY2 | 280 |
| 6 | Dendra2 | GE6KO | Photoconvertible red-to-green FP HYG | 26 | 5 | *2* | #F | MAPT 471-756 fragment | 75 |
| 7 | Venus | YUJWJ | Basic YFP GYG | 26 | 2(1)[5] | *See mEGFP* | - | - | - |
| 8 | TagRFP | S4HC8 | Basic Orange-red FP MYG | 26 | 2[5] | *See mKate2* | - | - | - |
| 9 | dsRed-Express | DCYCK | Basic FRP QYG | 26 | 2[5] | *See mCherry* | - | - | - |
| | | | | Non-fluorescent self-labelling proteins | | | | | |
| 9 | SNAP-Tag | 3kzy | | 19 | 3 | Organic fluorophores | #H | TJP2, MBP | 200 |
| | | | | | | | #B | FUS, MBP | 119 |
| | | | | | | | #J[6] | NMY2 | 280 |
| 10 | HaloTag | 6U32_A | | 34 | 3 | Organic fluorophores | #C | CLDN2 | 62 |

1- From www.fpbase.org or PDB
2- Number of Met-containing peptides is in brackets
3- The sequence should have at least two peptides identical with proxies
4- In-house sequence is missing Val at the position 2
5- Venus, TagRFP and dsRed-Exress sequences share peptides identical with mEGFP, mKate2 and mCherry proxies respectively
6- #J is a double-tagged FP-Fusion, with SNAP-Tag on N-terminus and EGFP on C-terminus

**Supplementary Table S2**

**Peptide proxies included in qFP-8 chimeric standard protein**

| N | Origin | Peptide Sequence | m/z light form | | m/z heavy form[1] | |
|---|---|---|---|---|---|---|
| | | | +2 | +3 | +2 | +3 |
| Fluorescent proteins and self-labelling protein tags | | | | | | |
| 1 | NeonGreen | YTYEGSHIK | 549.266 | 366.514 | 552.278 | 368.521 |
| 2 | | TIISTFK | 405.242 | | 408.253 | |
| 3 | | WSYTTGNGK | 507.238 | | 510.249 | |
| 4 | | TMQFEDGASLTVNYR[2] | 866.404 | 583.270 | 871.969 | 586.607 |
| 5 | eGFP | FSVSGEGEGDATYGK | 752.334 | | 755.344 | |
| 6 | | FEGDTLVNR | 525.764 | | 530.769 | |
| 7 | | SAMPEGYVQER[2] | 633.793 | | 638.798 | |
| 8 | Dendra2 | YPEDIPDYFK | 643.801 | | 646.811 | |
| 9 | | QSFPEGYSWER | 693.310 | | 698.315 | |
| 10 | | VVQLPDAHFVDHR | | 511.604 | | 514.941 |
| 11 | | IEILGNDSDYNK | 690.836 | | 693.847 | |
| 12 | | LYEHAVAR | 479.759 | | 484.764 | |
| 13 | mScarlet-I | HPADIPDYYK | 609.793 | 406.865 | 612.804 | 408.872 |
| 14 | | QSFPEGFK | 470.232 | | 473.243 | |
| | | QSFPEGFKWER | | 470.898 | | 476.241 |
| 15 | | LYPEDGVLK | 517.282 | | 520.293 | |
| 16 | mCherry | HPADIPDYLK | 584.804 | 390.205 | 587.814 | 392.212 |
| 17 | | LSFPEGFK | 462.745 | | 465.756 | |
| 18 | | LDITSHNEDYTIVEQYER | | 742.350 | | 745.687 |
| 19 | mKate | TFINHTQGIPDFFK | 555.619 | 832.925 | 557.626 | 834.932 |
| 20 | | QSFPEGFTWER | 692.320 | | 697.326 | |
| 21 | | TLGWEASTETLYPADGGLEGR | 1112.032 | 741.690 | 1117.037 | 745.027 |
| | | *AVEGGPLPFAFDILATSFMYGSK*[3] | | | | |
| 22 | SNAP-Tag | TTLDSPLGK | 466.258 | | 469.268 | |
| 23 | | FGEVISYSHLAALAGNPAATAAVK | | 786.753 | | 788.760 |
| 24 | | VVQGDLDVGGYEGGLAVK | 888.462 | 592.644 | 891.472 | 594.651 |
| 25 | HaloTag | NIIPHVAPTHR | 418.907 | | 422.243 | |
| 26 | | LLFWGTPGVLIPPAEAAR | 954.541 | 636.696 | 959.545 | 640.033 |
| 27 | | AVDIGPGLNLLQEDNPDLIGSEIAR | 1310.185 | 873.792 | | 877.129 |
| BSA Reference peptides | | | | | | |
| 1 | BSA | DAFLGSFLYEYSR | 784.375 | | 789.380 | |
| 2 | | HLVDEPQNLIK | 653.362 | | 656.373 | |
| 3 | | LGEYGFQNALIVR | 740.401 | | 745.406 | |
| 4 | | LVNELTEFAK | 582.319 | | 585.330 | |
| 5 | | YLYEIAR | 464.250 | | 469.255 | |

[1] - metabolic labelled with $^{13}C_6{}^{15}N_4$-Arg and $^{13}C_6$-Lys

[2] – peptides comprising Methionine are not suitable for quantification if chimeric standard is digested separately and spiked into analyte prior ms-analysis

[3] – chromophore-containing peptide is observed in multiple forms depending on maturation state of the chromophore-forming triad (underlined)

**Supplementary Table S3**

**MS-based approaches for absolute quantification of proteins using peptide references and spiked protein standards**

| | Approach | Reference peptides | | Spiked protein standard | Application | Peptides compared during quantification | Ref |
|---|---|---|---|---|---|---|---|
| | | Required features | Selected from | | | | |
| A | Targeted quantification | quantotypic peptides | FP-tag of the FP-fusion | qFP-8 | Targeted | Related peptides from FP (native) and qFP-8 (labelled) | *(34)** |
| B | MBAQ | "best 3": 3 peptides with most concordant abundance | Full length sequence | FUGIS | Untargeted | Median abundance of best 3 peptides in fusion vs all FUGIS peptides | *(40)* |
| C | Top3-Hi | "top 3": 3 most abundant peptides | Full length sequence | any protein with known amount (e.g.BSA) | Untargeted | Average abundance of top 3 peptides in fusion vs reference protein | *(41)* |

*The workflow for this work was adapted from *(34)*

**Supplementary Table S4**

**Amount of FPs-fusions ##A-J quantified using peptide proxies of the qFP-8 chimeric standard**

| FP-Fusion code[1] | #A | #B | #C | #D | #E | #F | #G | #H | #I | #J[2] | |
|---|---|---|---|---|---|---|---|---|---|---|---|
| FP/tag | mKate2 | SNAP-tag | HaloTag | mNeonGreen | mScarlet-I | Dendra2 | mEGFP | Snap-tag | mCherry | Snap-tag | mEGFP |
| Calculated amount, pmol[3] | 0.620 | 0.328 | 1.179 | 0.293 | 0.104 | 0.019 | 1.526 | 1.598 | 2.086 | 0.170 | 0.198 |
| CV, % | 14 | 15 | 5 | 3 | 18 | 3 | 5 | 5 | 5 | 16 | 23 |
| CV for individual peptide proxies, % | 15 | 11 | 18 | 10 | 11 | 23 | 19 | 7 | 15 | 15 | 24 |

1 - see Supplementary Dataset S1 for details
2 – #J is a double-tagged fusion carrying EGFP and Snap-Tag on its N- and C-termini respectively.
3 – in the gel bands corresponding to the full length of the FP-fusion. Equal volume aliquots of lysate of cells expressing an FP-fusion were separated by 1D SDS PAGE prior analysis.

**Supplementary Table S5**

**Examples of FP amounts in stably transfected cells quantified using qFP-8 standard**

| N | FP | ID (FPbase) | Expressed in (cell type) | FP concentration quantified using qFP-8 | FP amount per cell[2] | Cell line proprietor[3] |
|---|---|---|---|---|---|---|
| 1 | mEGFP | QKFJN | HeLa[1] | 81.7 µM | 147 amol | *A* |
| 2 | mEGFP | QKFJN | HeLa[1] | 212.1µM | 382 amol | *A* |
| 3 | Venus | YUJWJ | HCT116 | 3.9µM | 2 amol | *B* |
| 4 | TagRFP | S4HC8 | HCT116 | 17.7µM | 9 amol | *B* |
| 5 | mScarlet-I | 6VVTK | E.coli | 5.2mM | 5.7 amol | *C* |
| 6 | mKate2 | DBB08 | E.coli | 0.6mM | 0.6 amol | *C* |
| 7 | mCherry | ZERB6 | E.coli | 0.8mM | 0.9 amol | *C* |

[1] - two independently prepared cell lanes
[2] - recalculated from FP concentration
[3] – A: Technology Development Studio; B: Dr. M.Sarov (Genome Engineering Facility); C: Protein Biochemistry Facility (all at MPI CBG, Dresden)

**Supplementary Table S6**

**Abundance of chromophore-containing peptides in red FP mScarlet and dsRed-express**

| **FP** | **Chromophore containing peptide**[3] | **Chromophore maturation form**[4] | **m/z** | **Peptide abundance (XIC)** |
|---|---|---|---|---|
| mScarlet-I[1] | GGPLPFSWDILSPQ*MYG*SR | linear | 757.7017 | 4.99e8 |
| | | intermediate | 751.0268 | 8.21e9 |
| | | mature | 750.3548 | 2.23e10 |
| mScarlet mutant M190[2] | GGPLPFSWDILSPQ*MYG*SR | linear | 757.7017 | 2.05e10 |
| | | intermediate | 751.0268 | 1.48e8 |
| | | mature | 750.3548 | 5.38e8 |
| dsRED-express[1] | GGPLPFAWDILSPQF*QYG*SK | linear | 736.7090 | 2.27e10 |
| | | intermediate | 730.0350 | 9.73e9 |
| | | mature | 729.3620 | 1.58e7 |

[1] - as in Supplementary Table S1
[2] – non-fluorescent mutant described in *(49)*
[3] – chromophore tripeptide is shown in Italic, cyclic forms are underlined
[4] –as described in *(50)*

**Supplementary table S7**

**Phosphorylation status of the G3BP1 peptide SSSPAPADIAQTVQEDLR detected by mass spectrometry in G(WT)-mS and G(mut)-mS expressed in TnT and PURE cell free expression systems**

| FP-fusion | Peptide SSSPAPADIAQTVQEDLR, as detected by mass spectrometry[1] | |
|---|---|---|
| | With phosphorylated serine[2] | Unmodified |
| **Expressed in insect cell line SF9 (control)** | | |
| **G(mut)-mS** | gd_1.12064.12064.2.dta | gd_1.17063.17063.2.dta |
| **Expressed in PURExpress cell-free system** | | |
| **G(WT)-mS** | n/d | a_1pmolbsa_0_1fp21_pw_inf_1_dda_50ul_1_.25193.25193.2.dta |
| **G(mut)-mS** | n/d | e_inf_50ul_0_1_fp21_1pmolbsa_dda.16579.16579.2.dta |
| **Expressed in TnT®T7 cell-free system** | | |
| **G(WT)-mS** | a_1pmolbsa_0_1fp21_i_inf_1_dda_50ul_1.17350.17350.2.dta | a_1pmolbsa_0_1fp21_i_inf_1_dda_50ul_1.16627.16627.2.dta |
| **G(mut)-mS** | a_inf_50ul_02_20220418083554.23082.23082.2.dta | a_inf_50ul_02_20220418083554.22049.22049.2.dta |

[1] ID of the best MS2 spectra; n/d – not detected
[2] the peptide was detected only monophosphorylated by one of Serines
